# Reconstitution of BNIP3/NIX-mediated autophagy reveals two pathways and hierarchical flexibility of the initiation machinery

**DOI:** 10.1101/2024.08.28.609967

**Authors:** Elias Adriaenssens, Stefan Schaar, Annan S.I. Cook, Jan F. M. Stuke, Justyna Sawa-Makarska, Thanh Ngoc Nguyen, Xuefeng Ren, Martina Schuschnig, Julia Romanov, Grace Khuu, Michael Lazarou, Gerhard Hummer, James H. Hurley, Sascha Martens

## Abstract

Selective autophagy is a lysosomal degradation pathway that is critical for maintaining cellular homeostasis by disposing of harmful cellular material. While the mechanisms by which soluble cargo receptors recruit the autophagy machinery are becoming increasingly clear, the principles governing how organelle-localized transmembrane cargo receptors initiate selective autophagy remain poorly understood. Here, we demonstrate that transmembrane cargo receptors can initiate autophagosome biogenesis not only by recruiting the upstream FIP200/ULK1 complex but also via a WIPI-ATG13 complex. This latter pathway is employed by the BNIP3/NIX receptors to trigger mitophagy. Additionally, other transmembrane mitophagy receptors, including FUNDC1 and BCL2L13, exclusively use the FIP200/ULK1 complex, while FKBP8 and the ER-phagy receptor TEX264 are capable of utilizing both pathways to initiate autophagy. Our study defines the molecular rules for initiation by transmembrane cargo receptors, revealing remarkable flexibility in the assembly and activation of the autophagy machinery, with significant implications for therapeutic interventions.

## INTRODUCTION

Selective autophagy is a critical process for maintaining cellular homeostasis. It ensures the degradation of damaged or superfluous components, such as organelles, protein aggregates, and cytosol-invading pathogens within lysosomes. This targeted removal is orchestrated by specialized proteins called cargo receptors, which link the cargo material to the autophagy machinery ^1^.

A crucial distinction exists between soluble and transmembrane cargo receptors. Soluble cargo receptors, such as SQSTM1/p62, NBR1, TAX1BP1, NDP52 and OPTN are dispersed across the cytosol and dynamically recruited to the cargo material upon its ubiquitination. Once recruited, these receptors attract components of the upstream machinery to induce autophagosome biogenesis in proximity to the cargo ^2^. Canonically, the cargo receptors recruit the FIP200 proteins, a subunit of the upstream ULK1 kinase ^3-6^. Recently, it was shown that OPTN recruits the TBK1 kinase and ATG9A, which are also upstream factors in selective autophagy ^7,8^.

In contrast, transmembrane cargo receptors reside on the various organelles and display greater diversity in terms of number and structure. They can be single-pass, multi-pass, or tail-anchored proteins. Currently, over 15 different membrane-embedded cargo receptors are known, and the list is expanding rapidly. Notably, for mitochondria these include BNIP3^9-11^, NIX^12-15^ (also known as BNIP3L), FKBP8^16^, PHB2^17^, NLRX1^18^, MCL-1^19^, FUNDC1^20^, and BCL2L13^21^; for the endoplasmic reticulum (ER), ATL3^22^, CCPG1^23^, FAM134A^24^, FAM134B^25^, FAM134C^24,26^, Sec62^27^, RTN3^28^, and TEX264^29,30^; for the Golgi apparatus, YIPF3 and YIPF4^31^; and for peroxisomes, NIX and BNIP3^32^.

While the mechanisms by which soluble cargo receptors initiate autophagy have been elucidated, the process by which transmembrane cargo receptors recruit the autophagy machinery remains less clear. Given the large number of transmembrane cargo receptors spread across the different organelles, understanding their mode of action is crucial for a comprehensive understanding of selective autophagy.

In this study, we investigated the mechanism of autophagosome biogenesis by transmembrane cargo receptors. We found that, in contrast to soluble cargo receptors, transmembrane cargo receptors can initiate autophagosome biogenesis through two distinct pathways: one by recruiting the upstream FIP200/ULK1 complex, and another by recruiting a WIPI-ATG13 complex. Our results reveal an unexpected flexibility among selective autophagy pathways and show that the general principles of soluble cargo receptors do not universally apply to all transmembrane cargo receptors.

## RESULTS

### NIX and BNIP3 are unable to bind FIP200

Human cells express numerous transmembrane cargo receptors, typically several for each organelle ^33^. To understand how these receptors recruit the autophagy machinery, we focused on mitochondria, where several single-pass and multi-pass transmembrane cargo receptors have been identified (**Fig. 1a**) ^34^. Unlike other organelles such as the endoplasmic reticulum (ER), mitochondria can be targeted for selective autophagy using chemical agents like deferiprone (DFP), which induce mitophagy via individual receptors ^10^.

**Figure 1.**
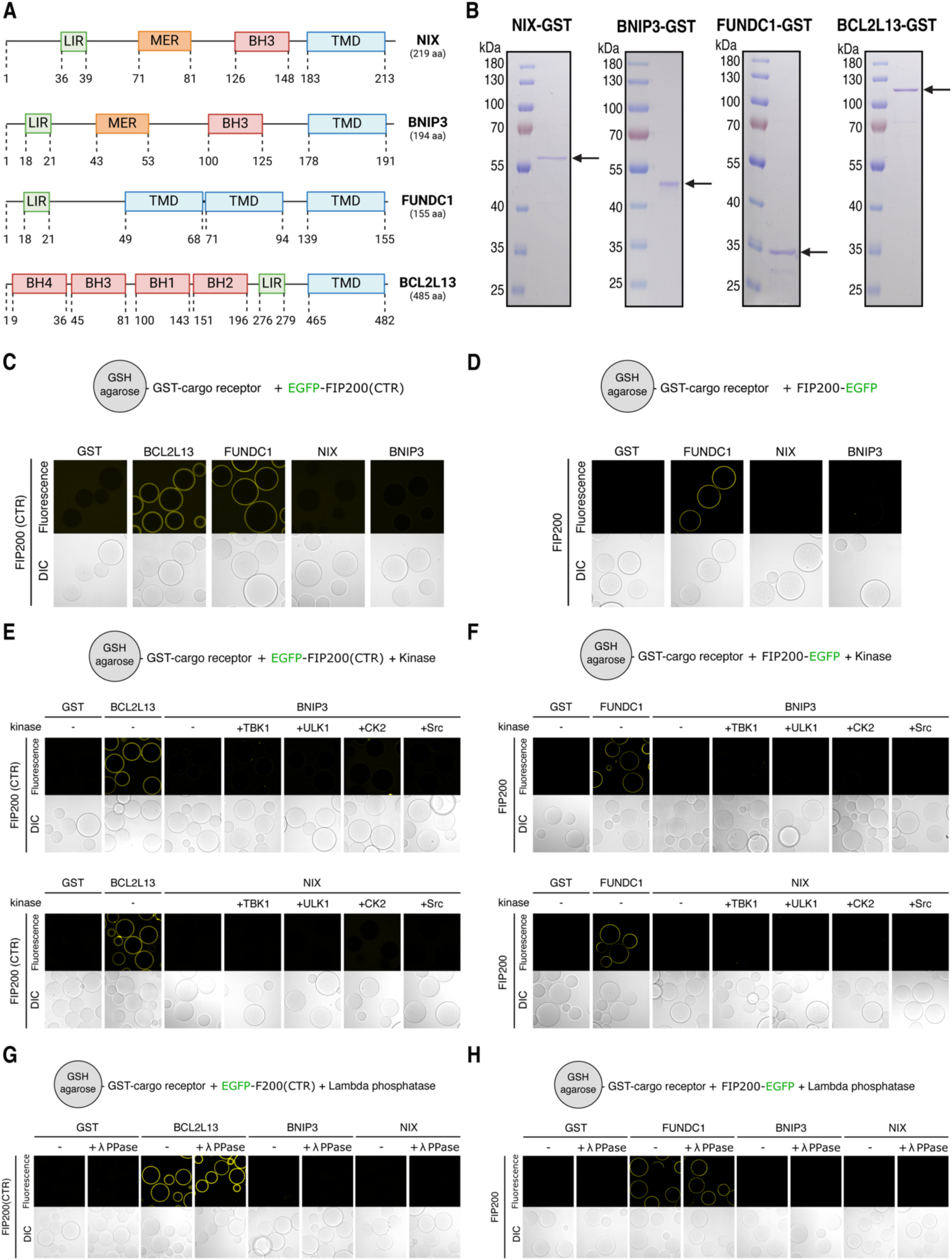
NIX and BNIP3 are unable to bind FIP200 *in vitro*. (**A**) Schematic of the domain structures of NIX, BNIP3, FUNDC1, and BCL2L13. LC3-interacting motif (LIR), Minimal essential region (MER), Bcl-2 Homology domain (BH), transmembrane domain (TMD). (**B**) Representative SDS-PAGE gels of NIX(1-182aa)-GST, BNIP3(1-158aa)-GST, FUNDC1(1-50aa)-GST, and BCL2L13(1-465aa)-GST. Arrows indicate the predicted molecular weight. (**C-H**) Microscopy-based bead assay of agarose beads coated with the indicated GST-tagged cargo receptors and incubated with (C) GFP-tagged FIP200-CTR (residues 1429-1591), (D) GFP-tagged full-length FIP200, (E) FIP200-CTR and kinases TBK1, MBP-ULK1, CK2, or Src (Y530F; constitutively active mutant), (F) full-length FIP200 and kinases TBK1, MBP-ULK1, CK2, or Src (Y530F; constitutively active mutant), (G) FIP200-CTR and Lambda Protein Phosphatase, (H) full-length FIP200 and Lambda Protein Phosphatase. Samples were analyzed by confocal imaging and one of three representative experiments is shown.

To investigate the recruitment process of the autophagy machinery by transmembrane mitophagy receptors, we reconstituted the initiation of autophagosome biogenesis using purified components. We purified the soluble, cytosol-exposed domains of BNIP3, NIX, FUNDC1, and BCL2L13 (**Fig. 1b**), substituting the transmembrane domains with GFP- or GST-moieties to study the mitophagy receptors in either a monomeric or dimeric state. For instance, for NIX and BNIP3, the activated state is thought to be a dimer^35^, while for FUNDC1 and BCL2L13, this is yet to be elucidated.

To confirm that our purified mitophagy receptors are active, we tested their ability to bind LC3 and GABARAP proteins using a microscopy-based bead assay (**Fig. S1a**). Similar to soluble cargo receptors, GABARAP proteins were bound more readily, whereas LC3 proteins showed varying degrees of binding depending on the receptor. Specificity was verified by mutating the LC3-interacting (LIR) motifs, resulting in the loss of binding for NIX, BNIP3, and FUNDC1 (**Fig. S1b**). For BCL2L13, multiple functional LIR motifs were observed (**Fig. S1c-d**), similar to how the yeast Atg19 interacts with Atg8^36^.

Having confirmed that our purified mitophagy receptors are active, we next sought to determine how they recruit the remaining autophagy machinery. Soluble cargo receptors, such as SQSTM1/p62, initiate autophagosome biogenesis by binding to FIP200 through a FIP200-interacting (FIR) motif that docks into a conserved groove of the C-terminal FIP200 Claw domain ^4^. We therefore tested whether the transmembrane mitophagy receptors could also bind the C-terminal region of FIP200, which encompasses the Claw domain and a portion of the coiled-coil domain (residues 1429-1591). Using microscopy-based bead assays, we observed that FUNDC1 and BCL2L13, but not BNIP3 or NIX, directly bind to the C-terminal FIP200 domain (**Fig. 1c**). Moreover, mutating the LIR/FIR motifs of FUNDC1 or BCL2L13 abrogated this interaction (**Fig. S1e**), confirming the specificity of the interaction.

Not all soluble cargo receptors bind FIP200 in the Claw domain. For instance, NDP52 binds the coiled-coil region just upstream of the C-terminal region ^5,6,37^. Therefore, we tested whether BNIP3 and NIX could bind to full-length FIP200 (**Fig. 1d**). However, we were unable to detect a direct interaction between BNIP3/NIX and FIP200.

Next, we asked if BNIP3/NIX require activation by a kinase, such as TBK1, which is known to phosphorylate soluble cargo receptors and cargo co-receptors to enhance their LC3-binding capacities ^38,39^. In particular, we tested four candidate kinases: TBK1, ULK1, Src, and casein kinase 2 (CK2). TBK1 and ULK1 have been previously shown to play essential roles in selective autophagy pathways involving soluble cargo receptors ^7,39,40^, while Src and CK2 have been associated with hypoxia-induced mitophagy ^41,42^. We therefore purified TBK1, MBP-ULK1, Src, and CK2 (**Fig. S2**) and performed microscopy-based protein-protein interaction assays between BNIP3/NIX and either full-length or the C-terminal region of FIP200. Our results show that, while the positive controls FUNDC1 and BCL2L13 were able to bind FIP200, the addition of the kinases and ATP/MgCl_2_ did not facilitate an interaction between BNIP3/NIX and FIP200 (**Fig. 1e-f**).

We hypothesized that purified BNIP3/NIX might already be pre-phosphorylated, which could potentially silence their activity towards FIP200. To test this, we performed an microscopy-based bead assay in the presence of Lambda Protein Phosphatase. While BCL2L13 and FUNDC1 would readily bind to FIP200 under these conditions, we could not observe a direct binding of NIX to FIP200 (**Fig. 1g-h**).

Some soluble cargo receptors, like Optineurin, have been shown to recruit other upstream autophagy machinery ^7,43,44^. Therefore, we tested whether BNIP3/NIX could initiate autophagy not by recruiting FIP200, but through the recruitment of TBK1, PI3KC3-C1 complex, or ATG9A-vesicles. However, in vitro binding assays with purified TBK1, PI3KC3-C1 complex, or ATG9A-vesicles did not reveal any direct interactions with BNIP3/NIX (**Fig. S3**).

In summary, while our findings confirm that the mitophagy receptors FUNDC1 and BCL2L13 directly bind to FIP200, similar to the mechanism by which most soluble cargo receptors initiate autophagosome biogenesis, we could not detect any direct binding between the mitophagy receptors BNIP3/NIX and FIP200 or other upstream autophagy machinery components.

### NIX and BNIP3 initiate mitophagy by recruiting WIPI proteins

Since we were unable to establish a direct interaction between BNIP3/NIX and any of the upstream autophagy machinery, we explored whether BNIP3/NIX utilize an alternative mechanism for recruiting the autophagy machinery upon mitophagy induction. Recent studies have shown that NIX interacts with WIPI2 ^45^, a downstream factor in the autophagy cascade, and PPTC7 ^46,47^, a mitochondrial phosphatase that accumulates on the mitochondrial surface upon iron-depletion by DFP treatment ^48-50^.

To identify other potential interactors of NIX and BNIP3 that could link these receptors to the upstream autophagy machinery, we coated GSH-beads with GST-tagged NIX, GST-tagged BNIP3, or GST alone as a control and incubated them with HeLa cell lysates. Mass spectrometry analysis revealed that PPTC7 was the strongest binder for NIX and one of the strongest binders for BNIP3 (**Fig. 2a**). Additionally, we detected WIPI2 among the top binders for NIX and WIPI3 as a top binder for BNIP3. The interaction between BNIP3 and WIPI3 has not been reported before, but given the concomitant interaction between NIX and WIPI2 and the absence of other upstream autophagy components in our dataset, it suggests a potentially important role for WIPI2 and WIPI3 in BNIP3/NIX-mediated mitophagy.

**Figure 2.**
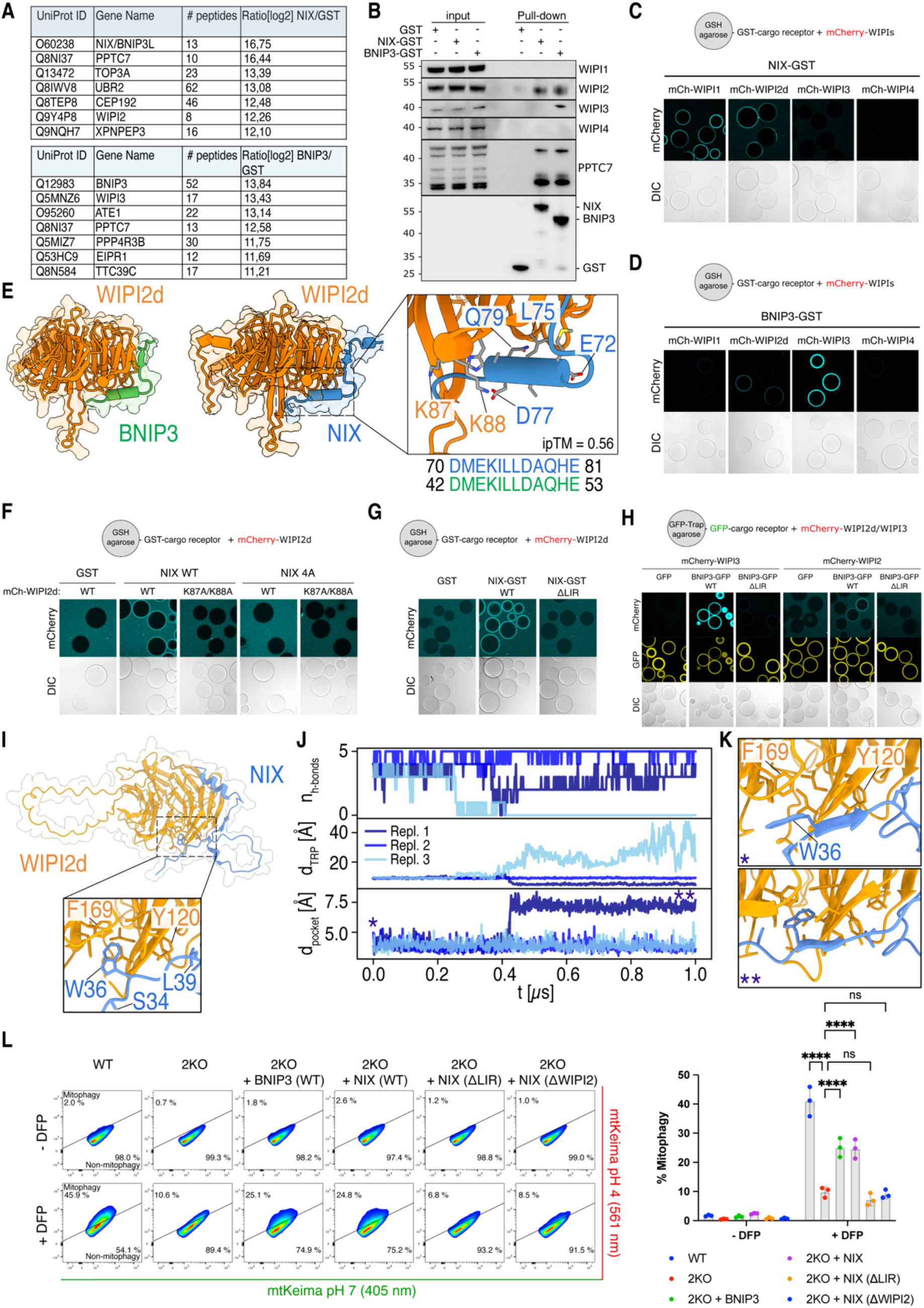
NIX and BNIP3 initiate mitophagy through WIPI2 and WIPI3. **(A)** Identification of interactors of NIX(1-182)-GST and BNIP3(1-158)-GST in comparison to a GST control by pull-down from HeLa cell lysates and label-free quantitative mass spectrometry analysis. Tables represent the top hits for NIX (upper) and BNIP3 (lower). **(B)** Validation of mass spectrometry data by analyzing the pull-downs with SDS-PAGE and western blot analysis. **(C-D)** Microscopy-based bead assay of agarose beads coated with cargo receptors NIX(1-182)-GST or BNIP3(1-158)-GST and incubated with mCherry-tagged WIPI1, WIPI2d, WIPI3, or WIPI4. **(E)** AlphaFold-2 predicted structure of NIX or BNIP3 and WIPI2d. Note that the indicated residue numbers for WIPI2, correspond to their residue number in the WIPI2d sequence (which match residue numbers K105 and K106 in WIPI2b). The conservation of the interaction interface, between NIX and BNIP3, is displayed below the zoom out. **(F-G)** As in (C) but with NIX wild-type (WT), E72A/L75A/D77A/E81A mutant (4A), or W36A/L39A (ΔLIR) and incubated with WIPI2d wild-type (WT) or K87A/K88A mutant. **(H)** As in (C) but with BNIP3 wild-type (WT) or W18A/L21A mutant (ΔLIR) and incubated with mCherry-tagged WIPI3 or WIPI2d. **(I)** AF Multimer predicted complex structure of WIPI2d and NIX (residues 30-82). Zoom highlights the interaction between the LIR of NIX and WIPI2d. The C-terminal intrinsically disordered region of WIPI2d is omitted for visual clarity. **(J)** Number of backbone h-bonds nh-bonds between the LIR of NIX and WIPI2d, insertion depth dTRP of NIX W36, and minimum heavy atom distance dpocket between WIPI2d F169 and I133 from three 1 μs MD simulations. **(K)** Representative snapshots of W36 interacting at the surface of WIPI2d (top) and inserted into an initially closed pocket (bottom). The symbols in the lower left corner indicate the point in the trajectory in **(J)** where the respective snapshots were extracted from. **(L)** Mitophagy flux was measured by flow cytometry of wild-type (WT) or NIX/BNIP3 double knockout (2KO) HeLa cells, rescued where indicated with V5-BNIP3, V5-NIX, V5-NIX E72A/L75A/D77A/E81A mutant (4A mutant; ΔWIPI2), or V5-NIX W36A/L39A mutant (ΔLIR), left untreated or treated with DFP for 24 h. Representative FACS plots are shown from one of three replicates (I). The percentage of non-induced cells (lower right) versus mitophagy-induced cells (upper left) is indicated. Two-way ANOVA with Tukey’s multiple comparisons test in (I). \*\*\*\**P*<0.0001. ns, not significant.

The importance of the direct recruitment of WIPI proteins by cargo receptors BNIP3/NIX in mitophagy, typically recruited only after the upstream ULK1- and PI3KC3-C1 complexes have been loaded onto ATG9-vesicle seeds, is unclear. However, given our failure to identify any upstream regulatory factors of the autophagy machinery, we decided to investigate the interaction with WIPI proteins in more detail.

First, to confirm the mass spectrometry results, we incubated GST, NIX-GST, and BNIP3-GST with HeLa cell lysate and immunoblotted for different WIPI proteins. Indeed, NIX and BNIP3 bound WIPI2, while BNIP3 also pulled down WIPI3 (**Fig. 2b**). To test whether NIX and BNIP3 bind WIPI2 and WIPI3 directly, we incubated purified WIPI1-4 with NIX- or BNIP3-coated agarose beads. This revealed that NIX binds to WIPI2, but not WIPI3 under these conditions (**Fig. 2c**), consistent with our mass spectrometry dataset. We also observed that NIX can bind to WIPI1, which is structurally related to WIPI2. For BNIP3, we detected an interaction with WIPI2 and a much stronger binding to WIPI3 (**Fig. 2d**).

Using AlphaFold-2 (AF2) Multimer, we modeled the NIX-WIPI2 and BNIP3-WIPI2 complexes. These predictions suggested that a short amino acid stretch, conserved between NIX and BNIP3, interacts with WIPI2 (**Fig. 2e and Fig. S4**). To test this model, we introduced point mutations in the predicted binding interfaces and observed a complete loss of binding between NIX and WIPI2 (**Fig. 2f**). Interestingly, we also observed a role for the LIR motif of NIX, as mutating the LIR motif abrogated the interaction (**Fig. 2g**). Consistently, mutating the LIR motif of BNIP3 abrogated the BNIP3-WIPI2 and BNIP3-WIPI3 interactions (**Fig. 2h**).

We then employed further AF2 Multimer modelling and molecular dynamics (MD) simulations to model where the LIR motif of NIX may engage with WIPI2d. This revealed an interaction of the LIR at the surface of WIPI2d (**Fig. 2i**), that was—with some minor structural rearrangements—stable for several hundred nanoseconds in our MD simulations (**Fig. 2j**). Interestingly, we observed the opening of a cryptic pocket in WIPI2d, which accommodated the Trp residue of the LIR of NIX (**Fig. 2k**), suggesting a possible mechanism for the LIR-WIPI2d interaction. When we mutated the LIR motif, it was no longer predicted to bind the cryptic pocket in WIPI2d (**Fig. S4j**), consistent with our biochemical data. Combined, our biochemical and MD data reveal that BNIP3/NIX bind WIPI2 using two motifs.

To assess the importance of the BNIP3/NIX-WIPI interactions in cells, we generated BNIP3/NIX double knockout HeLa cells and confirmed they are deficient in DFP-induced mitophagy (**Fig. S5a-b**). We then rescued the double knockout cells with wild-type BNIP3, wild-type NIX, WIPI2-binding-deficient or LIR-deficient NIX mutants. This revealed that the BNIP3/NIX-WIPI interactions are essential for DFP-induced mitophagy, as both WIPI2-binding-deficient and LIR-deficient NIX were unable to rescue the knockouts (**Fig. 2l**).

Our data thus reveal that NIX and BNIP3 use two binding motifs to interact with WIPI2 and/or WIPI3, respectively. Furthermore, we demonstrate that these interactions are essential for BNIP3/NIX-mediated mitophagy.

### Mitochondrial localization of WIPI1, WIPI2, and WIPI3 is sufficient to initiate autophagosome biogenesis

To investigate if the BNIP3/NIX-mediated recruitment of WIPI proteins—typically considered downstream factors—is sufficient for mitophagy initiation, we artificially tethered WIPI proteins to the mitochondrial surface. Using the FK506 binding protein (FKBP) and FKBP-rapamycin binding (FRB) system, which facilitates chemical-induced dimerization, we generated FKBP-GFP-WIPI fusion proteins for WIPI1, WIPI2, WIPI3, and WIPI4, and expressed those constructs via stable lentiviral transduction in HeLa cells expressing Fis1-FRB (**Fig. 3a**). By co-expressing the mitochondrially targeted monomeric Keima (mt-mKeima) probe, we assessed mitochondrial turnover to determine if the recruitment of WIPI proteins to the mitochondrial surface could initiate mitophagy. The addition of rapalog resulted in a strong induction of mitophagy for WIPI1, WIPI2, and WIPI3, but not WIPI4 (**Fig. 3b**).

**Figure 3.**
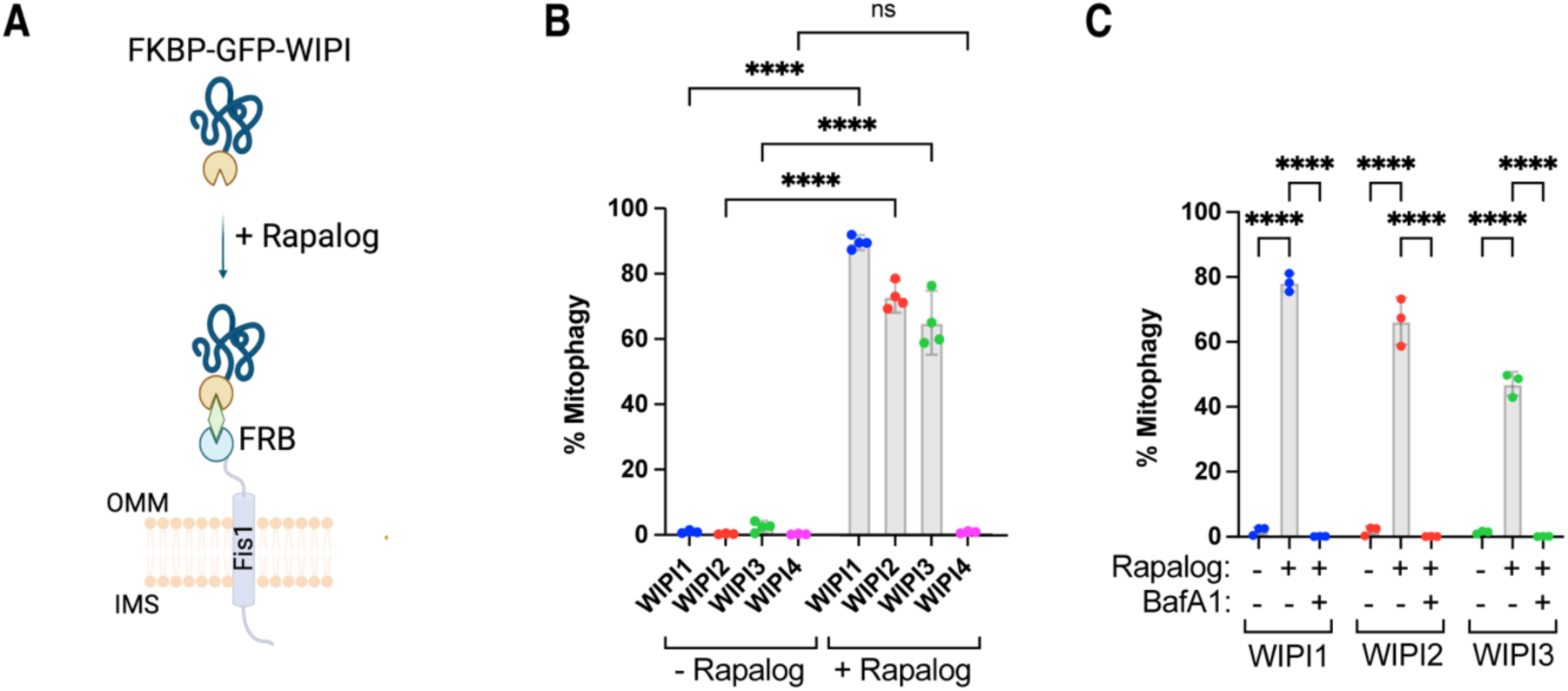
Mitochondrial localization of WIPI1, WIPI2, WIPI3 can initiate autophagosome biogenesis. **(A)** Diagram of the experimental set-up and the effect of rapalog treatment, resulting in the tethering of WIPI proteins to the outer mitochondrial membrane. IMS: intermembrane space, OMM: outer mitochondrial membrane. **(B)** Mitophagy flux was measured by flow cytometry in wild-type HeLa cells expressing Fis1-FRB, FKBP-GFP-WIPI1/2/3/4, and mt-mKeima, not induced or induced for 24 h by rapalog treatment. **(C)** As in (B) but with or without the addition of autophagy inhibitor Bafilomycin A1 (BafA1). Two-way ANOVA with Šídák’s multiple comparisons test in (B) and Dunnett’s multiple comparisons test in (C). **P<0.005, ***P<0.001, ****P<0.0001. ns, not significant.

To confirm that this mitochondrial turnover was mediated by autophagy, we repeated the experiment for WIPI1, WIPI2, and WIPI3 in the presence of Bafilomycin A1, which blocks autophagosome degradation (**Fig. 3c**). Bafilomycin A1 treatment completely inhibited mitochondrial turnover, confirming that tethering WIPI1, WIPI2, WIPI3 to the mitochondrial surface is sufficient to induce mitophagy.

This finding was unexpected, as WIPI proteins are generally considered downstream factors in autophagosome biogenesis, recruited to the expanding phagophore only after PI3KC3-C1 phosphorylation of phosphatidylinositol. However, our data suggest that the recruitment of these downstream factors to mitochondria is sufficient to initiate autophagosome formation.

### Mitophagy initiation through WIPI proteins requires the upstream ULK1 complex

To understand the mechanism by which WIPI proteins can initiate autophagosome biogenesis, we first tested whether the upstream autophagy complexes, such as the ULK1 complex (composed of FIP200, ATG13, ATG101, and the ULK1 kinase), are still required. We tethered WIPI2 to the mitochondrial surface using the rapalog system and immunostained the cells for ATG13, showing that ATG13 is still recruited to the mitochondrial surface upon artificial tethering of WIPI2 (**Fig. 4a**). To determine if this recruitment coincided with ULK1 complex activation, we performed immunoblotting for phosphorylated ATG13, showing that ATG13 becomes phosphorylated when WIPI2 is recruited to the mitochondrial surface (**Fig. 4b**).

**Figure 4.**
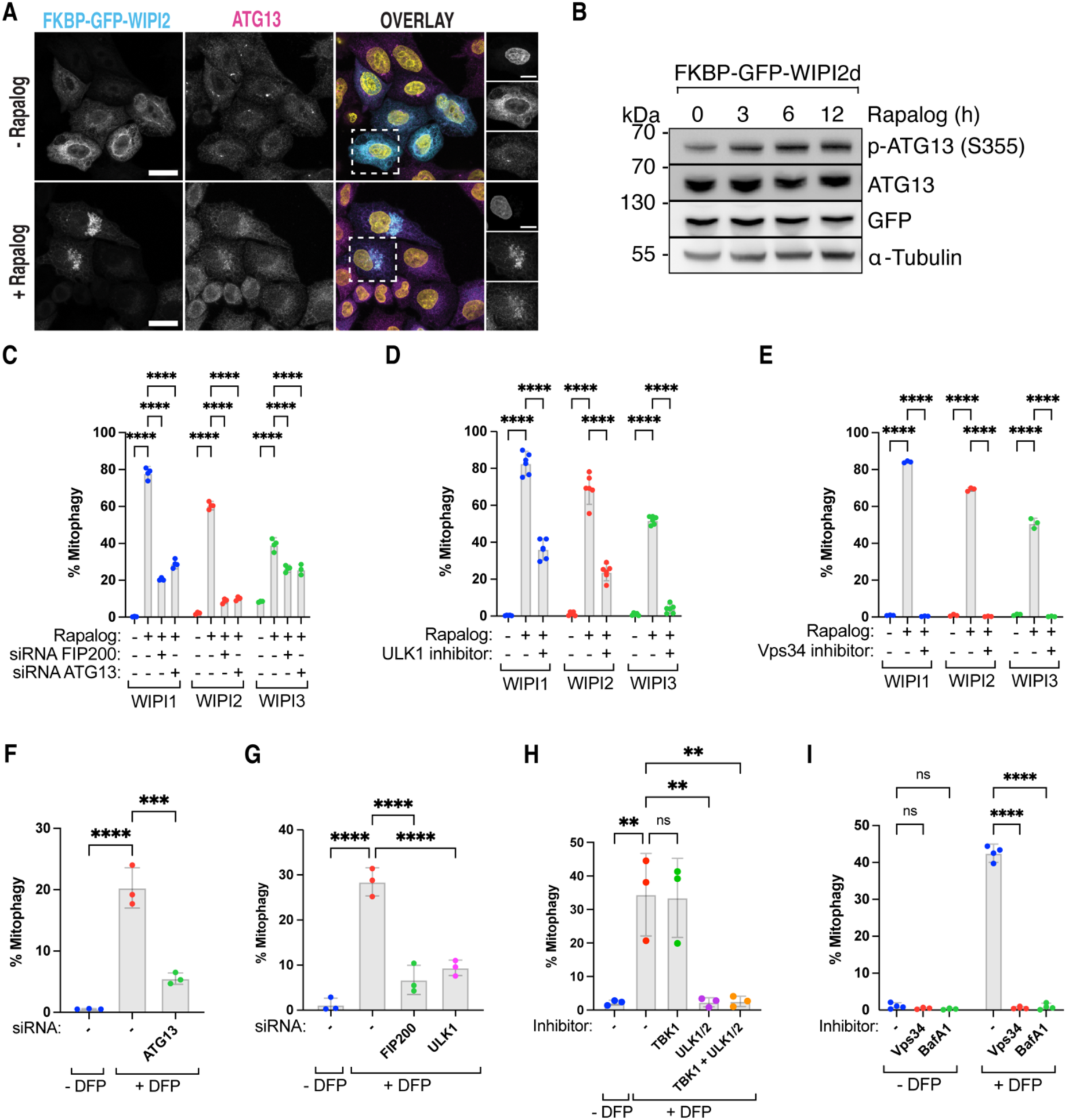
ULK1-complex and PI3KC3-C1 complex are required downstream of WIPI-driven autophagosome biogenesis. (**A**) Representative maximum intensity projection images of wild-type (WT) HeLa cells stably expressing Fis1-FRB and FKBP-GFP-WIPI2. Cells were left untreated (- Rapalog) or treated with Rapalog for 16 h (+ Rapalog) and immunostained for anti-ATG13. Scale bars: overviews, 20 µm; insets: 10 µm. (**B**) Immunoblotting for phosphorylated ATG13 in HeLa cells overexpressing Fis1-FRB and FKBP-EGFP-WIPI2d, treated with Rapalog for the indicated time. (**C**) Mitophagy flux was measured by flow cytometry in wild-type HeLa cells transfected with siRNAs targeting FIP200 or ATG13, and expressing Fis1-FRB, FKBP-GFP-WIPI1/2/3, and mt-mKeima, not induced or induced for 24 h by rapalog treatment. (**D-E**) As in (C) but with or without the addition of (D) the ULK1/2 inhibitor MRT68921, or (E) the Vps34-inhibitor VPS34-IN1. (**F-G**) Wild-type HeLa cells expressing mt-mKeima and transfected with siRNAs targeting ATG13, FIP200, or ULK1, and treated with DFP for 24 h. (**H-I**) As in (F) but with the kinase inhibitors GSK8612 for TBK1, MRT68921 for ULK1/2, VPS34-IN1 for Vps34, or Bafilomycin A1 (BafA1). Two-way ANOVA with Dunnett’s multiple comparisons test in (C-E, I) or One-way ANOVA with Dunnett’s multiple comparisons test (F-H). **P<0.005, ***P<0.001, ****P<0.0001. ns, not significant.

To assess whether upstream autophagy complexes are also required for WIPI-induced mitophagy initiation, we depleted ATG13 or FIP200 using siRNAs and inhibited the ULK1/2-kinase activity with MRT68921, resulting in a significant reduction of mitophagy (**Fig. 4c-d**). Additionally, inhibiting the kinase activity of VPS34, a component of the PI3KC3-C1 complex (composed of VPS34, VPS15, Beclin1, and ATG14), with a small molecule inhibitor also abrogated mitophagy (**Fig. 4e**). These results indicate that WIPI1, WIPI2, and WIPI3 recruitment to mitochondria occurs downstream of BNIP3/NIX and upstream of the ULK1 and PI3KC3-C1 complexes. Thus, despite being recruited in an unprecedented sequence, the ULK1 and PI3KC3-C1 complexes are still required for BNIP3/NIX mitophagy.

To confirm the necessity of the ULK1 and PI3KC3-C1 complexes during DFP-induced mitophagy, we depleted FIP200, ATG13, and ULK1 or inhibited the kinase activities of ULK1/2 and VPS34 (**Fig. 4f-i and S5c-d**). This confirmed that ULK1 and PI3KC3-C1 complexes are essential for DFP-induced mitophagy. Notably, ULK1 inhibition completely blocked mitophagy, consistent with previous work ^32^, while inhibition of the structurally related kinase TBK1 did not inhibit DFP-induced mitophagy (**Fig. 4h**).

Our data suggest a model where WIPI1, WIPI2, and WIPI3 can initiate autophagosome biogenesis, requiring the ULK1 and PI3KC3-C1 complexes but not TBK1. While TBK1 plays an important and sometimes essential role during selective autophagy initiated by soluble cargo receptors, our findings reveal that transmembrane mitophagy receptors BNIP3/NIX induce selective autophagy independent from TBK1, highlighting a critical distinction between selective mitochondrial turnover by soluble versus transmembrane cargo receptors.

### WIPI2 and WIPI3 bind directly to the ULK1 complex via ATG13/101

Given that BNIP3/NIX cannot directly recruit FIP200 but still require activation of the ULK1 complex downstream of the WIPIs, we aimed to elucidate how the FIP200/ULK1 complex is recruited and define the sequence in which the autophagy machinery components assemble in this pathway. We hypothesized that ATG16L1 might act as a bridging factor, given its known interactions with both WIPI2 and FIP200 ^51,52^. To test this, we generated a WIPI2 mutant (R108E/R125E) that is deficient in ATG16L1-binding ^51^. Upon rapalog treatment, mitophagy was induced not only by wild-type FKBP-GFP-WIPI2 but also by the ATG16L1-binding deficient WIPI2 mutant (**Fig. 5a**). This suggests that WIPI2 can recruit the ULK1 complex independently of its ATG16L1 binding ability. This finding aligns with our observation that BNIP3/NIX occupy the ATG16L1-binding site on WIPI2, indicating that these interactions are likely mutually exclusive, and that the R108E/R125E mutant co-immunoprecipitates more ULK1 ^53^.

**Figure 5.**
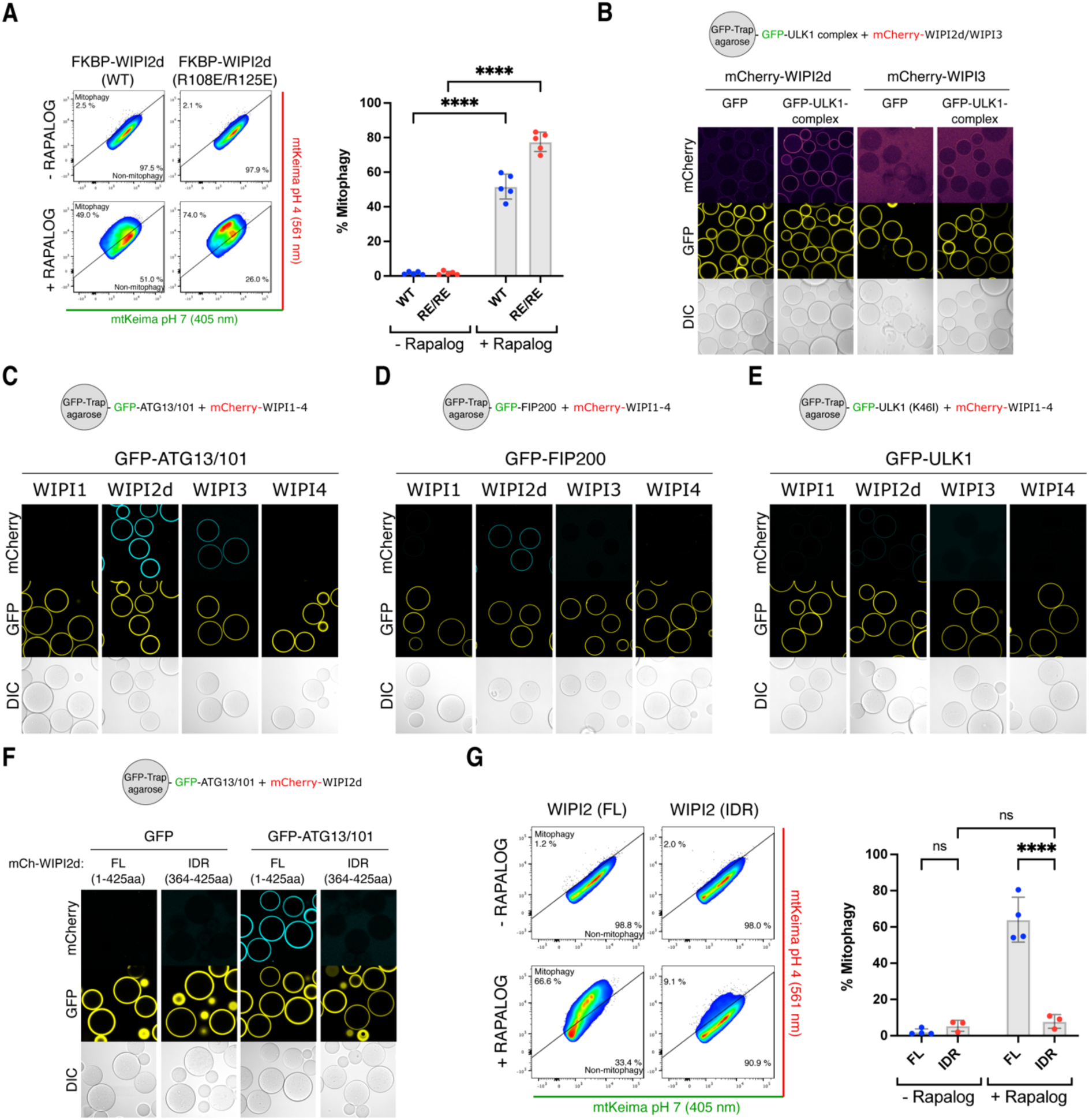
WIPI2 and WIPI3 bind directly to the ULK1 complex. **(A)** Mitophagy flux was measured by flow cytometry in wild-type HeLa cells expressing Fis1-FRB, FKBP-GFP-WIPI2 wild-type (WT) or ATG16L1-binding mutant R108E/R125E, and mt-mKeima, not induced or induced for 24 h by rapalog treatment. **(B)** Microscopy-based bead assay of agarose beads coated with GFP-tagged ULK1 complex (composed of FIP200-GFP, ULK1, ATG13, ATG101) and incubated with mCherry-tagged WIPI proteins. **(C)** As in (B) but with GFP-tagged ATG13/101 subcomplex and incubated with mCherry-tagged WIPI proteins. (**D**) As in (B) but with GFP-tagged FIP200 coated beads and incubated with mCherry-tagged WIPI proteins. (**E)** As in (B) but with GFP-tagged kinase dead ULK1 (K46I) coated beads and incubated with mCherry-tagged WIPI proteins. **(F)** As in (B) but with GFP-tagged ATG13/101 coated agarose beads incubated with mCherry-tagged full-length (FL) or IDR-only (residues 364-425) WIPI2d. (**G**) Mitophagy flux was measured by flow cytometry in wild-type HeLa cells expressing Fis1-FRB, full-length (FL) or IDR-only (364-425aa) FKBP-GFP-WIPI2, and mt-mKeima, not induced or induced for 24 h by rapalog treatment. Two-way ANOVA with Šídák’s multiple comparisons test in (A,G). ****P<0.0001. ns, not significant.

We then investigated whether WIPI proteins might directly bind the ULK1 complex. Indeed, WIPI2d and WIPI3 were recruited to beads coated with GFP-tagged ULK1 complex (**Fig. 5b**), with WIPI2d showing stronger binding than WIPI3. To identify which ULK1 complex subunits interact with WIPI proteins, we incubated mCherry-tagged WIPIs with individual ULK1 complex subunits. WIPI2d bound to the heterodimeric ATG13/101 subcomplex and weakly to FIP200, but not the ULK1 kinase subunit (**Fig. 5c-e**). WIPI3 bound only to the ATG13/101 subcomplex.

Structurally, the four WIPI proteins share a similar seven blade ß-propeller domain, with each blade composed of four antiparallel ß-strands ^54-56^. WIPI2 contains a binding site for ATG16L1 between blades 2 and 3 ^57,58^. Both WIPI1 and WIPI2 have a C-terminal intrinsically disordered region (IDR), while WIPI3 lacks this IDR but still binds ATG13/101. We therefore hypothesized that the interaction is mediated by the ß-propeller domains.

To test this, we attempted to purify WIPI2d without its C-terminal IDR, but its low solubility prevented successful purification of the ß-propeller domain alone. Instead, we purified the C-terminal IDR and, consistent with our hypothesis, found that it was unable to recruit ATG13/101 to mCherry-WIPI2d-IDR coated beads (**Fig. 5f**). Additionally, when we artificially tethered the IDR of WIPI2d to the mitochondrial surface in HeLa cells, robust mitophagy induction was no longer observed, unlike when the full-length WIPI2d was tethered (**Fig. 5g**).

These findings suggest the existence of a novel autophagy initiation complex involving the ß-propeller domains of WIPI proteins and the ATG13/101 subcomplex.

### Biochemical characterization of the WIPI-ULK1 autophagy initiation complex

To structurally characterize the WIPI-ATG13/101 mitophagy initiation complex in more detail, we set out to identify the minimal binding region between WIPI proteins and ATG13/101. ATG13 contains a HORMA domain and a C-terminal IDR region ^59-61^, while ATG101 only contains a HORMA domain necessary for dimerization with ATG13 ^60,61^. We investigated whether WIPI proteins bind to the ATG13/101 HORMA dimer or the ATG13 IDR by incubating WIPI2d and WIPI3 with either the ATG13 IDR or the ATG13/101 HORMA dimer lacking the IDR. Our results showed that WIPI2d and WIPI3 bind to the ATG13 IDR but not the HORMA domain dimer (**Fig. 6a**).

**Figure 6.**
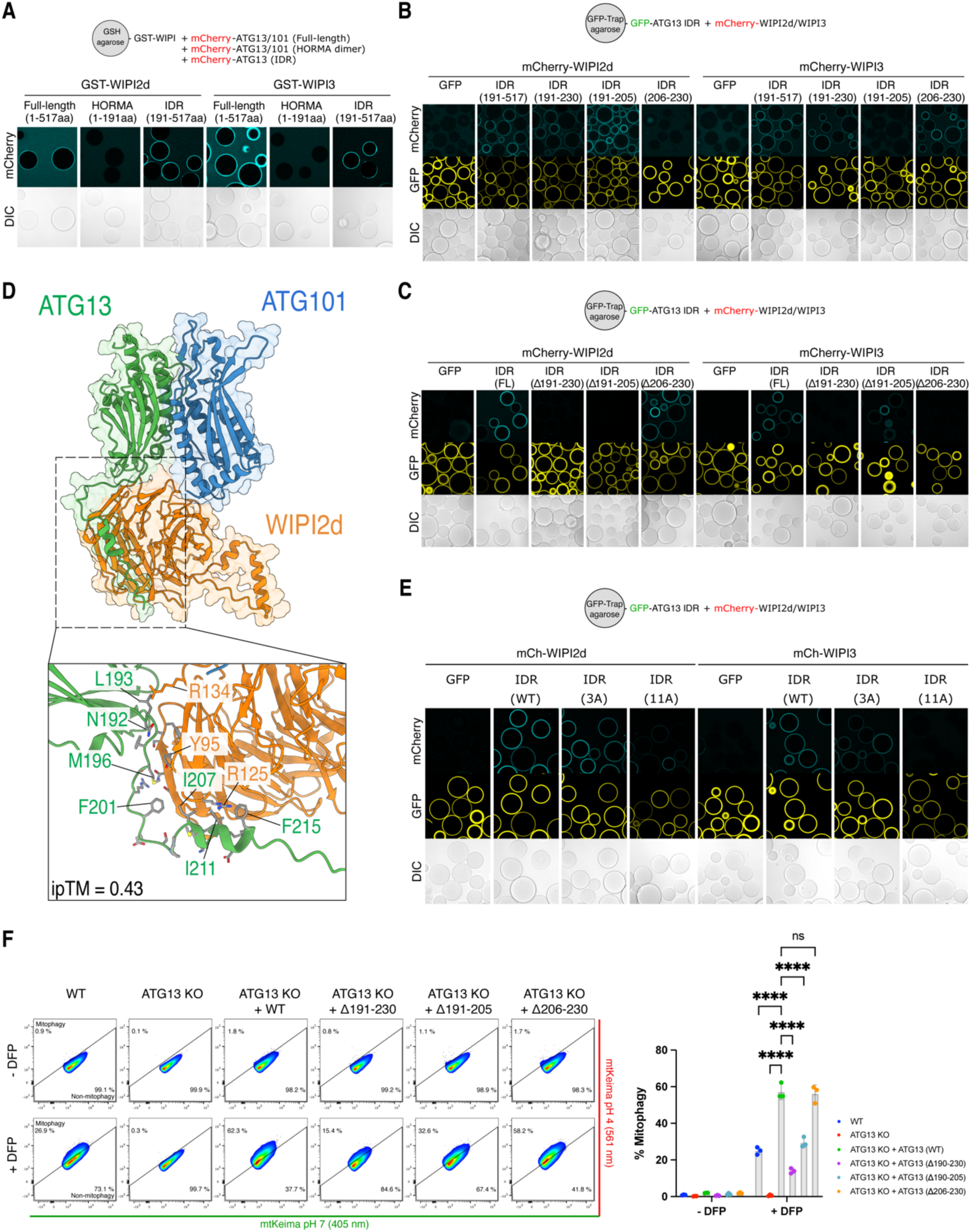
Biochemical characterization of the WIPI-ULK1 mitophagy initiation complex. **(A)** Microscopy-based bead assay of agarose beads coated with GST-tagged WIPI2d or WIPI3 and incubated mCherry-tagged ATG13/101 complex which was composed of full-length ATG13 (mCh-ATG13/101), HORMA-domain only (mCh-HORMA; ATG13 1-191aa/101), or IDR only (mCh-IDR; ATG13 191-517aa). **(B)** As in (A) but with GFP-tagged ATG13 IDR coated beads, either as full IDR (191-517aa) or fragments (191-230aa), (191-205aa), or (206-230), and incubated with mCherry-tagged WIPI2d or WIPI3. **(C)** As in (A) but with GFP-tagged ATG13 IDR coated beads, either as full IDR (191-517aa) or with variants containing deletion fragments (Δ191-230aa), (Δ191-205aa), or (Δ206-230), and incubated with mCherry-tagged WIPI2d or WIPI3. (**D**) AlphaFold predicted structure of WIPI2d (orange) and ATG13 (green) plus ATG101 (blue) with zoom in on the interaction interface. Note that the indicated residue numbers for WIPI2 correspond to their residue number in the WIPI2d sequence (which match residue numbers Y113 and R143 in WIPI2b). Structures were trimmed for visual clarity. Displayed are ATG13 (residues 1-223), ATG101 (residues 1-218), and WIPI2d (residues 1-383). (**E**) As in (A) but with GFP-tagged ATG13 IDR (191-517aa) coated beads and incubated with mCherry-tagged WIPI2d or WIPI3. The IDR is composed of the indicated either the wild-type (WT), 3x Ala mutant (3A), or 11x Ala mutant (11A). (**F**) Mitophagy flux was measured by flow cytometry of wild-type (WT) or ATG13 knockout (KO) HeLa cells, where indicated rescued with ATG13 wild-type (WT), ATG13 lacking residues 191-230 (Δ191-230), ATG13 lacking residues 191-205 (Δ191-205), or ATG13 lacking residues 206-230 (Δ206-230), left untreated or treated with DFP for 24 h. One of three representative experiments is shown. Two-way ANOVA with Dunnett’s multiple comparisons test. ****P<0.0001. ns, not significant.

Next, we mapped the minimal binding region using truncated versions of ATG13. We found that the initial stretch of the ATG13 IDR (191-230aa) is both required and sufficient to bind both WIPI2d and WIPI3 (**Fig. 6b and S6**). Our biochemical mapping suggests that WIPI2d and WIPI3 bind neighboring sequences on the ATG13 IDR (residues 191-202 for WIPI2d; residues 206-230 for WIPI3). We confirmed this by expressing the ATG13 IDR alone, without the HORMA domain, and deleting the entire binding region (residues 191-230) or only the minimal binding regions for WIPI2d (residues 191-205) or WIPI3 (residues 206-230). The results confirmed that the ATG13 IDR could still recruit WIPI2d if residues 191-205 were present, and WIPI3 if residues 206-230 were present (**Fig. 6c**).

To identify the interacting residues within these minimal binding regions, we predicted the structure of the complex using AF2 Multimer. After removing the ten most carboxyl-terminal residues from WIPI2d, which were incorrectly predicted to bind the HORMA dimer, AF2 Multimer correctly predicted that WIPI2d binds the initial segment of the ATG13 IDR (**Fig. 6d**). The prediction suggested that approximately 20 residues interact directly with the WIPI2d ß-propeller domain. To validate this, we created two ATG13 IDR variants: one with three residues and another with eleven residues replaced by alanine. Only the 11x Ala mutant abrogated the interaction, demonstrating that an extended stretch of the ATG13 IDR interacts with WIPI2d (**Fig. 6e**).

We then assessed the functional relevance of the identified binding interface during BNIP3/NIX mitophagy by measuring mitophagy flux in wild-type HeLa cells, ATG13 knockout cells, and ATG13 knockout cells rescued with wild-type or mutant ATG13 (Δ190-230, Δ190-205, Δ206-230) (**Fig. 6f**). DFP treatment induced mitophagy in approximately 20% of wild-type HeLa cells, which was completely abrogated in ATG13 knockout cells but rescued to nearly 60% with wild-type ATG13 overexpression. The ATG13 Δ190-230 mutant exhibited a significant defect, reducing mitophagy to 13%. The ATG13 Δ190-205 mutant displayed an intermediate phenotype with approximately 30% mitophagy, while the ATG13 Δ206-230 mutant showed a near wild-type phenotype with 57% mitophagy.

Our results demonstrate that BNIP3/NIX initiate autophagosome biogenesis by recruiting WIPI proteins, which in turn recruit the upstream ULK1 complex. WIPI2d and WIPI3 binding to the initial segment of the ATG13 IDR is critical for the formation of the WIPI-ULK1 complex during BNIP3/NIX mitophagy.

### Flexibility in the productive assembly of autophagy machinery

Our findings reveal distinct assembly sequences during autophagosome biogenesis in the BNIP3/NIX versus PINK1/Parkin mitophagy pathways. Specifically, in BNIP3/NIX mitophagy, WIPI protein recruitment to mitochondria occurs upstream of the ULK1 and PI3KC3-C1 complexes, underscoring the crucial role of the WIPI-ATG13 interaction. This observation raises the question of whether this interaction is also important in other forms of selective or non-selective autophagy.

To investigate this, we first examined the role of ATG13 in basal autophagy. In ATG13 knockout cells, we observed significant accumulation of activated SQSTM1/p62 (**Fig. 7a**), a pattern also seen in FIP200 knockout cells ^4^. The elevated levels of heavily phosphorylated SQSTM1/p62 suggest a blockage in basal turnover of protein aggregates. Notably, reintroducing wild-type ATG13 or the Δ190-230 variant (which is deficient in BNIP3/NIX mitophagy) restored SQSTM1/p62 levels, indicating that the WIPI-ATG13 interaction is not essential for basal autophagy.

**Figure 7.**
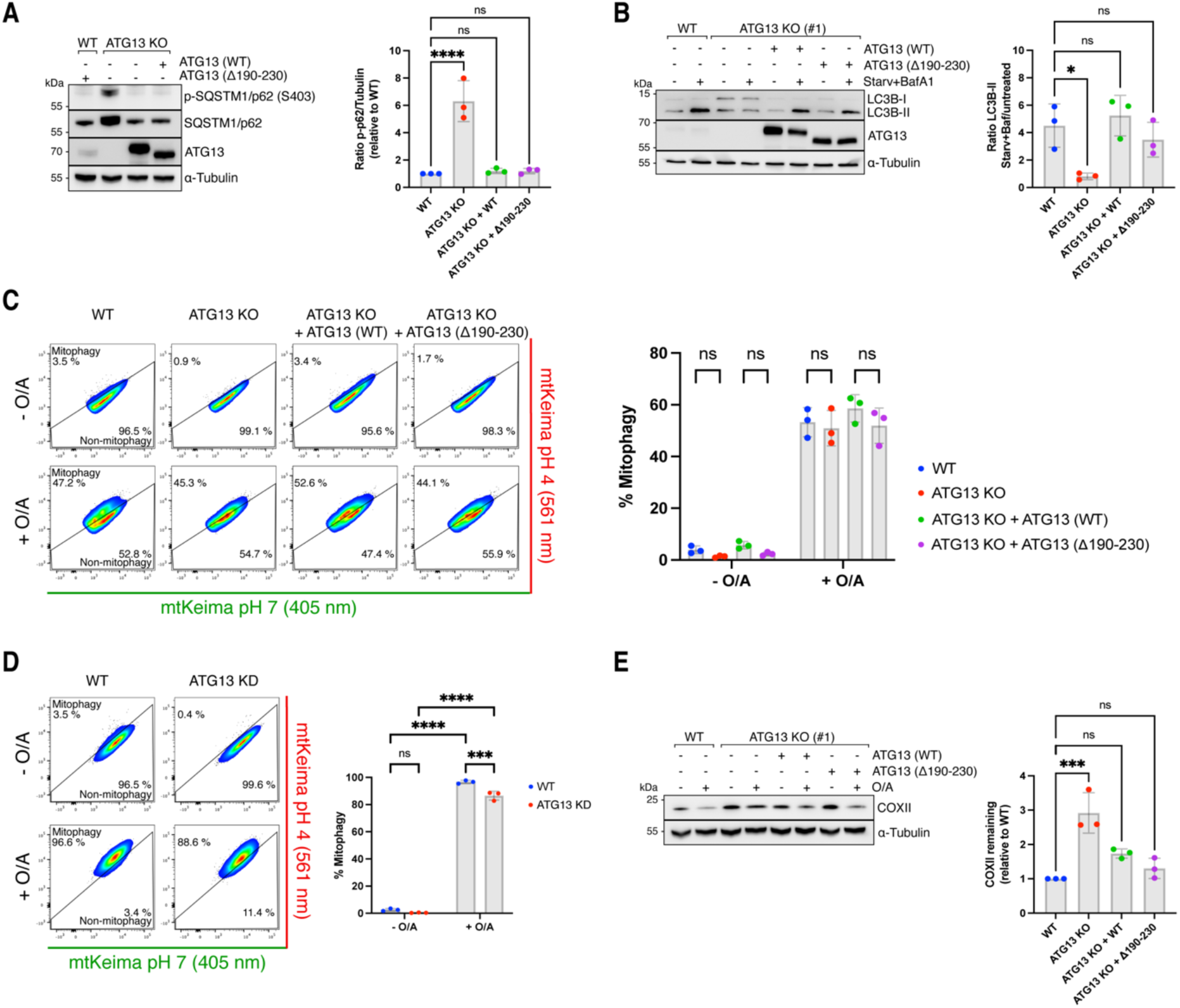
Distinct hierarchy of assembly between WIPI-driven mitophagy and FIP200-driven mitophagy or starvation-induced autophagy. **(A)** Immunoblotting for phosphorylated SQSTM1/p62 in wild-type (WT) or ATG13 knockout (KO) cells (clone #1), where indicated rescued with ATG13 WT or ATG13 lacking residues 190-230 (Δ190-230) **(B)** Immunoblotting for LC3B in the same cell lines as used in (A) but treated with 2 h starvation and Bafilomycin A1 (BafA1) where indicated. **(C)** Mitophagy flux was measured by flow cytometry of wild-type (WT) or ATG13 knockout (KO) HeLa cells, where indicated rescued with ATG13 wild-type (WT) or ATG13 lacking residues 190-230 (Δ190-230), left untreated or treated with O/A for 5 h. One of three representative experiments is shown. (**D**) As in (C) but with wild-type HeLa cells transfected with siRNAs targeting ATG13, left untreated or treated with O/A for 5 h. (**E**) Immunoblotting of COXII levels in wild-type (WT) or ATG13 knockout (KO) HeLa cells, overexpressing BFP-Parkin, and where indicated rescued with ATG13 wild-type (WT) or ATG13 lacking residues 190-230 (Δ190-230), left untreated or treated with O/A for 24 h. Densitometric analysis was performed for the percentage of COXII remaining relative to WT cells (mean ± s.d.) (n = 3 biologically independent experiments). One-way ANOVA with Dunnett’s multiple comparison test was performed. One-way ANOVA with Dunnett’s multiple comparisons test (A, E) or Tukey’s multiple comparisons test (B), or a Two-way ANOVA with Tukey’s multiple comparisons test (C-D). *P<0.05, ***P<0.001, ****P<0.0001. ns, not significant.

We then assessed the impact of the WIPI-ATG13 interaction on starvation-induced non-selective autophagy. In ATG13 knockout cells, lipidated LC3-II levels remained unchanged following starvation plus Bafilomycin A1 treatment, demonstrating a complete blockage of autophagy flux (**Fig. 7b**). However, this blockade was rescued by reintroducing either wild-type ATG13 or the Δ190-230 variant, suggesting that the WIPI-ATG13 complex is not critical for non-selective autophagy induction.

Next, we explored the role of the WIPI-ATG13 interaction in PINK1/Parkin mitophagy. Unlike BNIP3/NIX mitophagy, where ATG13 is absolutely essential, PINK1/Parkin mitophagy was only mildly affected by ATG13 deletion. Both ATG13 knockout and ATG13 siRNA-depleted cells showed a modest reduction in mitophagy flux but did not impair PINK1/Parkin mitophagy (**Fig. 7c-e**). This supports our model that BNIP3/NIX and soluble cargo receptors assemble the autophagy machinery in distinct sequences, explaining the differential requirement for ATG13.

Our data therefore show that transmembrane cargo receptors like BNIP3/NIX can recruit the autophagy machinery in a distinct order compared to soluble cargo receptors, and use a WIPI-driven pathway instead of a FIP200-driven pathway.

### Recruitment of WIPI proteins is a common feature of transmembrane cargo receptors

Inspired by our findings that BNIP3/NIX initiate autophagosome biogenesis by first recruiting WIPI proteins, we investigated if other transmembrane cargo receptors could also bind and recruit WIPIs. To explore this possibility, we performed an AF3 screen to identify additional candidate autophagy receptors that might interact with WIPI2. The predictions were ranked using the ipTM score, which estimates the quality of the complex based on predicted protein interfaces. The AF3 predictions identified potential interactions between WIPI2 and several transmembrane autophagy receptors, including the ER-phagy receptors TEX264 and FAM134C, as well as the mitophagy receptor FKBP8 (**Fig. 8a**). Notably, TEX264 (ipTM 0.54) and FKBP8 (ipTM 0.58) scored above the 0.5 threshold, similar to BNIP3 (ipTM 0.66) and NIX (ipTM 0.65). However, FAM134C (ipTM 0.46) scored slightly below this cut-off. We repeated the predictions with AF2 Multimer, which also predicted interactions for TEX264 and FKBP8 but not for FAM134C. Interestingly, TEX264 and FKBP8 were predicted to bind the same pocket on WIPI2 as BNIP3/NIX (**Fig. S7a-b**), suggesting a potentially conserved feature among different autophagy receptors.

**Figure 8.**
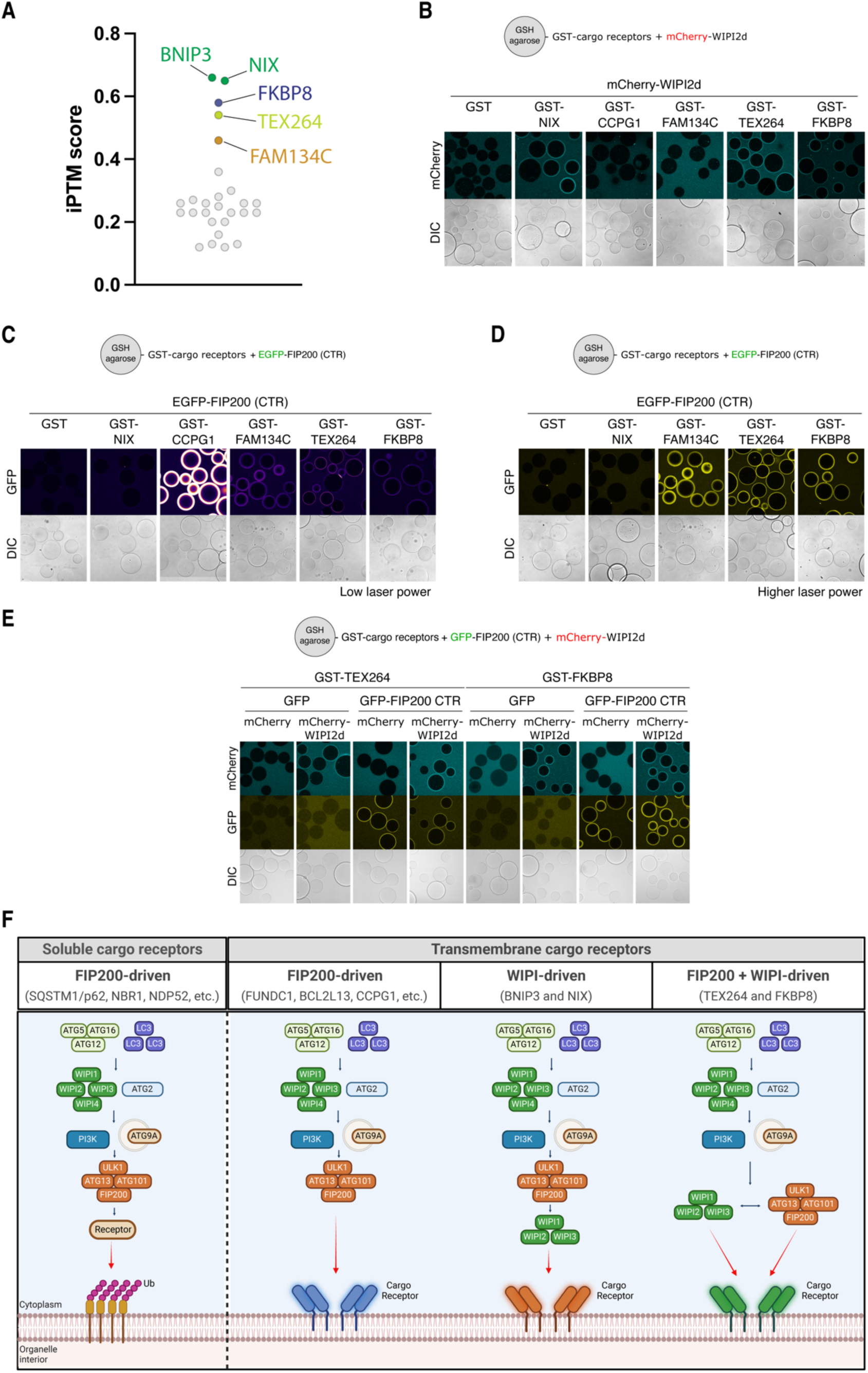
Several transmembrane cargo receptors can bind WIPI proteins. **(A)** AF3 screen for interaction between all known cargo receptors, soluble and transmembrane, and WIPI2. Predicted interactions are plotted for their ipTM score. **(B)** Microscopy-based bead assay of agarose beads coated with GST-tagged NIX, CCPG1, FAM134C, TEX264, and FKBP8 or GST alone as negative control, and incubated with mCherry-tagged WIPI2d. **(C-D)** As in (B), but with GFP-tagged C-terminal region of FIP200 (CTR). The laser power was either very low to visualize CCPG1-FIP200 interaction (C) or with higher laser power to visualize FAM134C, TEX264, FKBP8 and FIP200 interaction (D). In panel (C) we used the Fire LUT to better visualize the difference in binding strength between the different receptors. (E) As in (B), but with mCherry-tagged WIPI2d and/or GFP-tagged C-terminal region of FIP200 (CTR). (F) Schematic overview of the different selective autophagy pathways. Soluble cargo receptors are recruited to ubiquitinylated organelles and recruit the ULK1 complex through FIP200 to initiate autophagosome biogenesis. Transmembrane cargo receptors can initiate autophagosome biogenesis either through recruiting FIP200 or through recruiting WIPI proteins. The latter then recruit the ULK1 complex through interactions with ATG13, and in case of WIPI2 also through interaction with FIP200. Depending on the cargo receptor, autophagosome biogenesis can be initiated through FIP200- and/or WIPI-driven mechanisms.

We next tested these predicted interactions using recombinant proteins, focusing on TEX264, FKBP8, and FAM134C, with CCPG1 (ipTM 0.2) as a negative control. We expressed and purified the soluble domains of each receptor, replacing their transmembrane regions with GST (**Fig. S7c**). A microscopy-based bead assay was used to assess their capacity to bind mCherry-tagged WIPI2d. We observed that TEX264 and FKBP8 bound to WIPI2d, while FAM134C and CCPG1 did not (**Fig. 8b**). This suggests that WIPI-mediated autophagy initiation is a conserved mechanism across multiple organelles.

Next, we investigated whether TEX264 and FKBP8 can also bind FIP200 in addition to WIPI2d. We found that both TEX264 and FKBP8 could bind FIP200 (**Fig. 8c-d**), similar to FAM134C and CCPG1. This indicates that TEX264 and FKBP8 can recruit both FIP200 and WIPI2, whereas BNIP3/NIX exclusively recruit WIPI2. Notably, the binding strength for FIP200 was comparable between FAM134C, TEX264, and FKBP8, but significantly stronger for CCPG1, likely due to CCPG1’s dual FIR motifs, as previously demonstrated ^23^.

Finally, since TEX264 and FKBP8 can bind both FIP200 and WIPI2d, we examined whether these receptors could recruit both autophagy initiation arms simultaneously. We coated agarose beads with GST-tagged TEX264 or FKBP8 and incubated the cargo receptors with GFP-tagged FIP200 C-terminal region and mCherry-tagged WIPI2d. This revealed that both TEX264 and FKBP8 can bind and recruit FIP200 and WIPI2d at the same time (**Fig. 8e**), suggesting the potential formation of a mega-initiation complex.

In summary, our study reveals that selective autophagy can be initiated through two distinct modes: either by first recruiting FIP200 or by recruiting WIPI proteins (**Fig. 8f**). While WIPI proteins were previously considered downstream factors, our work shows that several transmembrane cargo receptors contain motifs enabling them to bind and recruit WIPI proteins to initiate autophagosome biogenesis. This finding highlights an unexpected flexibility in the hierarchical assembly of the autophagy machinery during autophagosome formation.

## DISCUSSION

In this study, we uncover the mechanisms by which selective autophagy receptors can initiate selective autophagy, expanding our understanding beyond the well-characterized pathways involving soluble cargo receptors. Through a combination of biochemical reconstitution, cell biology, AF modeling, and molecular dynamics simulations, we have delineated distinct pathways utilized by different transmembrane receptors to initiate selective autophagy.

Our findings demonstrate that various transmembrane cargo receptors, including FUNDC1, BCL2L13, CCPG1, and FAM134C, recruit the autophagy machinery through interaction with FIP200. This mechanism mirrors the way soluble cargo receptors initiate autophagosome biogenesis, underscoring the conservation of autophagy initiation processes across different receptor types. The depletion of ULK1-complex components was shown to impair mitophagy driven by FUNDC1 and BCL2L13 ^62,63^, and co-immunoprecipitation experiments confirmed that ULK1 interacts with both receptors ^62,63^, highlighting the crucial role of the ULK1 complex in these processes. Moreover, the binding of these transmembrane receptors to the C-terminal domain of FIP200 further emphasizes the critical role of FIP200 in autophagosome biogenesis^1,64-71^, supporting the notion that transmembrane receptors engage autophagy machinery through conserved motifs.

In stark contrast, NIX and BNIP3 utilize a fundamentally different strategy to initiate mitophagy, which does not involve direct interaction with FIP200 or other upstream components of the canonical autophagy pathway. While these results do not rule out the possibility that BNIP3/NIX can bind to FIP200 under different conditions than those tested here, we were unable to establish a direct interaction between the two mitophagy receptors and FIP200. Instead, our data demonstrate that NIX and BNIP3 recruit downstream WIPI proteins to the mitochondrial surface, which in turn engage the upstream ULK1 complex via ATG13/101 subunits. This order of recruitment represents a previously unrecognized mode of autophagy initiation, highlighting an extraordinary flexibility in the assembly and activation of autophagy machinery.

The interaction of BNIP3/NIX with ATG13 via WIPI2 and WIPI3 suggests a mechanism where downstream autophagy factors can facilitate the recruitment of upstream components, thereby reversing the classical sequence of autophagy initiation events. This reverse recruitment mechanism was validated by our experiments showing that tethering WIPI proteins to the mitochondrial surface is sufficient to initiate autophagosome biogenesis, contingent upon the presence of functional ULK1 and PI3KC3-C1 complexes.

Further biochemical characterization and AF modeling provided structural insights into the interactions between WIPI proteins and the ULK1 complex. We identified specific binding interfaces within the β-propeller domains of WIPI2 and WIPI3 that interact with the ATG13/101 subcomplex. These interactions were essential for mitophagy, as mutations disrupting the WIPI-ULK1 complex formation abrogated autophagic flux. Given that WIPI2 and WIPI3 bind neighboring sequences on the ATG13 IDR, and that BNIP3/NIX form dimers in their active state^35^, suggests that the same ATG13 molecule might interact with two WIPI molecules. This interaction could thus result in the formation of one large mitophagy initiation complex composed of BNIP3/NIX-WIPI2-WIPI3-ATG13/101-FIP200-ULK1.

The recruitment of WIPI proteins by NIX and BNIP3 and their ability to initiate mitophagy independently of TBK1, a kinase often essential in soluble cargo receptor-mediated autophagy^1,66-69^, together with the critical role of ATG13 during BNIP3/NIX mitophagy but not PINK1/Parkin mitophagy, delineates a critical distinction between the autophagy pathways initiated by soluble versus transmembrane cargo receptors. This distinction not only underscores the diversity of autophagy initiation mechanisms but also suggests that cells might employ different strategies to ensure the turnover of specific organelles under varying physiological conditions.

Importantly, the WIPI-ATG13 axis we uncover here may be widely used by transmembrane cargo receptors as we found that another mitophagy transmembrane receptor, FKBP8, as well as the ER-phagy receptor TEX264, bind to WIPI2. Notably, these receptors also bind to FIP200, suggesting that they can activate selective autophagy through both the WIPI and FIP200 pathways.

Overall, our study advances our understanding of the molecular mechanisms underlying transmembrane receptor-mediated selective autophagy. The discovery of distinct pathways for different receptors enriches the conceptual framework of autophagy and opens new avenues for targeted therapeutic interventions in diseases characterized by dysfunctional autophagy. Future studies will be necessary to further dissect the regulatory mechanisms governing these pathways and to explore their implications in various cellular contexts and disease states.

**Figure S1.**
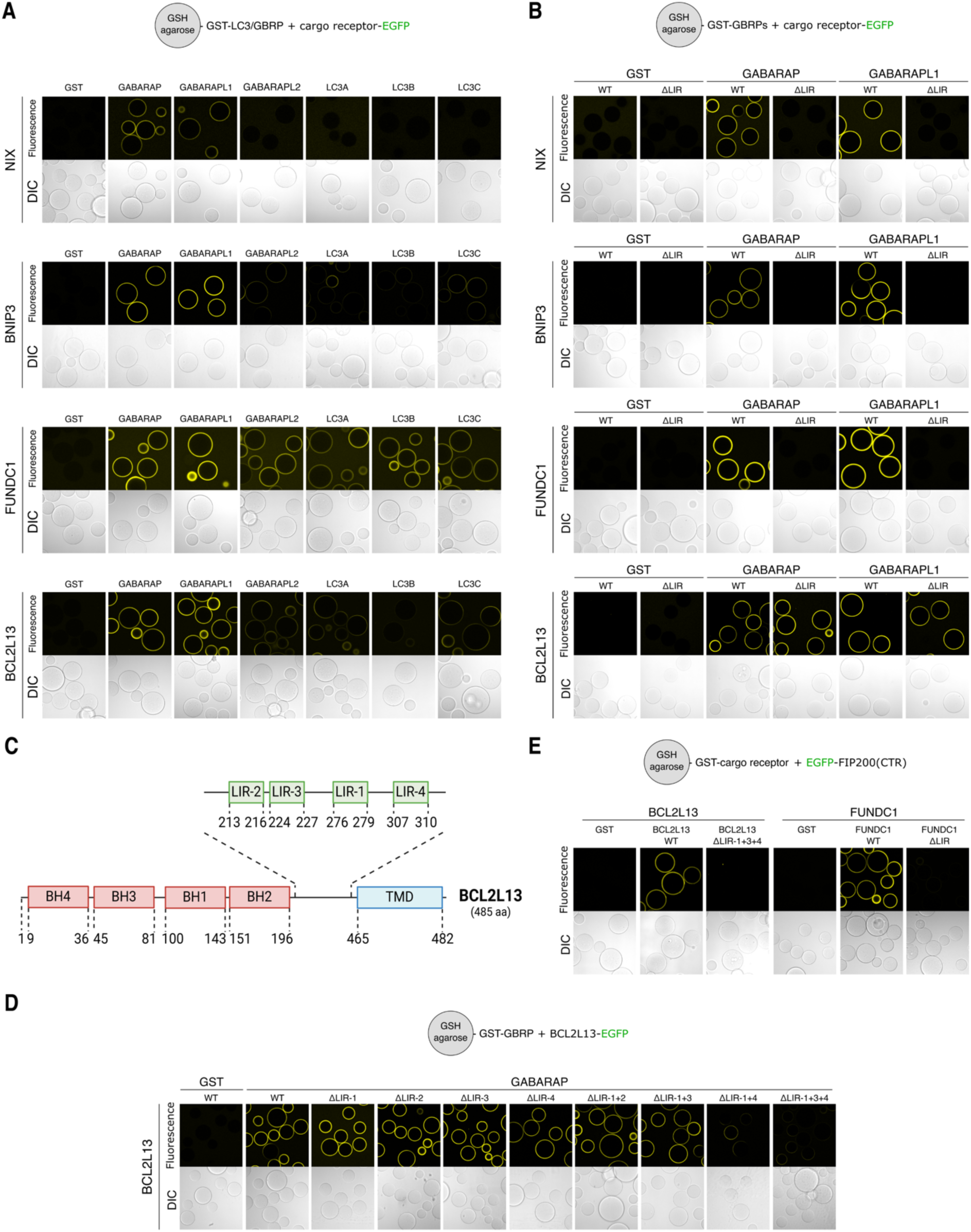
In vitro validation of mitophagy cargo receptors and their LIR/FIR motifs. **(A)** Microscopy-based bead assay of agarose beads coated with GST-tagged LC3A/B/C or GBRP/GBRPL1/GBRPL2 and incubated with GFP-tagged cargo receptors FUNDC1, BCL2L13, NIX, and BNIP3. (**B**) As in (A) but with wild-type (WT) or alanine-mutated LIR-motifs (ΔLIR) of the GFP-tagged cargo receptors. (**C**) Schematic of domain structure of BCL2L13 with the candidate LIR/FIR motifs indicated with residue numbers. LIR-1 was previously annotated in literature as the active LIR motif. (**D**) As in (A), but with different alanine-mutated variants of the different LIR-motifs (ΔLIR) of GFP-tagged BCL2L13. (**E**) As in (A) but with GST-tagged cargo receptors and GFP-tagged C-terminal region (CTR; 1429-1591aa) of FIP200.

**Figure S2.**
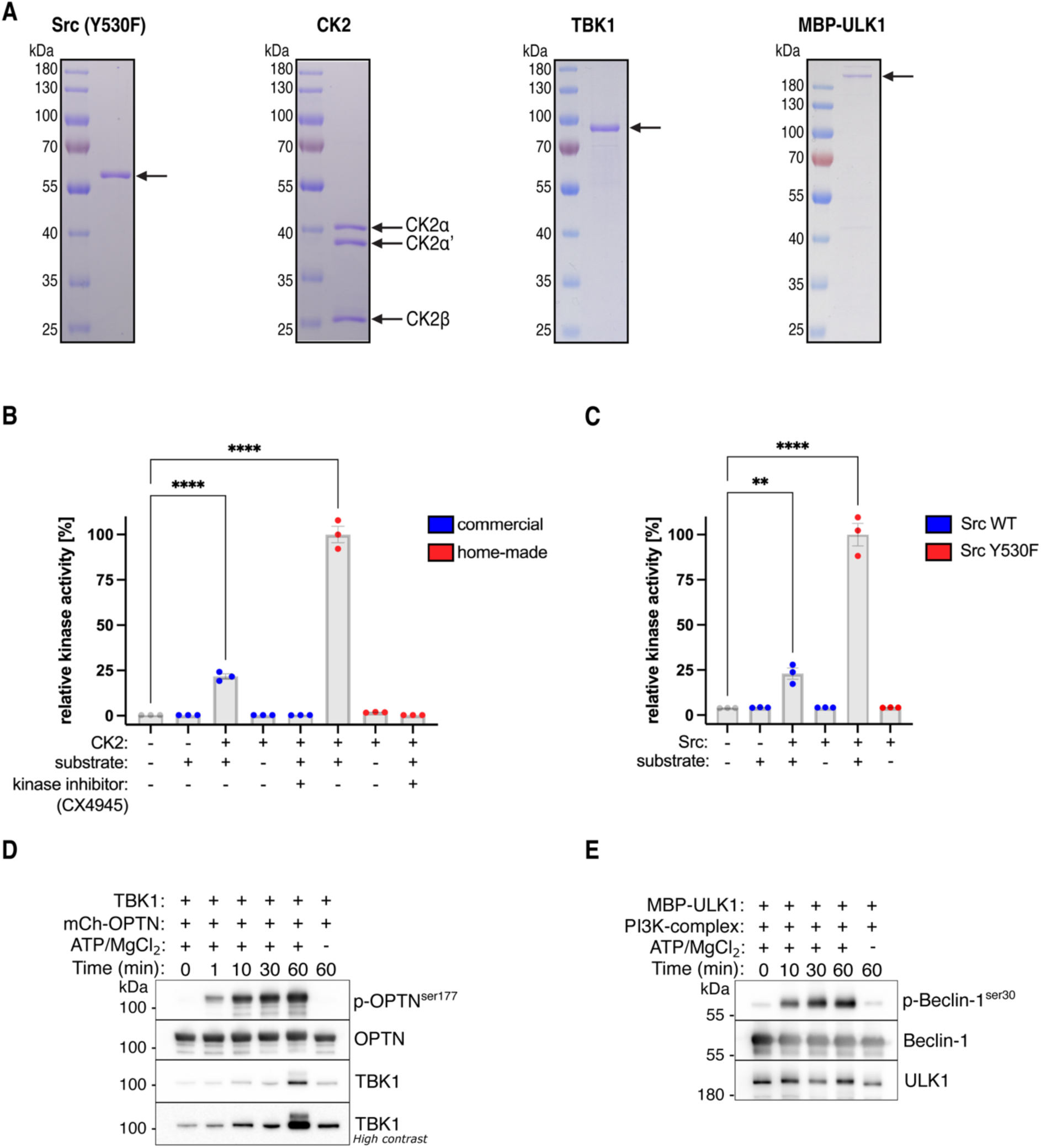
Purified kinases and validation of their activity. **(A)** Representative SDS-PAGE gels of purified Src (Y530F), CK2 complex, TBK1, and MBP-ULK1. Arrows indicate the predicted molecular weight. (**B-C**) Measurement of kinase activity using a plate-reader based read-out. Kinases were incubated with or without a substrate peptide or kinase inhibitor. Kinase activity was compared between our purified CK2 complex (home-made) and commercially available CK2, or between wild-type (WT) and Y530F mutant Src. (**D**) Measurement of kinase activity by mixing recombinantly purified mCherry-OPTN and TBK1 for the indicated time and western blot analysis using antibodies for phosphorylated OPTN (S177) as a read out for TBK1 activity. (**E**) As in D, but after mixing recombinantly purified MBP-ULK1 and the PI3KC3-C1 complex (composed of ATG14, Beclin-1, Vps15, Vps34) for the indicated time and using antibodies for phosphorylated Beclin-1 (Ser30) as a read out for ULK1 activity. One-way ANOVA with Dunnett’s multiple comparisons test (B, C). **P<0.005, ****P<0.0001. ns, not significant.

**Figure S3.**
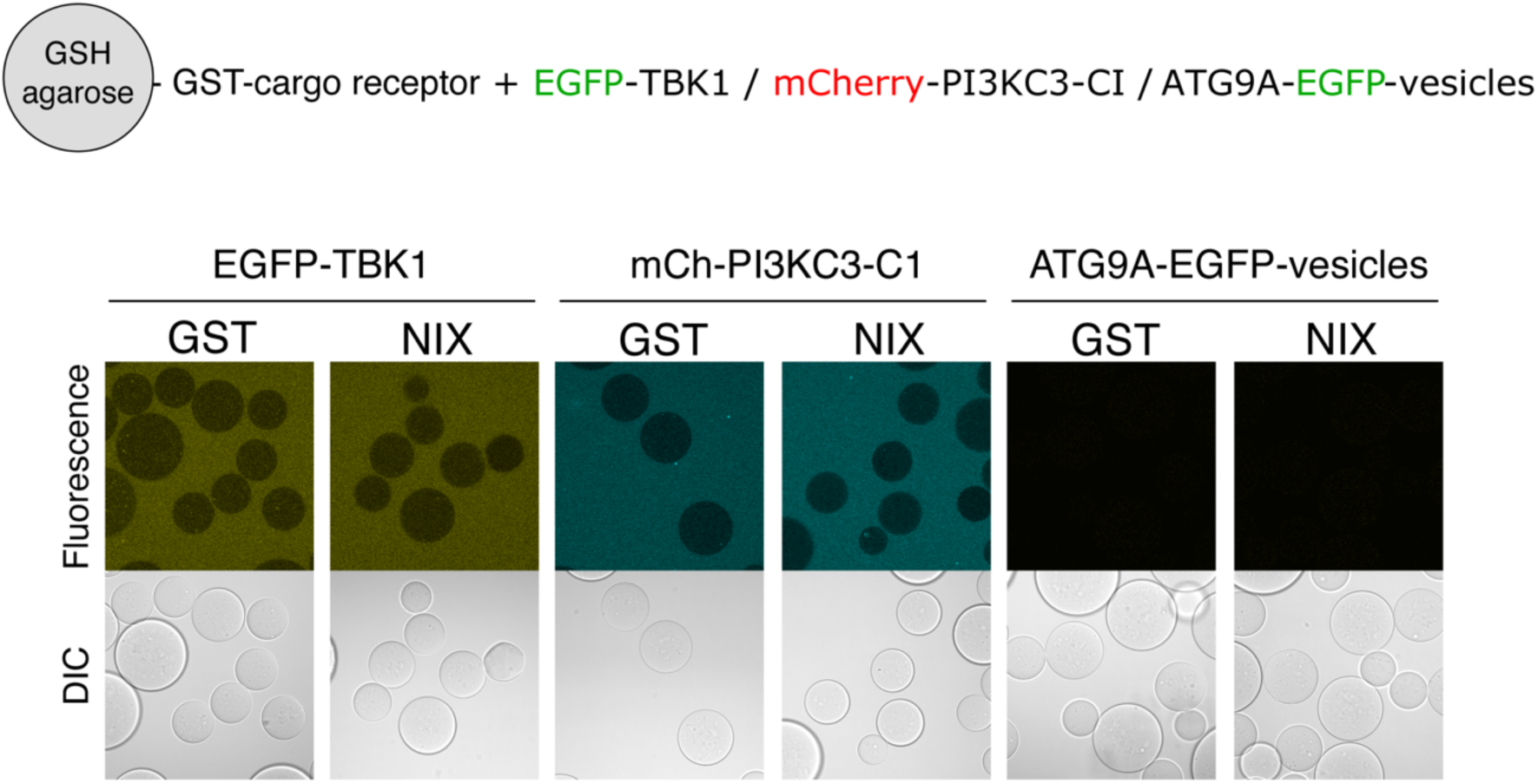
NIX does not interact with TBK1, PI3KC3-C1 complex, or purified ATG9A-vesicles. Microscopy-based bead assay of agarose beads coated with GST-tagged NIX and incubated with GFP-tagged TBK1, mCherry-tagged PI3KC3-C1, or GFP-tagged ATG9A-vesicles purified from HAP1 cells. GST served as negative control.

**Figure S4.**
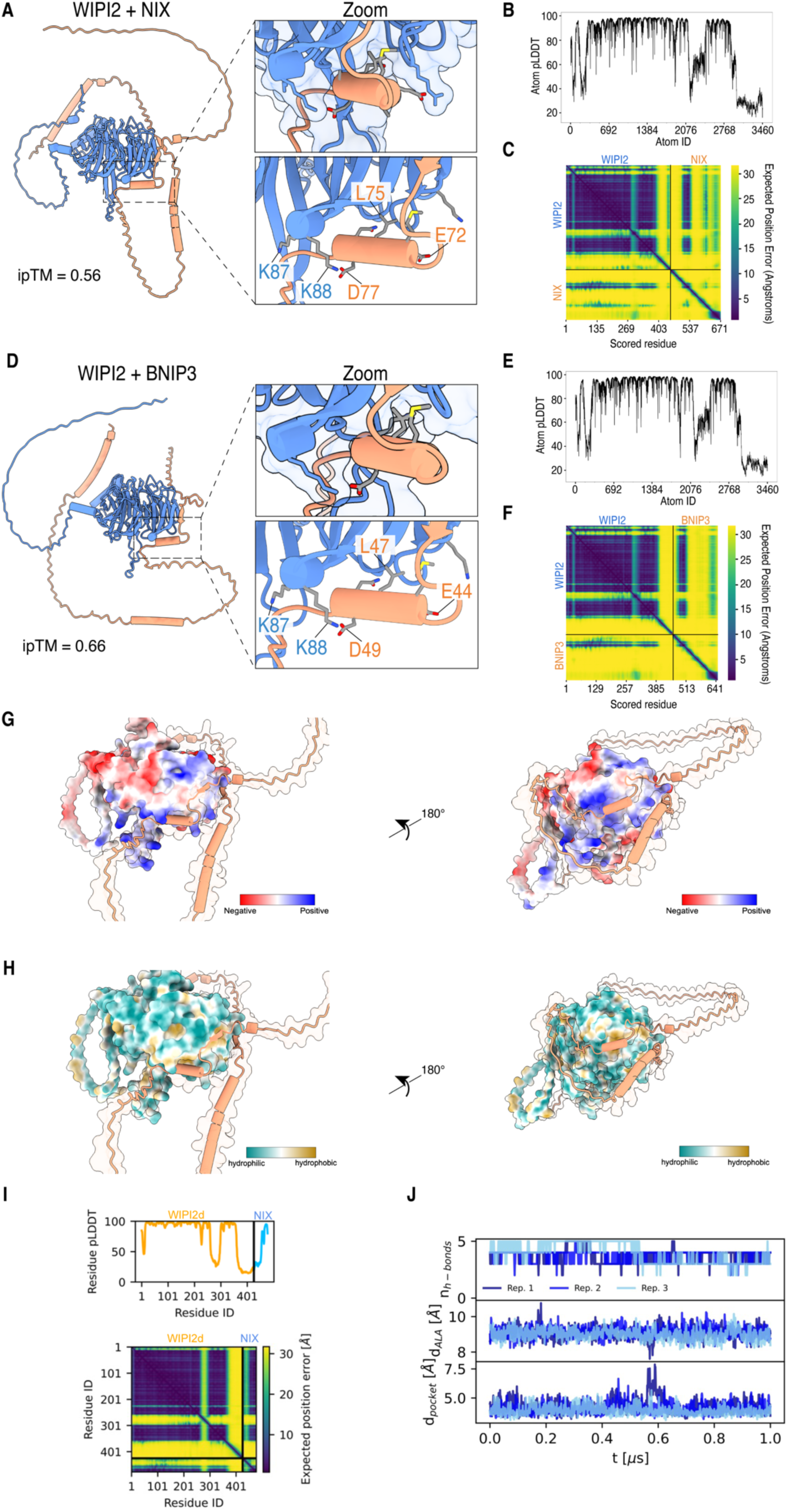
AlphaFold-2 prediction and MD simulations of BNIP3/NIX-WIPI2 complex. **(A)** AlphaFold-2 predicted structure of NIX (orange) and WIPI2 (blue) with zoom in on the interaction interface. (**B-C**) pLDDT and PAE plots for NIX-WIPI2 structure. (**D**) AlphaFold-2 predicted structure of BNIP3 (orange) and WIPI2 (blue) with zoom in on the interaction interface. (**E-F**) pLDDT and PAE plots for BNIP3-WIPI2 structure. (**G**) Predicted structure for the NIX-WIPI2 complex with the surface of WIPI2 colored based on electrostatics. (**H**) Predicted structure for the NIX-WIPI2 complex with the surface of WIPI2 colored based on hydrophobics. Note that the indicated residue numbers for WIPI2 correspond to their residue number in the WIPI2d sequence (which match residue numbers K105 and K106 in WIPI2b). **(I)** Residue pLDDT and PAE scores for the prediction in Fig. 2i. (**J**) The NIX W36A/L39A (ΔLIR) mutant does not bind the cryptic pocket of WIPI2d. Number of backbone h-bonds n_h-bonds_ between the LIR of NIX and WIPI2d, insertion depth d_ALA_ of NIX ΔLIR A36, and minimum heavy atom distance d_pocket_ between WIPI2d F169 and I133 from three 1 μs MD simulations.

**Figure S5.**
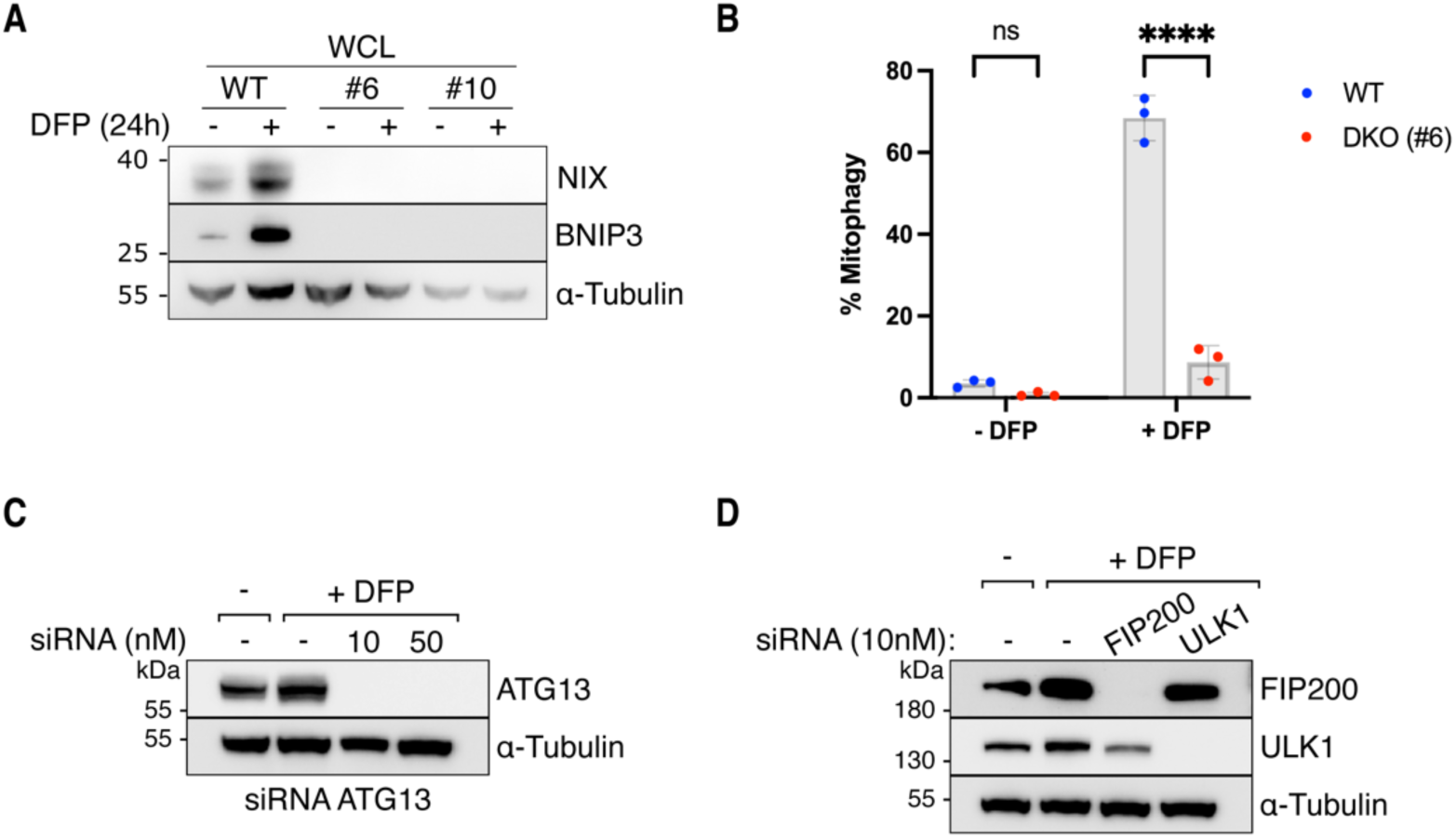
Validation knockout and knockdown cell lines. **(A)** Analysis of whole cell lysates (WCL) by SDS-PAGE and western blotting for NIX/BNIP3 double knockout clones #6 and #10, with and without induction of mitophagy by 24 h of DFP treatment. (**B**) Mitophagy flux was measured by flow cytometry of wild-type (WT) or NIX/BNIP3 double knockout (DKO) HeLa cells (clone #6), left untreated or treated with DFP for 24 h. (**C**) Analysis of knockdown efficiency for ATG13. HeLa cells were transfected 72 h prior to the FACS experiment, treated with DFP for 24 h to induce mitophagy, and analyzed by flow cytometry. Cells were collected after the experiment and analyzed by SDS-PAGE and western blotting. The concentration of 10 nM was used for the FACS experiment represented in the manuscript. (**D**) As in (C), but for HeLa cells transfected with siRNAs against FIP200, ULK1 or scrambled as a control (-). Two-way ANOVA with Tukey’s multiple comparisons test. ****P<0.0001. ns, not significant.

**Figure S6.**
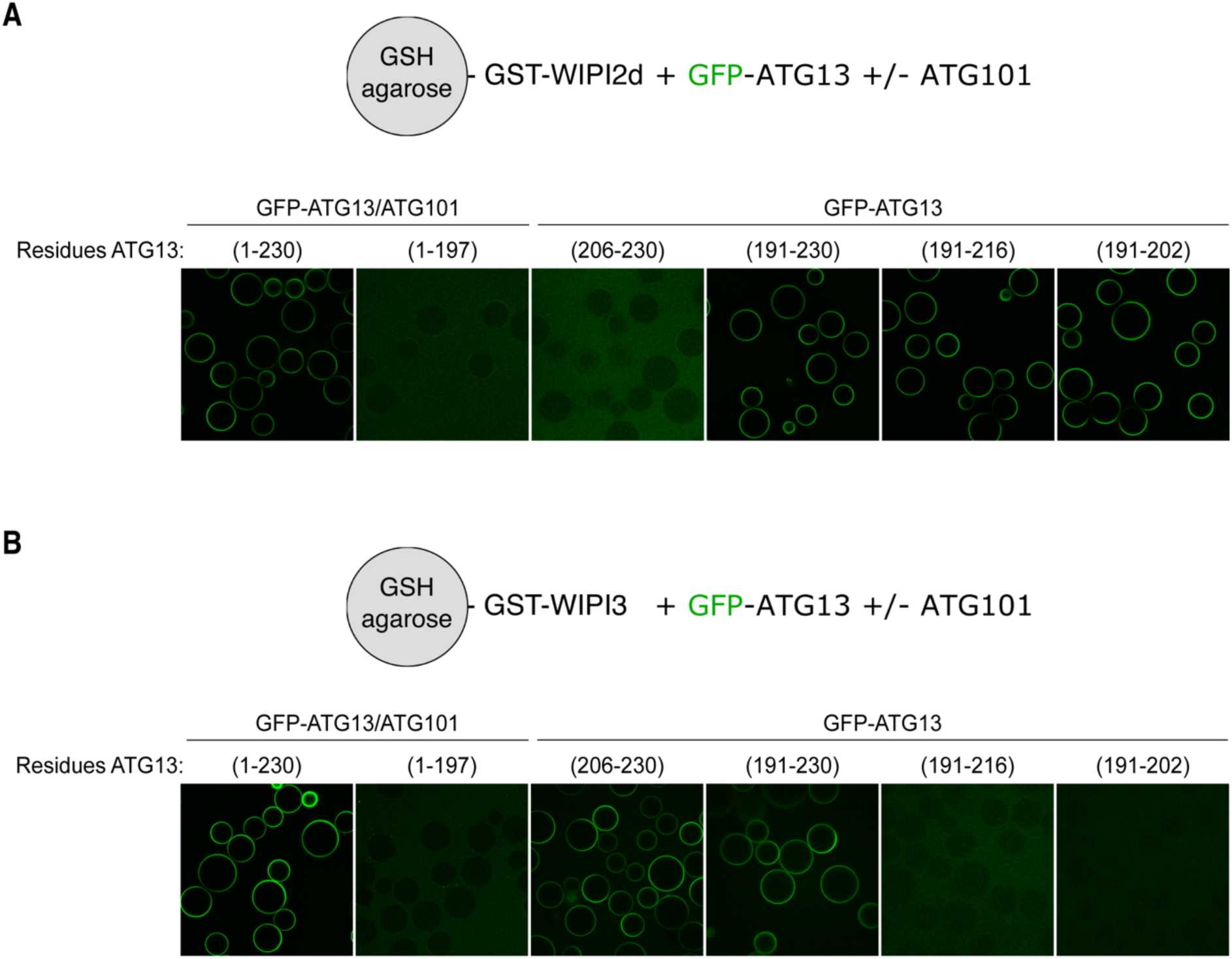
Biochemical mapping of binding sites of WIPI-ATG13 interaction. Microscopy-based bead assay of agarose beads coated with GST-tagged (**A**) WIPI2d or (**B**) WIPI3 and incubated with GFP-tagged ATG13/ATG101 subcomplex or fragments of ATG13 alone.

**Figure S7.**
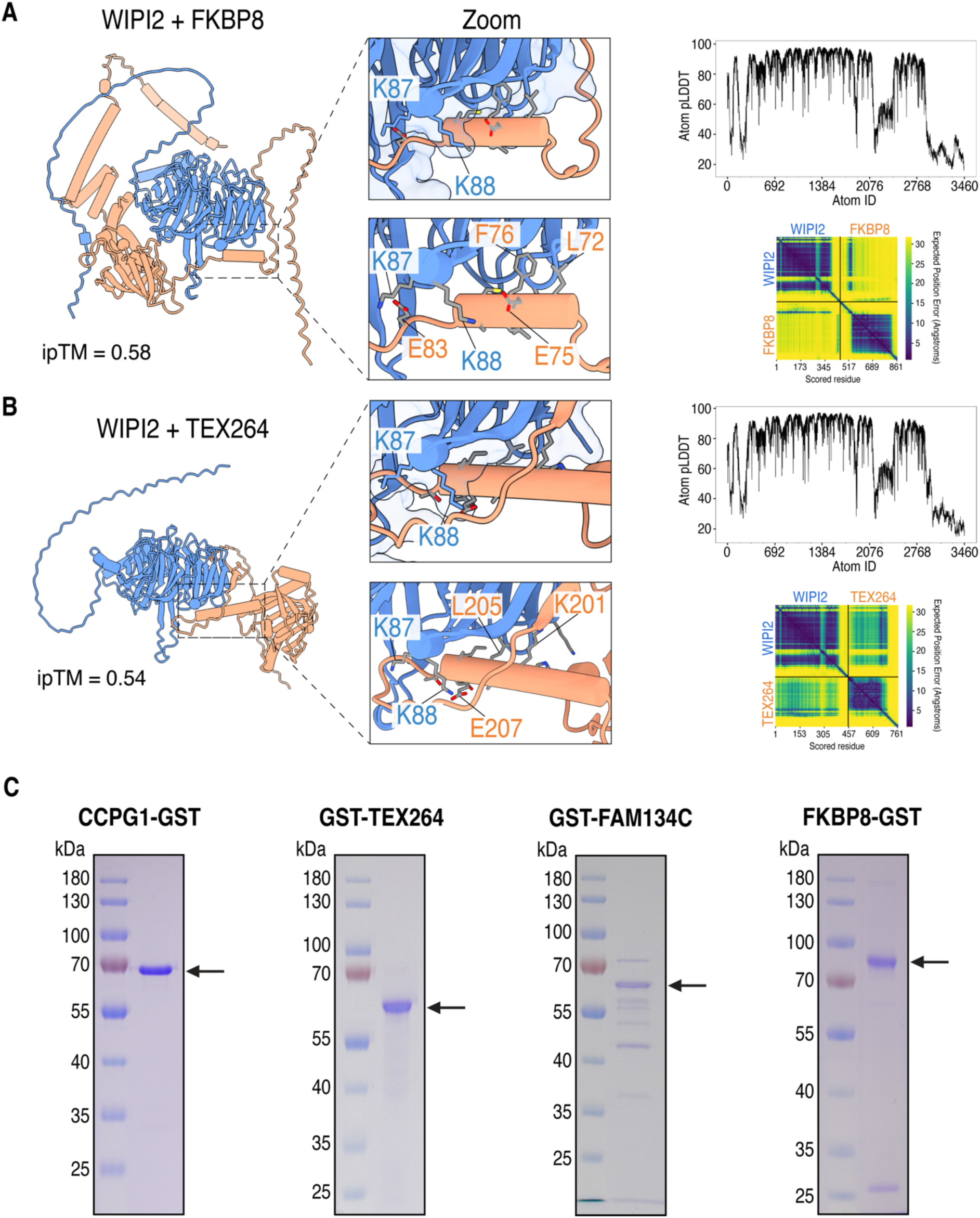
AlphaFold-2 prediction of WIPI2d and transmembrane cargo receptors. (**A-B**) AlphaFold-3 predicted structure for WIPI2 with (A) TEX264, or (B) FKBP8, with zoom in on the interaction interface. Note that the indicated residue numbers for WIPI2 correspond to their residue number in the WIPI2d sequence (which match residue numbers K105 and K106 in WIPI2b). pLDDT plots and predicted alignment error (PAE) heatmap are also shown. (**C**) Representative SDS-PAGE gels stained with Coomassie Brilliant Blue of purified CCPG1(1-212aa)-GST, GST-TEX264(28-313aa), GST-FAM134C(250-466aa), and FKBP8(1-391aa)-GST. Arrows indicate the predicted molecular weight.

## MATERIAL AND METHODS

### Reagents

The following chemicals were used in this study: Rapalog A/C hetero-dimerizer (635057, Takara), Bafilomycin A1 (sc-201550, Santa Cruz Biotech), TBK1 inhibitor GSK8612 (S8872, Selleck Chemicals), ULK1/2 inhibitor (MRT68921, BLDpharm), Vps34-IN1 inhibitor (APE-B6179, ApexBio), CK2 kinase inhibitor (CX4945, Selleckchem), Deferiprone (379409, Sigma Aldrich), oligomycin A (A5588, ApexBio), Antimycin A1 (A8674, Sigma-Aldrich), Q-VD-OPh (A1901, ApexBio), and DMSO (D2438, Sigma). The following siRNAs were used in this study: FIP200 (SMARTPOOL; LQ-021117-00-0002), ATG13 (SMARTPOOL; L-020765-01-0005), ULK1 (SMARTPOOL; L-005049-00-0005), ATG13 (SMARTPOOL; L-020765-01-0005), and non-targeting control pool (D-001810-10).

### Plasmid Construction

The sequences of all cDNAs were obtained by amplifying from existing plasmids, HAP1 cDNA, or gene synthesis (Genscript). For insect cell expressions, the sequences were codon optimized and gene synthesized (Genscript). Plasmids were generated by Gibson cloning. Inserts and vector backbones were generated by PCR amplification or excised from agarose gels after restriction enzyme digestion at 37°C for two hours. The inserts and plasmid backbones were purified with Promega Wizard SV gel and PCR Cleanup System (Promega). Purified inserts and backbones were mixed in a molar 3:1 ratio, respectively, supplemented by a 2x NEBuilder HiFi DNA assembly enzyme mix (New England Biolabs). Gibson reactions were incubated for one hour at 50°C and then transformed into DH5-alpha competent *E. coli* cells (ThermoFisher Cat#18265017). Transformed Gibson reactions were grown overnight on agar plates containing the appropriate selection marker (ampicillin, kanamycin, or chloramphenicol). Single colonies were picked, grown overnight in liquid cultures, and pelleted for DNA plasmid extraction using the GeneJet Plasmid Miniprep kit (Thermo Fisher). The purified plasmid DNA was submitted for DNA Sanger sequencing (MicroSynth AG). All insert sequences were verified by Sanger sequencing. Positive clones were further analyzed by whole plasmid sequencing (Plasmidsaurus). A detailed protocol is available (https://doi.org/10.17504/protocols.io.8epv5x11ng1b/v1).

### Cell lines

Cell lines were cultured at 37°C in humidified 5% C0_2_ atmosphere. HeLa (RRID:CVCL_0058) and HEK293T (RRID:CVCL_0063) cells were acquired from the American Type Culture Collection (ATCC). HeLa BNIP3/NIX double knockout clone #6 (RRID:CVCL_E1HA) and clone #10 (RRID:CVCL_E1HB), were generated with CRISPR/Cas9. HeLa and HEK293T cells were grown in Dulbecco Modified Eagle Medium (DMEM, Thermo Fisher) supplemented with 10% (v/v) Fetal Bovine Serum (FBS, Thermo Fisher), 25 mM HEPES (15630080, Thermo Fisher), 1% (v/v) non-essential amino acids (NEAA, 11140050, Thermo Fisher), and 1% (v/v) Penicillin-Streptomycin (15140122, Thermo Fisher). HAP1 cells were cultured in Iscove’s modified Dulbecco’s medium (Thermo Fisher) supplemented with 10% (v/v) FBS (Thermo Fisher) and 1% (v/v) penicillin–streptomycin (15140122, Thermo Fisher). All cell lines were tested regularly for mycoplasma contaminations. A detailed protocol is available (https://doi.org/10.17504/protocols.io.n2bvj3y5blk5/v1).

### Generation of CRISPR/Cas9 knockout cells

Knockout cell lines were generated using CRISPR/Cas9. Candidate single-guide RNAs (sgRNAs) were identified using CRISPick (RRID:SCR_025148; https://portals.broadinstitute.org/gppx/crispick/public), targeting all common splicing variants. The sgRNAs were ordered as short oligonucleotides (Microsynth) and cloned into pSpCas9(BB)-2A-GFP vector (RRID:Addgene_48138). The successful insertion of the sgRNAs was verified by Sanger sequencing. A detailed description of this cloning is available (https://doi.org/10.17504/protocols.io.j8nlkkzo6l5r/v1).

Plasmids containing a sgRNA were transfected into HeLa cells with Lipofectamine 3000 (Thermo Fisher). After 48 h, single GFP-positive cells were sorted by fluorescence-activated cell sorting (FACS) into 96 well plates. Single-cell colonies were expanded and positive clones were identified clones by immunoblotting. Candidate knockout clones with loss of protein expression for the target of interest were further analyzed by Sanger sequencing of the respective genomic regions. After DNA extraction, the regions of interest surrounding the sgRNA target sequence were amplified by PCR and analyzed by Sanger sequencing. The DNA sequences were compared to sequences from the parental line, and the edits were identified using the Synthego ICE v2 CRISPR Analysis Tool (https://www.synthego.com/products/bioinformatics/crispr-analysis). For NIX and BNIP3 double knockout cells or ATG13 single knockout cells, we transfected sgRNAs for the respective target genes into naïve HeLa cells (RRID:CVCL_0058) to obtain BNIP3/NIX double knockout cells #6 (RRID:CVCL_E1HA) and #10 (RRID: CVCL_E1HB) or ATG13 knockout cells #1 (RRID:CVCL_CVCL_E1HE). A detailed protocol is available (https://doi.org/10.17504/protocols.io.8epv59yx5g1b/v1).

### Generation of stable cell lines

Stable cell lines were generated using lentiviral or retroviral expression systems. For retroviral transductions, HEK293T cells (RRID:CVCL_0063) were transfected with VSV-G (a kind gift from Richard Youle), Gag-Pol (a kind gift from Richard Youle), and pBMN constructs containing our gene-of-interest using Lipofectamine 3000 (L3000008, Thermo Fisher). The next day, the medium was exchanged with fresh media. Viruses were harvested 48 h and 72 h after transfection. The retrovirus-containing supernatant was collected and filtered to avoid cross-over of HEK293T cells into the destination HeLa cells. After seeding HeLa cells at a density of 800k per well, cells were infected by the retrovirus-containing supernatant in the presence of 8 mg/ml polybrene (Sigma-Aldrich) for 24 h. The infected HeLa cells were expanded, and 10 days after infection, they were sorted by FACS to match equal expression levels where possible. A detailed protocol is available (https://doi.org/10.17504/protocols.io.81wgbyez1vpk/v1). The following retroviral vectors were used in this study: pCHAC-mito-mKeima (RRID:Addgene_72342).

For lentiviral transductions, HEK293T cells (RRID:CVCL_0063) were transfected with VSV-G, Gag-Pol, and pHAGE or pGenLenti constructs containing our gene-of-interest using Lipofectamine 3000 (L3000008, Thermo Fisher). The next day, the medium was exchanged with fresh media. Viruses were harvested 48 h and 72 h after transfection. The lentivirus-containing supernatant was collected and filtered to avoid cross-over of HEK293T cells into the HeLa cultures. After seeding HeLa cells at a density of 800k per well, cells were infected by the lentivirus-containing supernatant in the presence of 8 mg/ml polybrene (Sigma) for 24 h. The infected HeLa cells were expanded, and 10 days after infection, they were used for experiments. A detailed protocol is available (https://doi.org/10.17504/protocols.io.6qpvr3e5pvmk/v1). The following lentiviral vectors were used in this study: pHAGE-FKBP-GFP-WIPI1 (RRID:Addgene_223767), pHAGE-FKBP-GFP-WIPI2 (RRID:Addgene_223757), pHAGE-FKBP-GFP-WIPI3 (RRID:Addgene_223768), pHAGE-FKBP-GFP-WIPI4 (RRID:Addgene_223769), pHAGE-FKBP-GFP-WIPI2 R108E/R125E (RRID:Addgene_223770), pHAGE-FKBP-GFP-WIPI2 IDR (364-425aa) (RRID:Addgene_223758), pHAGE-mt-mKeima-P2A-FRB-Fis1 (RRID:Addgene_135295), pGenLenti V5-BNIP3 (RRID:Addgene_223732), pGenLenti V5-NIX (RRID:Addgene_223731), pGenLenti V5-NIX W36A/L39A (ΔLIR) (RRID:Addgene_223788), pGenLenti V5-NIX E72A/L75A/D77A/E81A (4A mutant; ΔWIPI2) (RRID:Addgene_223789), pGenLenti ATG13 (WT) (RRID: Addgene_ 223771), pGenLenti ATG13 (delta 191-230) (RRID: Addgene_223772), pGenLenti ATG13 (delta 191-205) (RRID: Addgene_223773), pGenLenti ATG13 (delta 206-230) (RRID: Addgene_223774).

### Mitophagy experiments

To induce BNIP3/NIX-mitophagy, cells were treated for 24 h with 1 mM Deferiprone (DFP) (379409, Sigma Aldrich), an iron chelator that mimics hypoxic conditions through stabilization of the transcription factor HIF1α and subsequent upregulation of NIX and BNIP3. Samples were analyzed by flow cytometry. A detailed protocol is available (https://doi.org/10.17504/protocols.io.e6nvw11m9lmk/v1). To induce PINK1/Parkin-mitophagy, cells were treated with 10 μM oligomycin (A5588, ApexBio) and 4 μM antimycin A (A8674, Sigma-Aldrich). In case cells were treated for more than 8 h, we also added 10 μM Q-VD-OPh (A1901, ApexBio) to suppress apoptosis. Samples were then analyzed by SDS– PAGE and western blot or flow cytometry. A detailed protocol is available (https://doi.org/10.17504/protocols.io.n2bvj3yjnlk5/v1).

### Non-selective autophagy experiments

To induce non-selective bulk autophagy, cells were starved by culturing them in Earle’s balanced salt medium (Cat# E3024, Sigma-Aldrich). Cells were collected and analyzed by SDS–PAGE and western blot analysis. A detailed protocol is available (https://doi.org/10.17504/protocols.io.4r3l228b3l1y/v1).

### Rapalog-induced chemical dimerization experiments

The chemical-induced dimerization (CID) experiments were performed using the FRB-Fis1 and FKBP fused to our gene of interest system. After consecutive lentiviral transduction of HeLa cells with both constructs, in which the FRB-Fis1 also expresses mitochondrially targeted monoKeima (mt-mKeima), cells were treated with 500 nM Rapalog A/C hetero-dimerizer rapalog (635057, Takara) for 24 h. Cells were then analyzed by flow cytometry. A detailed protocol is available (https://doi.org/10.17504/protocols.io.n92ldmyynl5b/v1).

### Flow cytometry

For mitochondrial flux experiments, 700K cells were seeded in 6 well plates one day before the experiment. Mitophagy was induced by treating the cells for the indicated times with deferiprone (DFP) or oligomycin A plus antimycin A1 (O/A), as described above. Cells were collected by removing the medium, washing the cells with 1x PBS (14190169, Thermo Fisher), trypsinization (T3924, Sigma), and resuspending in complete DMEM medium (41966052, Thermo Fisher). Filtered through 35 µm cell-strainer caps (352235, Falcon) and analyzed by an LSR Fortessa Cell Analyzer (BD Biosciences). Lysosomal mt-mKeima was measured using dual excitation ratiometric pH measurements at 405 (pH 7) and 561 (pH 4) nm lasers with 710/50-nm and 610/20-nm detection filters, respectively. Additional channels used for fluorescence compensation was GFP. Single fluorescence vector expressing cells were prepared to adjust photomultiplier tube voltages to make sure the signal was within detection limits, and to calculate the compensation matrix in BD FACSDiva Software. Depending on the experiment, we gated for GFP-positive and/or mKeima-positive cells with the appropriate compensation. For each sample, 10,000 mKeima-positive events were collected, and data were analyzed in FlowJo (RRID:SCR_008520; version 10.9.0; https://www.flowjo.com/solutions/flowjo). Our protocol was based on the previously described protocol (https://doi.org/10.17504/protocols.io.q26g74e1qgwz/v1).

For Rapalog-induced mitophagy experiments, cells were seeded as described above and treated for 24 h with 500 nM Rapalog A/C hetero-dimerizer (Takara). Cells were collected as described above, and the mt-mKeima ratio (561/405) was quantified by an LSR Fortessa Cell Analyzer (BD Biosciences). The cells were gated for GFP/mt-mKeima double-positive cells. Data were analyzed using FlowJo (version 10.9.0). A detailed protocol is available (https://doi.org/10.17504/protocols.io.n92ldmyynl5b/v1).

### SDS-PAGE and western blot analysis

For SDS-PAGE and western blot analysis, we collected cells by trypsinization and subsequent centrifugation at 300*g* for 5 min at 4°C. Cell pellets were washed in PBS and centrifuged once more at 300*g* for 5 min at 4°C. The supernatant was removed and the cell pellets were lysed in RIPA buffer (50 mM Tris-HCl pH 8.0, 150 mM NaCl, 0.5% sodium deoxycholate, 0.1% SDS, 1% NP-40) supplemented by cOmplete EDTA-free protease inhibitors (11836170001, Roche) and phosphatase inhibitors (Phospho-STOP, 4906837001, Roche). After incubating in RIPA buffer for 20 min on ice, samples were cleared by centrifugation at 20,000*g* for 10 min at 4°C. The soluble supernatant fraction was collected and protein concentrations were measured using the Pierce Detergent Compatible Bradford Assay Kit (23246, Thermo Fisher). Samples were then adjusted for equal loading and mixed with 6x protein loading dye, supplemented with 100 mM DTT, and boiled for 5 min at 95°C. Samples were loaded on 4-12% SDS-PAGE gels (NP0321BOX, NP0322BOX, or NP0323BOX, Thermo Fisher) with PageRuler Prestained protein marker (Thermo Fisher). Proteins were transferred onto nitrocellulose membranes (RPN132D, GE Healthcare) for 1 h at 4°C using the Mini Trans-Blot Cell (Bio-Rad). After the transfer, membranes were blocked with 5% milk powder dissolved in PBS-Tween (0.1% Tween 20) for 1 h at room temperature. The membranes were incubated overnight at 4°C with primary antibodies dissolved in the blocking buffer, washed three times for 5 min, and incubated with species-matched secondary horseradish peroxidase (HRP)-coupled antibodies diluted 1:10,000 in blocking buffer for 1 h at room temperature. Membranes were afterward washed three times with PBS-T and processed further for western blot detection. Membranes were incubated with SuperSignal West Femto Maximum Sensitivity Substrate (34096, Thermo Fisher) and imaged with a ChemiDoc MP Imaging system (Bio-Rad). Images were analyzed with ImageJ ^72^ (RRID:SCR_003070; https://imagej.net/). A detailed protocol is available (https://doi.org/10.17504/protocols.io.eq2lyj33plx9/v1). The primary antibodies used in this study are: anti-α-Tubulin (1:5000, Abcam Cat# ab7291, RRID:AB_2241126), anti-ATG13 (Cell Signaling Technology Cat# 13468, RRID:AB_2797419), anti-Beclin1 (1:1000, Cell Signaling Technology Cat# 3738, RRID:AB_490837), anti-phospho-Beclin1 Ser30 (1:1000, Cell Signaling Technology Cat# 54101, RRID:AB_3102019), anti-BNIP3 (1:1000, Cell Signaling Technology Cat# 44060, RRID:AB_2799259), anti-COXII (1:1000, Abcam Cat# ab110258, RRID:AB_10887758) or (1:1000, Cell Signaling Technology Cat# 31219, RRID:AB_2936222), anti-FIP200 (1:1000, Cell Signaling Technology Cat# 12436, RRID:AB_2797913), anti-LC3B (1:500, Nanotools Cat# 0260-100/LC3-2G6, RRID:AB_2943418), anti-BNIP3/NIXL (1:1000, Cell Signaling Technology Cat# 12396, RRID:AB_2688036), anti-OPTN (1:500, Sigma Aldrich Cat# HPA003279, RRID:AB_1079527), anti-phospho-OPTN Ser177 (1:1000, Cell Signaling Technology Cat# 57548, RRID:AB_2799529), anti-p62/SQSTM1 (1:1000, Abnova Cat# H00008878-M01, RRID:AB_437085), anti-phospho-p62/SQSTM1 Ser403 (1:1000, Cell Signaling Technology Cat# 39786, RRID:AB_2799162), anti-ULK1 (1:1000, Cell Signaling Technology Cat# 8054, RRID:AB_11178668). The secondary antibodies used in this study are: HRP conjugated polyclonal goat anti-mouse (Jackson ImmunoResearch Labs Cat# 115-035-003, RRID:AB_10015289), HRP conjugated polyclonal goat anti-rabbit (Jackson ImmunoResearch Labs Cat# 111-035-003, RRID:AB_2313567).

### Immunofluorescence and confocal microscopy

Cells were seeded on glass coverslips (12 mm #1.5) at a concentration of 100.000 cells/well, and after treatment with Rapalog for the indicated time, fixed in 4% paraformaldehyde (28906, Thermo Fisher Scientific) for 10 min at room temperature. After washing with PBS, cells were permeabilized with 0.1% (v/v) Triton X-100 (9002-93-1, Sigma-Aldrich) in PBS for 5 min. Blocking was performed with 5% (v/v) BSA (9048-46-8, Sigma-Aldrich) diluted in PBS for 1 h at room temperature. Primary and secondary antibodies were diluted in 5% BSA and incubated for 1 h at room temperature with three PBS washing steps in between. Cells were mounted on microscopy slides in DAPI Fluoromount-G mounting medium (0100-20, Southern Biotech), which stains the nuclei, and stored at 4 °C until use. Confocal microscopy was performed with a Zeiss LSM700 laser scanning confocal microscopy with Plan-Apochromat 40×/1.30 Oil DIC, WD 0.21 mm objective. A detailed protocol is available (https://doi.org/10.17504/protocols.io.6qpvr8p1olmk/v1). The primary antibodies used in this study are: anti-ATG13 (1:200, Cell Signaling Technology, Cat# 13468; RRID:AB_2797419). The secondary antibodies used in this study are: AlexaFluor-546 goat anti-rabbit IgG (H+L) (1:500, Thermo Fisher, Cat# A-11035; RRID: AB_2534093).

### Purification ATG9A-vesicles

HAP1 cells were CRISPR-edited to introduce a C-terminal GFP-TEV-Flag tag into the endogenous locus of ATG9A (RRID:CVCL_E2TR). For the isolation of native ATG9A-vesicles, five 15 cm dishes were seeded. Cells were collected by trypsinization and centrifugation. After washing with PBS, cell pellets were flash-frozen and stored at 80°C until use. For vesicle isolation, cell pellets were thawed on ice and resuspended in 1665 µl of Vesicle Isolation Buffer (20 mM HEPES pH 7.5, 150 mM NaCl, 250 mM sucrose, 1x cOmplete EDTA-free protease inhibitors (Roche), 20 mM β-Glycerophosphate, 1 mM Sodium Orthovanadate, 1 mM NaF, and 1 mM EDTA pH 8.0). The cell suspension was transferred to a 2 ml microcentrifuge tube and, after incubation on ice for 20 minutes, lysed by passing the suspension through a 26G needle 30 times, chilling on ice for 10 minutes, followed by another 30 passes through the needle. The lysate was centrifuged at 2000*g* for 6 minutes at 4°C to pellet cell debris and nuclei. The supernatant was collected and 70 µl of pre-equilibrated FLAG beads were added. The mixture was incubated overnight at 4°C on a roller. On the second day, beads were pelleted at 2000*g* for 3 minutes at 4°C, and the unbound supernatant was removed. Beads were resuspended in 1 ml of vesicle isolation buffer, transferred to a fresh 2 ml microcentrifuge tube containing 665 µl of Vesicle Isolation Buffer (VIB). In the second washing step, the beads were resuspended with 1 ml VIB and subsequently 600 µl of Elution Buffer (20 mM HEPES pH 7.5, 150 mM NaCl, 1x cOmplete EDTA-free protease inhibitors (Roche), 20 mM β-Glycerophosphate, 1 mM Sodium Orthovanadate, and 1 mM NaF) was added slowly. For the third wash 1.6 mL Elution Buffer was used. After the third wash the beads were resuspended in 665 µl Elution Buffer and transferred to a 1.5 ml LoBind tube. To elute the ATG9A-vesicles from the beads, 16.65 µl of 4 mg/ml FLAG peptide (Sigma, F32920-4MG) was added to the suspension and incubated for 3 hours at 4°C while rolling. The supernatant was collected after pelleting the beads at 2000*g* for 3 minutes and used for experiments. All procedures involving the handling of cells and vesicles were performed on ice to maintain sample integrity. A detailed protocol is available (https://doi.org/10.17504/protocols.io.dm6gpzok8lzp/v1).

### Protein expression and purification

To purify NIX-GST, the cytosol-exposed domain of NIX (1-182aa) was fused to a C-terminal GST-tag through cloning into a pET-DUET1 vector (RRID:Addgene_223733). Point mutants were introduced by *in vitro* mutagenesis to generate NIX E72A/L75A/D77A/E81A (4A; ΔWIPI2) (RRID:Addgene_223753), and NIX W36A/L39A (ΔLIR) (RRID:Addgene_223738).

After the transformation of the pET-DUET1 vector encoding NIX-GST wild-type or mutants in *E. coli* Rosetta pLysS cells (Novagen Cat# 70956-4), cells were grown in 2x Tryptone Yeast extract (TY) medium at 37°C until an OD_600_ of 0.4 and then continued at 18°C. Once the cells reached an OD_600_ of 0.8, protein expression was induced with 100 µM isopropyl β-D-1-thiogalactopyranoside (IPTG) for 16 h at 18°C. Cells were collected by centrifugation and resuspended in lysis buffer (50 mM Tris-HCl pH 7.4, 300 mM NaCl, 1% Triton X-100, 5% glycerol, 2 mM MgCl_2_, 1 mM DTT, 2mM β-mercaptoethanol, cOmplete EDTA-free protease inhibitors (Roche), CIP protease inhibitor (Sigma), and DNase (Sigma)). Cell lysates were sonicated twice for 30 s and cleared by centrifugation at 18,000 rpm for 45 min at 4°C in a SORVAL RC6+ centrifuge with an F21S-8x50Y rotor (Thermo Scientific). The supernatant was collected and incubated with pre-equilibrated Glutathione Sepharose 4B beads (GE Healthcare) for 2 h at 4°C with gentle shaking to bind NIX-GST. Samples were centrifuged to pellet the beads and remove the unbound lysate. Beads were then washed twice with wash buffer (50 mM Tris-HCl pH 7.4, 300 mM NaCl, 1 mM DTT), once with high salt wash buffer (50 mM Tris-HCl pH 7.4, 700 mM NaCl, 1 mM DTT), and two more times with wash buffer (50 mM Tris-HCl pH 7.4, 300 mM NaCl, 1 mM DTT). Beads were incubated overnight with 4 ml of 50 mM reduced glutathione dissolved in wash buffer (50 mM Tris-HCl pH 7.4, 300 mM NaCl, 1 mM DTT) at 4°C, to elute NIX-GST from the beads. To collect the supernatant, the beads were collected by centrifugation. The beads were washed twice with 4 ml of wash buffer, and the supernatant was collected. The supernatant fractions were pooled, filtered through a 0.45 µm syringe filter, concentrated with 10 kDa cut-off Amicon filter (Merck Millipore), and loaded onto a pre-equilibrated Superdex 200 Increase 10/300 GL column (Cytiva). Proteins were eluted with SEC buffer (25 mM Tris-HCl pH 7.4, 300 mM NaCl, 1 mM DTT). Fractions were analyzed by SDS-PAGE and Coomassie staining. Fractions containing purified NIX-GST were pooled. After concentrating the purified protein, the protein was aliquoted and snap-frozen in liquid nitrogen. Proteins were stored at -80°C. A detailed protocol is available (https://doi.org/10.17504/protocols.io.q26g711dkgwz/v1).

To purify BNIP3-GST, we purchased the gene-synthesized codon-optimized cytosol-exposed domain of BNIP3 (1-158aa) fused to a C-terminal GST-tag in a pFastBac-Dual vector from Genscript (RRID:Addgene_223764). Point mutants were introduced by *in vitro* mutagenesis to generate BNIP3 E44A/L47A/D49A/A50K/Q51A (5A; ΔWIPI2) (RRID:Addgene_223777), and BNIP3 W18A/L21A (ΔLIR) (RRID:Addgene_223778). The constructs were used to generate bacmid DNA, using the Bac-to-Bac system, by amplification in DH10BacY cells ^73^. After the bacmid DNA was verified by PCR for insertion of the transgene, we purified bacmid DNA for transfection into Sf9 insect cells (12659017, Thermo Fisher, RRID:CVCL_0549). To this end, we mixed 2500 ng of plasmid DNA with FuGene transfection reagent (Promega) and transfected 1 million Sf9 cells seeded in a 6 well plate. About 7 days after transfection, the V0 virus was harvested and used to infect 40 ml of 1 million cells per ml of Sf9 cells. The viability of the cultures was closely monitored and upon the decrease in viability and confirmation of yellow fluorescence, we collected the supernatant after centrifugation and stored this as V1 virus. For expressions, we infected 1 L of Sf9 cells, at 1 million cells per ml, with 1 ml of V1 virus. When the viability of the cells decreased to 90-95%, cells were collected by centrifugation. Cell pellets were washed with 1x PBS and flash-frozen in liquid nitrogen. Pellets were stored at -80°C. For purification of BNIP3-GST wild-type or mutants, pellets were resuspended in 25 ml lysis buffer (50 mM Tris-HCl pH 7.4, 300 mM NaCl, 1 mM DTT, 2 mM MgCl_2_, 2 mM β-mercaptoethanol, 1% Triton X-100, 5% glycerol, 1 µl benzonase (Sigma), cOmplete EDTA-free protease inhibitors (Roche), CIP protease inhibitor (Sigma)). Cells were homogenized with a douncer and cell lysates were cleared by centrifugation at 18,000 rpm for 45 min at 4°C in a SORVAL RC6+ centrifuge with an F21S-8x50Y rotor (Thermo Scientific). The supernatant was collected and incubated with pre-equilibrated Glutathione Sepharose 4B beads (GE Healthcare) for 2 h at 4°C with gentle shaking to bind BNIP3-GST. Samples were centrifuged to pellet the beads and remove the unbound lysate. Beads were then washed twice with wash buffer (50 mM Tris-HCl pH 7.4, 300 mM NaCl, 1 mM DTT), once with high salt wash buffer (50 mM Tris-HCl pH 7.4, 700 mM NaCl, 1 mM DTT), and two more times with wash buffer (50 mM Tris-HCl pH 7.4, 300 mM NaCl, 1 mM DTT). Beads were incubated overnight with 4 ml of 50 mM reduced glutathione dissolved in wash buffer (50 mM Tris-HCl pH 7.4, 300 mM NaCl, 1 mM DTT) at 4°C, to elute BNIP3-GST from the beads. To collect the supernatant, the beads were collected by centrifugation. The beads were washed twice with 4 ml of wash buffer, and the supernatant was collected. The supernatant fractions were pooled, filtered through a 0.45 µm syringe filter, concentrated with 10 kDa cut-off Amicon filter (Merck Millipore), and loaded onto a pre-equilibrated Superdex 200 Increase 10/300 GL column (Cytiva). Proteins were eluted with SEC buffer (25 mM Tris-HCl pH 7.4, 300 mM NaCl, 1 mM DTT). Fractions were analyzed by SDS-PAGE and Coomassie staining. Fractions containing purified BNIP3-GST were pooled. After concentrating the purified protein, the protein was aliquoted and snap-frozen in liquid nitrogen. Proteins were stored at -80°C. A detailed protocol is available (https://doi.org/10.17504/protocols.io.261ge5527g47/v1).

To purify FUNDC1-GST, the cytosol-exposed domain of FUNDC1 (1-50aa) was fused to a C-terminal GST-tag through cloning into a pET-DUET1 vector (RRID:Addgene_223734). Point mutants were introduced by *in vitro* mutagenesis to generate FUNDC1 Y18A/L21A (ΔLIR) (RRID:Addgene_223739). After the transformation of the pET-DUET1 vector encoding FUNDC1-GST wild-type or mutants in *E. coli* Rosetta pLysS cells (Novagen Cat# 70956-4), cells were grown in 2x Tryptone Yeast extract (TY) medium at 37°C until an OD_600_ of 0.4 and then continued at 18°C. Once the cells reached an OD_600_ of 0.8, protein expression was induced with 100 µM isopropyl β-D-1-thiogalactopyranoside (IPTG) for 16 h at 18°C. Cells were collected by centrifugation and resuspended in lysis buffer (50 mM Tris-HCl pH 7.4, 300 mM NaCl, 1% Triton X-100, 5% glycerol, 2 mM MgCl_2_, 1 mM DTT, 2mM β-mercaptoethanol, cOmplete EDTA-free protease inhibitors (Roche), CIP protease inhibitor (Sigma), and DNase (Sigma)). Cell lysates were sonicated twice for 30 s and cleared by centrifugation at 18,000 rpm for 45 min at 4°C in a SORVAL RC6+ centrifuge with an F21S-8x50Y rotor (Thermo Scientific). The supernatant was collected and incubated with pre-equilibrated Glutathione Sepharose 4B beads (GE Healthcare) for 2 h at 4°C with gentle shaking to bind FUNDC1-GST. Samples were centrifuged to pellet the beads and remove the unbound lysate. Beads were then washed twice with wash buffer (50 mM Tris-HCl pH 7.4, 300 mM NaCl, 1 mM DTT), once with high salt wash buffer (50 mM Tris-HCl pH 7.4, 700 mM NaCl, 1 mM DTT), and two more times with wash buffer (50 mM Tris-HCl pH 7.4, 300 mM NaCl, 1 mM DTT). Beads were incubated overnight with 4 ml of 50 mM reduced glutathione dissolved in wash buffer (50 mM Tris-HCl pH 7.4, 300 mM NaCl, 1 mM DTT) at 4°C, to elute FUNDC1-GST from the beads. To collect the supernatant, the beads were collected by centrifugation. The beads were washed twice with 4 ml of wash buffer, and the supernatant was collected. The supernatant fractions were pooled, filtered through a 0.45 µm syringe filter, concentrated with 10 kDa cut-off Amicon filter (Merck Millipore), and loaded onto a pre-equilibrated Superdex 200 Increase 10/300 GL column (Cytiva). Proteins were eluted with SEC buffer (25 mM Tris-HCl pH 7.4, 300 mM NaCl, 1 mM DTT). Fractions were analyzed by SDS-PAGE and Coomassie staining. Fractions containing purified FUNDC1-GST were pooled. After concentrating the purified protein, the protein was aliquoted and snap-frozen in liquid nitrogen. Proteins were stored at -80°C. A detailed protocol is available (https://doi.org/10.17504/protocols.io.14egn66nyl5d/v1).

To purify BCL2L13-GST, the cytosol-exposed domain of BCL2L13 (1-465aa) was fused to a C-terminal GST-tag through cloning into a pET-DUET1 vector (RRID:Addgene_223744). Point mutants were introduced by *in vitro* mutagenesis to generate BCL2L13 W276A/I279A (ΔLIR1) (RRID:Addgene_223749), BCL2L13 Y213A/I216A/W276A/I279A (ΔLIR1+2) (RRID:Addgene_223752), BCL2L13 I224A/L227A/W276A/I279A (ΔLIR1+3) (RRID:Addgene_223754), BCL2L13 W276A/I279A/I307A/V310A (ΔLIR1+4) (RRID:Addgene_223755), BCL2L13 I224A/L227A/W276A/I279A/I307A/V310A (ΔLIR1+3+4) (RRID:Addgene_223756). After the transformation of the pET-DUET1 vector encoding BCL2L13-GST wild-type or mutants in *E. coli* Rosetta pLysS cells (Novagen Cat# 70956-4), cells were grown in 2x Tryptone Yeast extract (TY) medium at 37°C until an OD_600_ of 0.4 and then continued at 18°C. Once the cells reached an OD_600_ of 0.8, protein expression was induced with 100 µM isopropyl β-D-1-thiogalactopyranoside (IPTG) for 16 h at 18°C. Cells were collected by centrifugation and resuspended in lysis buffer (50 mM Tris-HCl pH 7.4, 300 mM NaCl, 1% Triton X-100, 5% glycerol, 2 mM MgCl_2_, 1 mM DTT, 2mM β-mercaptoethanol, cOmplete EDTA-free protease inhibitors (Roche), CIP protease inhibitor (Sigma), and DNase (Sigma)). Cell lysates were sonicated twice for 30 s and cleared by centrifugation at 18,000 rpm for 45 min at 4°C in a SORVAL RC6+ centrifuge with an F21S-8x50Y rotor (Thermo Scientific). The supernatant was collected and incubated with pre-equilibrated Glutathione Sepharose 4B beads (GE Healthcare) for 2 h at 4°C with gentle shaking to bind BCL2L13-GST. Samples were centrifuged to pellet the beads and remove the unbound lysate. Beads were then washed twice with wash buffer (50 mM Tris-HCl pH 7.4, 300 mM NaCl, 1 mM DTT), once with high salt wash buffer (50 mM Tris-HCl pH 7.4, 700 mM NaCl, 1 mM DTT), and two more times with wash buffer (50 mM Tris-HCl pH 7.4, 300 mM NaCl, 1 mM DTT). Beads were incubated overnight with 4 ml of 50 mM reduced glutathione dissolved in wash buffer (50 mM Tris-HCl pH 7.4, 300 mM NaCl, 1 mM DTT) at 4°C, to elute BCL2L13-GST from the beads. To collect the supernatant, the beads were collected by centrifugation. The beads were washed twice with 4 ml of wash buffer, and the supernatant was collected. The supernatant fractions were pooled, filtered through a 0.45 µm syringe filter, concentrated with 10 kDa cut-off Amicon filter (Merck Millipore), and loaded onto a pre-equilibrated Superdex 200 Increase 10/300 GL column (Cytiva). Proteins were eluted with SEC buffer (25 mM Tris-HCl pH 7.4, 300 mM NaCl, 1 mM DTT). Fractions were analyzed by SDS-PAGE and Coomassie staining. Fractions containing purified BCL2L13-GST were pooled. After concentrating the purified protein, the protein was aliquoted and snap-frozen in liquid nitrogen. Proteins were stored at -80°C. A detailed protocol is available (https://doi.org/10.17504/protocols.io.rm7vzjj12lx1/v1).

To purify GFP-tagged NIX-GFP (RRID:Addgene_223736), NIX(W36A/L39A)-GFP (ΔLIR) (RRID:Addgene_223748), BNIP3-GFP (RRID:Addgene_223765), BNIP3(W18A/L21A)-GFP (ΔLIR) (RRID:Addgene_223766), BCL2L13-GFP (RRID:Addgene_223745), BCL2L13(W276A/I279A)-GFP (ΔLIR1) (RRID:Addgene_223746), BCL2L13(Y213A/I216A)-GFP (ΔLIR2) (RRID:Addgene_223783), BCL2L13(I224A/L227A)-GFP (ΔLIR3) (RRID:Addgene_223775), BCL2L13(I307A/V310A)-GFP (ΔLIR4) (RRID:Addgene_223776), BCL2L13(Y213A/I216A/W276A/I279A)-GFP (ΔLIR1+2) (RRID:Addgene_223782), BCL2L13(I224A/L227A/W276A/I279A)-GFP (ΔLIR1+3) (RRID:Addgene_223780), BCL2L13(W276A/I279A/I307A/V310A)-GFP (ΔLIR1+4) (RRID:Addgene_223781), BCL2L13(I224A/L227A/W276A/I279A/I307A/V310A)-GFP (ΔLIR1+3+4) (RRID:Addgene_ 223784), FUNDC1-GFP (RRID:Addgene_223737), FUNDC1(Y18A/L21A)-GFP (ΔLIR) (RRID:Addgene_223750), the same expression and purification methods were used as described above. However, as we introduced a TEV-cleavage site between the C-terminally GFP-tagged cargo receptor and the GST-tag (i.e. cargo receptor-GFP-TEV-GST), we cleaved off the GST-tag overnight by eluting the GFP-tagged cargo receptor from the GSH beads by the addition of TEV protease. The rest of the purification was the same as described above. Detailed protocols are available (https://doi.org/10.17504/protocols.io.x54v9219zl3e/v1), (https://doi.org/10.17504/protocols.io.kqdg328r7v25/v1), (https://doi.org/10.17504/protocols.io.4r3l2qnx3l1y/v1), (https://doi.org/10.17504/protocols.io.36wgqno2ogk5/v1).

To purify GST-TEX264, the cytosol-exposed domain of TEX264 (28-313aa) fused to a N-terminal GST-tag was gene synthesized by Genscript and cloned into a pGEX-4T1 vector. After the transformation of the pGEX-4T1 vector encoding GST-TEX264 in *E. coli* Rosetta pLysS cells (Novagen Cat# 70956-4), cells were grown in 2x Tryptone Yeast extract (TY) medium at 37°C until an OD_600_ of 0.4 and then continued at 18°C. Once the cells reached an OD_600_ of 0.8, protein expression was induced with 100 µM isopropyl β-D-1-thiogalactopyranoside (IPTG) for 16 h at 18°C. Cells were collected by centrifugation and resuspended in lysis buffer (50 mM Tris-HCl pH 7.4, 300 mM NaCl, 1% Triton X-100, 5% glycerol, 2 mM MgCl_2_, 1 mM DTT, 2mM β-mercaptoethanol, cOmplete EDTA-free protease inhibitors (Roche), CIP protease inhibitor (Sigma), and DNase (Sigma)). Cell lysates were sonicated twice for 30 s and cleared by centrifugation at 18,000 rpm for 45 min at 4°C in a SORVAL RC6+ centrifuge with an F21S-8x50Y rotor (Thermo Scientific). The supernatant was collected and incubated with pre-equilibrated Glutathione Sepharose 4B beads (GE Healthcare) for 2 h at 4°C with gentle shaking to bind GST-TEX264. Samples were centrifuged to pellet the beads and remove the unbound lysate. Beads were then washed twice with wash buffer (50 mM Tris-HCl pH 7.4, 300 mM NaCl, 1 mM DTT), once with high salt wash buffer (50 mM Tris-HCl pH 7.4, 700 mM NaCl, 1 mM DTT), and two more times with wash buffer (50 mM Tris-HCl pH 7.4, 300 mM NaCl, 1 mM DTT). Beads were incubated overnight with 4 ml of 50 mM reduced glutathione dissolved in wash buffer (50 mM Tris-HCl pH 7.4, 300 mM NaCl, 1 mM DTT) at 4°C, to elute GST-TEX264 from the beads. To collect the supernatant, the beads were collected by centrifugation. The beads were washed twice with 4 ml of wash buffer, and the supernatant was collected. The supernatant fractions were pooled, filtered through a 0.45 µm syringe filter, concentrated with 30 kDa cut-off Amicon filter (Merck Millipore), and loaded onto a pre-equilibrated Superdex 200 Increase 10/300 GL column (Cytiva). Proteins were eluted with SEC buffer (25 mM Tris-HCl pH 7.4, 150 mM NaCl, 1 mM DTT). Fractions were analyzed by SDS-PAGE and Coomassie staining. Fractions containing purified GST-TEX264 were pooled. After concentrating the purified protein, the protein was aliquoted and snap-frozen in liquid nitrogen. Proteins were stored at -80°C.

To purify GST-FAM134C, the cytosol-exposed domain of FAM134C (250-466aa) fused to a N-terminal GST-tag was gene synthesized by Genscript and cloned into a pGEX-4T1 vector. After the transformation of the pGEX-4T1 vector encoding GST-FAM134C in *E. coli* Rosetta pLysS cells (Novagen Cat# 70956-4), cells were grown in 2x Tryptone Yeast extract (TY) medium at 37°C until an OD_600_ of 0.4 and then continued at 18°C. Once the cells reached an OD_600_ of 0.8, protein expression was induced with 100 µM isopropyl β-D-1-thiogalactopyranoside (IPTG) for 16 h at 18°C. Cells were collected by centrifugation and resuspended in lysis buffer (50 mM Tris-HCl pH 7.4, 300 mM NaCl, 1% Triton X-100, 5% glycerol, 2 mM MgCl_2_, 1 mM DTT, 2mM β-mercaptoethanol, cOmplete EDTA-free protease inhibitors (Roche), CIP protease inhibitor (Sigma), and DNase (Sigma)). Cell lysates were sonicated twice for 30 s and cleared by centrifugation at 18,000 rpm for 45 min at 4°C in a SORVAL RC6+ centrifuge with an F21S-8x50Y rotor (Thermo Scientific). The supernatant was collected and incubated with pre-equilibrated Glutathione Sepharose 4B beads (GE Healthcare) for 2 h at 4°C with gentle shaking to bind GST-FAM134C. Samples were centrifuged to pellet the beads and remove the unbound lysate. Beads were then washed twice with wash buffer (50 mM Tris-HCl pH 7.4, 300 mM NaCl, 1 mM DTT), once with high salt wash buffer (50 mM Tris-HCl pH 7.4, 700 mM NaCl, 1 mM DTT), and two more times with wash buffer (50 mM Tris-HCl pH 7.4, 300 mM NaCl, 1 mM DTT). Beads were incubated overnight with 4 ml of 50 mM reduced glutathione dissolved in wash buffer (50 mM Tris-HCl pH 7.4, 300 mM NaCl, 1 mM DTT) at 4°C, to elute GST-FAM134C from the beads. To collect the supernatant, the beads were collected by centrifugation. The beads were washed twice with 4 ml of wash buffer, and the supernatant was collected. The supernatant fractions were pooled, filtered through a 0.45 µm syringe filter, concentrated with 30 kDa cut-off Amicon filter (Merck Millipore), and loaded onto a pre-equilibrated Superdex 200 Increase 10/300 GL column (Cytiva). Proteins were eluted with SEC buffer (25 mM Tris-HCl pH 7.4, 150 mM NaCl, 1 mM DTT). Fractions were analyzed by SDS-PAGE and Coomassie staining. Fractions containing purified GST-FAM134C were pooled. After concentrating the purified protein, the protein was aliquoted and snap-frozen in liquid nitrogen. Proteins were stored at -80°C.

To purify CCPG1-GST, the cytosol-exposed domain of CCPG1 (1-212aa) fused to a C-terminal GST-tag was gene synthesized by Genscript and cloned into a pET-DUET1 vector. After the transformation of the pET-DUET1 vector encoding CCPG1-GST in *E. coli* Rosetta pLysS cells (Novagen Cat# 70956-4), cells were grown in 2x Tryptone Yeast extract (TY) medium at 37°C until an OD_600_ of 0.4 and then continued at 18°C. Once the cells reached an OD_600_ of 0.8, protein expression was induced with 100 µM isopropyl β-D-1-thiogalactopyranoside (IPTG) for 16 h at 18°C. Cells were collected by centrifugation and resuspended in lysis buffer (50 mM Tris-HCl pH 7.4, 300 mM NaCl, 1% Triton X-100, 5% glycerol, 2 mM MgCl_2_, 1 mM DTT, 2mM β-mercaptoethanol, cOmplete EDTA-free protease inhibitors (Roche), CIP protease inhibitor (Sigma), and DNase (Sigma)). Cell lysates were sonicated twice for 30 s and cleared by centrifugation at 18,000 rpm for 45 min at 4°C in a SORVAL RC6+ centrifuge with an F21S-8x50Y rotor (Thermo Scientific). The supernatant was collected and incubated with pre-equilibrated Glutathione Sepharose 4B beads (GE Healthcare) for 2 h at 4°C with gentle shaking to bind CCPG1-GST. Samples were centrifuged to pellet the beads and remove the unbound lysate. Beads were then washed twice with wash buffer (50 mM Tris-HCl pH 7.4, 300 mM NaCl, 1 mM DTT), once with high salt wash buffer (50 mM Tris-HCl pH 7.4, 700 mM NaCl, 1 mM DTT), and two more times with wash buffer (50 mM Tris-HCl pH 7.4, 300 mM NaCl, 1 mM DTT). Beads were incubated overnight with 4 ml of 50 mM reduced glutathione dissolved in wash buffer (50 mM Tris-HCl pH 7.4, 300 mM NaCl, 1 mM DTT) at 4°C, to elute CCPG1-GST from the beads. To collect the supernatant, the beads were collected by centrifugation. The beads were washed twice with 4 ml of wash buffer, and the supernatant was collected. The supernatant fractions were pooled, filtered through a 0.45 µm syringe filter, concentrated with 30 kDa cut-off Amicon filter (Merck Millipore), and loaded onto a pre-equilibrated Superdex 200 Increase 10/300 GL column (Cytiva). Proteins were eluted with SEC buffer (25 mM Tris-HCl pH 7.4, 300 mM NaCl, 1 mM DTT). Fractions were analyzed by SDS-PAGE and Coomassie staining. Fractions containing purified CCPG1-GST were pooled. After concentrating the purified protein, the protein was aliquoted and snap-frozen in liquid nitrogen. Proteins were stored at -80°C. A detailed protocol is available (https://doi.org/10.17504/protocols.io.e6nvw14dzlmk/v1).

To purify FKBP8-GST, the cytosol-exposed domain of FKBP8 (1-391aa) fused to a C-terminal GST-tag was gene synthesized by Genscript and cloned into a pET-DUET1 vector. After the transformation of the pET-DUET1 vector encoding FKBP8-GST in *E. coli* Rosetta pLysS cells (Novagen Cat# 70956-4), cells were grown in 2x Tryptone Yeast extract (TY) medium at 37°C until an OD_600_ of 0.4 and then continued at 18°C. Once the cells reached an OD_600_ of 0.8, protein expression was induced with 100 µM isopropyl β-D-1-thiogalactopyranoside (IPTG) for 16 h at 18°C. Cells were collected by centrifugation and resuspended in lysis buffer (50 mM Tris-HCl pH 7.4, 300 mM NaCl, 1% Triton X-100, 5% glycerol, 2 mM MgCl_2_, 1 mM DTT, 2mM β-mercaptoethanol, cOmplete EDTA-free protease inhibitors (Roche), CIP protease inhibitor (Sigma), and DNase (Sigma)). Cell lysates were sonicated twice for 30 s and cleared by centrifugation at 18,000 rpm for 45 min at 4°C in a SORVAL RC6+ centrifuge with an F21S-8x50Y rotor (Thermo Scientific). The supernatant was collected and incubated with pre-equilibrated Glutathione Sepharose 4B beads (GE Healthcare) for 2 h at 4°C with gentle shaking to bind FKBP8-GST. Samples were centrifuged to pellet the beads and remove the unbound lysate. Beads were then washed twice with wash buffer (50 mM Tris-HCl pH 7.4, 300 mM NaCl, 1 mM DTT), once with high salt wash buffer (50 mM Tris-HCl pH 7.4, 700 mM NaCl, 1 mM DTT), and two more times with wash buffer (50 mM Tris-HCl pH 7.4, 300 mM NaCl, 1 mM DTT). Beads were incubated overnight with 4 ml of 50 mM reduced glutathione dissolved in wash buffer (50 mM Tris-HCl pH 7.4, 300 mM NaCl, 1 mM DTT) at 4°C, to elute FKBP8-GST from the beads. To collect the supernatant, the beads were collected by centrifugation. The beads were washed twice with 4 ml of wash buffer, and the supernatant was collected. The supernatant fractions were pooled, filtered through a 0.45 µm syringe filter, concentrated with 30 kDa cut-off Amicon filter (Merck Millipore), and loaded onto a pre-equilibrated Superdex 200 Increase 10/300 GL column (Cytiva). Proteins were eluted with SEC buffer (25 mM Tris-HCl pH 7.4, 300 mM NaCl, 1 mM DTT). Fractions were analyzed by SDS-PAGE and Coomassie staining. Fractions containing purified FKBP8-GST were pooled. After concentrating the purified protein, the protein was aliquoted and snap-frozen in liquid nitrogen. Proteins were stored at -80°C. A detailed protocol is available (https://doi.org/10.17504/protocols.io.n2bvjne3pgk5/v1).

To purify FIP200-GFP from insect cells, we purchased gene-synthesized codon-optimized GST-3C-FIP200-EGFP in a pGB-02-03 vector from Genscript (RRID:Addgene_187832). The V1 virus was generated as described above for BNIP3-GST. For expressions, we infected 1 L of Sf9 cells (12659017, Thermo Fisher, RRID:CVCL_0549), at 1 million cells per ml, with 1 ml of V1 virus. When the viability of the cells decreased to 90-95%, cells were collected by centrifugation. Cell pellets were washed with 1x PBS and flash-frozen in liquid nitrogen. Pellets were stored at -80°C. For purification of FIP200-GFP, the pellet was resuspended in 25 ml lysis buffer (50 mM HEPES pH 7.5, 300 mM NaCl, 1 mM MgCl_2_, 10% glycerol, 1mM DTT, 0.5% CHAPS, 1 µl benzonase (Sigma), cOmplete EDTA-free protease inhibitors (Roche), CIP protease inhibitor (Sigma)). Cells were homogenized with a douncer. Cell lysates were cleared by centrifugation at 72,000*g* for 45 min at 4°C with a Beckman Ti45 rotor. The supernatant was collected and incubated with pre-equilibrated Glutathione Sepharose 4B beads (GE Healthcare) for overnight at 4°C with gentle shaking to bind GST-3C-FIP200-EGFP. Samples were centrifuged to pellet the beads and remove the unbound lysate. Beads were washed seven times with wash buffer (50 mM HEPES pH 7.5, 200 mM NaCl, 1 mM MgCl_2_, 1 mM DTT). Beads were incubated overnight with precision 3C protease in wash buffer at 4°C. After the proteins were released from the beads by the 3C protease, the supernatant was collected after centrifugation of the beads. The beads were washed twice with 4 ml of wash buffer, and the supernatant was collected. The supernatant fractions were pooled, filtered through a 0.45 µm syringe filter, and concentrated with a 100 kDa cut-off Amicon filter (Merck Millipore). The proteins were loaded onto a pre-equilibrated Superose 6 Increase 10/300 GL column (Cytiva). Proteins were eluted with SEC buffer (25 mM HEPES pH 7.5, 200 mM NaCl, 1 mM DTT). Fractions were analyzed by SDS-PAGE and Coomassie staining. Fractions containing purified FIP200-GFP were pooled. After concentrating the purified protein, the protein was aliquoted and snap-frozen in liquid nitrogen. Proteins were stored at -80°C. A detailed protocol can be found here (https://doi.org/10.17504/protocols.io.dm6gpbkq5lzp/v1).

To purify GFP-FIP200 C-terminal region (CTR), as described previously ^4^, the C-terminal domain of FIP200 (1429-1591aa) was fused to a N-terminal 6xHis-TEV-GFP-tag through cloning into a pET-DUET1 vector (RRID:Addgene_223724). After the transformation of the pET-DUET1 vector encoding 6xHis-TEV-GFP-FIP200(CTR) in *E. coli* Rosetta pLysS cells (Novagen Cat# 70956-4), cells were grown in 2x Tryptone Yeast extract (TY) medium at 37°C until an OD_600_ of 0.4 and then continued at 18°C. Once the cells reached an OD_600_ of 0.8, protein expression was induced with 100 µM isopropyl β-D-1-thiogalactopyranoside (IPTG) for 16 h at 18°C. Cells were collected by centrifugation and resuspended in lysis buffer (50 mM Tris-HCl pH 7.4, 300 mM NaCl, 2 mM MgCl_2_, 5% glycerol, 10 mM Imidazole, 2 mM β-mercaptoethanol, cOmplete EDTA-free protease inhibitors (Roche), CIP protease inhibitor (Sigma), and DNase (Sigma)). Cell lysates were sonicated twice for 30 s. Lysates were cleared by centrifugation at 18,000 rpm for 45 min at 4°C in a SORVAL RC6+ centrifuge with an F21S-8x50Y rotor (Thermo Scientific). The supernatant was filtered through an 0.45 µm filter and loaded onto a pre-equilibrated 5 ml His-Trap HP column (Cytiva). After His tagged proteins were bound to the column, the column was washed with three column volumes of wash buffer (50 mM Tris-HCl pH 7.4, 300 mM NaCl, 10 mM Imidazole, 2 mM β-mercaptoethanol). Proteins were then eluted with a stepwise imidazole gradient (30, 75, 100, 150, 225, 300 mM). Fractions containing the 6xHis-TEV-GFP-FIP200(CTR) were pooled and incubated overnight with TEV protease at 4°C. After the 6xHis tag was cleaved off, 6xHis tag and His-tagged TEV protease was recaptured with nickel beads for 1 h at 4 degrees. The beads were pelleted by centrifugation and the supernatant, containing the GFP-FIP200(CTR) protein was concentrated using a 30 kDa cut-off Amicon filter (Merck Millipore) and loaded onto a pre-equilibrated Superdex 200 Increase 10/300 GL column (Cytiva). Proteins were eluted with SEC buffer (25 mM HEPES pH 7.4, 150 mM NaCl, 1 mM DTT). Fractions were analyzed by SDS-PAGE and Coomassie staining. Fractions containing purified GFP-FIP200(CTR) were pooled. After concentrating the purified protein, the protein was aliquoted and snap-frozen in liquid nitrogen. Proteins were stored at -80°C. A detailed protocol is available (https://doi.org/10.17504/protocols.io.j8nlk8866l5r/v1).

To purify the MBP-ULK1 from HEK293F cells, we expressed the ULK1 kinase from a pCAG backbone encoding MBP-TSF-TEV-ULK1 (RRID:Addgene_171416). The protein was expressed in FreeStyle^TM^ HEK293F cells, grown at 37°C in FreeStyle^TM^ 293 Expression Medium (Thermo, 12338-026). The day before transfection, cells were seeded at a density of 0.7 × 10^6 cells per ml. On the day of transfection, a 400 ml culture was transfected with 400 ug of the MAXI-prep DNA, diluted in 13 ml of Opti-MEMR I Reduced Serum Medium (Thermo, 31985-062), and 800 ug Polyethylenimine (PEI 25K, Polysciences CatNo 23966-1), also diluted in 13 ml of Opti-MEM media. One day post transfection, the culture was supplemented with 100 ml EXCELL R 293 Serum-Free Medium (SigmaA-ldrich, 14571C-1000ML). Another 24 h later, cells were harvested by centrifugation at 270 *g* for 20 min. The pellet was washed with PBS to remove medium and then flash-frozen in liquid nitrogen. Pellets were stored at - 80°C. For purification of MBP-TSF-TEV-ULK1, the cell pellet was resuspended in 25 ml lysis buffer (50 mM Tris-HCl pH 7.4, 200 mM NaCl, 1 mM MgCl_2_, 10% glycerol, 0.5% CHAPS, 1 mM TCEP, 1 µl benzonase (Sigma), cOmplete EDTA-free protease inhibitors (Roche), CIP protease inhibitor (Sigma)). Cells were homogenized with a douncer. Cell lysates were cleared by centrifugation at 10,000*g* for 45 min at 4°C with a SORVAL RC6+ centrifuge with an F21S-8x50Y rotor (Thermo Scientific). The soluble supernatant was collected and loaded on a StrepTrap 5ml HP column for binding of the Twin-Strep-tagged ULK1 protein, washed with 6 column volumes of wash buffer (50 mM Tris-HCl pH 7.4, 200 mM NaCl, 1 mM DTT), and eluted with elution buffer (50 mM Tris-HCl pH 7.4, 200 mM NaCl, 1 mM DTT, and 2.5 mM Desthiobiotin). Fractions were analyzed by SDS-PAGE and Coomassie staining. Fractions containing MBP-ULK1 were pooled and concentrated 50 kDa cut-off Amicon filter (Merck Millipore). The proteins were loaded onto a pre-equilibrated Superose 6 Increase 10/300 GL column (Cytiva). Proteins were eluted with SEC buffer (25 mM HEPES pH 7.5, 150 mM NaCl, 1 mM DTT). Fractions were analyzed by SDS-PAGE and Coomassie staining. Fractions containing purified MBP-ULK1 were pooled. After concentrating the purified protein, the protein was aliquoted and snap-frozen in liquid nitrogen. Proteins were stored at -80°C. A detailed protocol can be found here (https://doi.org/10.17504/protocols.io.bvn2n5ge).

To purify TBK1, we purchased gene-synthesized codon-optimized GST-TEV-TBK1 in a pFastBac-Dual vector from Genscript (RRID:Addgene_208875, RRID:Addgene_187830, RRID:Addgene_198033) for expression in insect cells. The V1 virus was generated as described above for BNIP3-GST. For expressions, we infected 1 L of Sf9 cells (12659017, Thermo Fisher, RRID:CVCL_0549), at 1 million cells per ml, with 1 ml of V1 virus. When the viability of the cells decreased to 90-95%, cells were collected by centrifugation. Cell pellets were washed with 1x PBS and flash-frozen in liquid nitrogen. Pellets were stored at -80°C. For purification of TBK1, pellets were resuspended in 25 ml lysis buffer (50 mM Tris-HCl pH 7.4, 300 mM NaCl, 1 mM DTT, 2 mM MgCl_2_, 5% glycerol, 2 mM β-mercaptoethanol, 1 µl benzonase (Sigma), cOmplete EDTA-free protease inhibitors (Roche), CIP protease inhibitor (Sigma)). Cells were homogenized with a douncer and lysates were cleared by centrifugation at 18,000 rpm for 45 min at 4°C in a SORVAL RC6+ centrifuge with an F21S-8x50Y rotor (Thermo Scientific). The supernatant was collected and incubated with pre-equilibrated Glutathione Sepharose 4B beads (GE Healthcare) for 2 h at 4°C with gentle shaking to bind GST-TEV-TBK1. Samples were centrifuged to pellet the beads and remove the unbound lysate. Beads were then washed five times with wash buffer (50 mM Tris-HCl pH 7.4, 300 mM NaCl, 5% glycerol, 1 mM DTT). Beads were incubated overnight with TEV protease in wash buffer (50 mM Tris-HCl pH 7.4, 300 mM NaCl, 5% glycerol, 1 mM DTT) at 4°C. After the proteins were released from the beads by the TEV protease, the supernatant was collected after centrifugation of the beads. The beads were washed twice with 4 ml of wash buffer, and the supernatant was collected. The supernatant fractions were pooled, filtered through a 0.45 µm syringe filter, and concentrated with a 30 kDa cut-off Amicon filter (Merck Millipore). The proteins were loaded onto a pre-equilibrated Superdex 200 Increase 10/300 GL column (Cytiva). Proteins were eluted with SEC buffer (25 mM Tris-HCl pH 7.4, 300 mM NaCl, 1 mM DTT). Fractions were analyzed by SDS-PAGE and Coomassie staining. Fractions containing purified TBK1 were pooled. After concentrating the purified protein, the protein was aliquoted and snap-frozen in liquid nitrogen. Proteins were stored at -80°C. A detailed protocol can be found here (https://doi.org/10.17504/protocols.io.81wgb6wy1lpk/v1).

To purify Src (WT and Y530F), we purchased gene-synthesized codon-optimized GST-TEV-Src in a pFastBac-Dual vector from Genscript (RRID:Addgene_223742; Addgene_223743) for expression in insect cells. The V1 virus was generated as described above for BNIP3-GST. For expressions, we infected 1 L of Sf9 cells (12659017, Thermo Fisher, RRID:CVCL_0549), at 1 million cells per ml, with 1 ml of V1 virus. When the viability of the cells decreased to 90-95%, cells were collected by centrifugation. Cell pellets were washed with 1x PBS and flash-frozen in liquid nitrogen. Pellets were stored at -80°C. For purification of Src(Y530F), pellets were resuspended in 25 ml lysis buffer (50 mM Tris-HCl pH 7.4, 300 mM NaCl, 1 mM DTT, 2 mM MgCl_2_, 2 mM β-mercaptoethanol, 5% glycerol, 1% Triton X-100, 1 µl benzonase (Sigma), cOmplete EDTA-free protease inhibitors (Roche), CIP protease inhibitor (Sigma)). Cells were homogenized with a douncer and lysates were cleared by centrifugation at 18,000 rpm for 45 min at 4°C in a SORVAL RC6+ centrifuge with an F21S-8x50Y rotor (Thermo Scientific). The supernatant was collected and incubated with pre-equilibrated Glutathione Sepharose 4B beads (GE Healthcare) for 2 h at 4°C with gentle shaking to bind GST-TEV-Src(Y530F). Samples were centrifuged to pellet the beads and remove the unbound lysate. Beads were then washed twice with wash buffer (50 mM Tris-HCl pH 7.4, 300 mM NaCl, 5% glycerol, 1 mM DTT), once with high salt wash buffer (50 mM Tris-HCl pH 7.4, 700 mM NaCl, 5% glycerol, 1 mM DTT), and two more times with wash buffer (50 mM Tris-HCl pH 7.4, 300 mM NaCl, 5% glycerol, 1 mM DTT). Beads were incubated overnight with TEV protease in wash buffer (50 mM Tris-HCl pH 7.4, 300 mM NaCl, 5% glycerol, 1 mM DTT) at 4°C. After the proteins were released from the beads by the TEV protease, the supernatant was collected after centrifugation of the beads. The beads were washed twice with 4 ml of wash buffer, and the supernatant was collected. The supernatant fractions were pooled, filtered through a 0.45 µm syringe filter, and concentrated with a 30 kDa cut-off Amicon filter (Merck Millipore). The proteins were loaded onto a pre-equilibrated Superdex 200 Increase 10/300 GL column (Cytiva). Proteins were eluted with SEC buffer (25 mM Tris-HCl pH 7.4, 300 mM NaCl, 1 mM DTT). Fractions were analyzed by SDS-PAGE and Coomassie staining. Fractions containing purified Src(WT or Y530F) were pooled. After concentrating the purified protein, the protein was aliquoted and snap-frozen in liquid nitrogen. Proteins were stored at -80°C. A detailed protocol can be found here (https://doi.org/10.17504/protocols.io.bp2l622mrgqe/v1).

To purify the CK2 kinase complex, we subcloned GST-TEV-CK2α together with CK2β in a pFastBac-Dual vector (RRID:Addgene_223740) and GST-TEV-CK2α’ together with CK2β in a pFastBac-Dual vector (RRID:Addgene_223741) for co-expression in insect cells. The V1 virus was generated as described above for BNIP3-GST. For expressions, we infected 1 L of Sf9 cells (12659017, Thermo Fisher, RRID:CVCL_0549), at 1 million cells per ml, with 1 ml of V1 virus for GST-TEV-CK2α/CK2β and 1 ml of V1 virus for GST-TEV-CK2α’/CK2β. When the viability of the co-infected cells decreased to 90-95%, cells were collected by centrifugation. Cell pellets were washed with 1x PBS and flash-frozen in liquid nitrogen. Pellets were stored at -80°C. For purification of the CK2 kinase complex, pellets were resuspended in 25 ml lysis buffer (50 mM Tris-HCl pH 7.4, 300 mM NaCl, 1 mM DTT, 2 mM MgCl_2_, 2 mM β-mercaptoethanol, 5% glycerol, 1% Triton X-100, 1 µl benzonase (Sigma), cOmplete EDTA-free protease inhibitors (Roche), CIP protease inhibitor (Sigma)). Cells were homogenized with a douncer and lysates were cleared by centrifugation at 18,000 rpm for 45 min at 4°C in a SORVAL RC6+ centrifuge with an F21S-8x50Y rotor (Thermo Scientific). The supernatant was collected and incubated with pre-equilibrated Glutathione Sepharose 4B beads (GE Healthcare) for 2 h at 4°C with gentle shaking to bind the CK2 complex. Samples were centrifuged to pellet the beads and remove the unbound lysate. Beads were then washed twice with wash buffer (50 mM Tris-HCl pH 7.4, 300 mM NaCl, 5% glycerol, 1 mM DTT), once with high salt wash buffer (50 mM Tris-HCl pH 7.4, 700 mM NaCl, 5% glycerol, 1 mM DTT), and two more times with wash buffer (50 mM Tris-HCl pH 7.4, 300 mM NaCl, 5% glycerol, 1 mM DTT). Beads were incubated overnight with TEV protease in wash buffer (50 mM Tris-HCl pH 7.4, 300 mM NaCl, 5% glycerol, 1 mM DTT) at 4°C. After the proteins were released from the beads by the TEV protease, the supernatant was collected after centrifugation of the beads. The beads were washed twice with 4 ml of wash buffer, and the supernatant was collected. The supernatant fractions were pooled, filtered through a 0.45 µm syringe filter, and concentrated with a 10 kDa cut-off Amicon filter (Merck Millipore). The proteins were loaded onto a pre-equilibrated Superdex 200 Increase 10/300 GL column (Cytiva). Proteins were eluted with SEC buffer (25 mM Tris-HCl pH 7.4, 300 mM NaCl, 1 mM DTT). Fractions were analyzed by SDS-PAGE and Coomassie staining. Fractions containing purified CK2α/CK2α’/CK2β were pooled. After concentrating the purified protein, the protein was aliquoted and snap-frozen in liquid nitrogen. Proteins were stored at -80°C. A detailed protocol can be found here (https://doi.org/10.17504/protocols.io.eq2lyww1evx9/v1).

To purify Lambda protein phosphatase (λ PPase), the protein phosphatase was fused to a N-terminal 6xHis-tag through cloning into a pET-DUET1 vector (RRID:Addgene_223747). After the transformation of the pET-DUET1 vector encoding 6xHis-TEV-λ PPase in *E. coli* Rosetta pLysS cells (Novagen Cat# 70956-4), cells were grown in 2x Tryptone Yeast extract (TY) medium at 37°C until an OD_600_ of 0.4 and then continued at 18°C. Once the cells reached an OD_600_ of 0.8, protein expression was induced with 100 µM isopropyl β-D-1-thiogalactopyranoside (IPTG) for 16 h at 18°C. Cells were collected by centrifugation and resuspended in lysis buffer (50 mM Tris-HCl pH 7.4, 300 mM NaCl, 2 mM MgCl_2_, 5% glycerol, 10 mM Imidazole, 2 mM β-mercaptoethanol, cOmplete EDTA-free protease inhibitors (Roche), CIP protease inhibitor (Sigma), and DNase (Sigma)). Cell lysates were sonicated twice for 30 s. Lysates were cleared by centrifugation at 18,000 rpm for 45 min at 4°C in a SORVAL RC6+ centrifuge with an F21S-8x50Y rotor (Thermo Scientific). The supernatant was filtered through an 0.45 µm filter and loaded onto a pre-equilibrated 5 ml His-Trap HP column (Cytiva). After His-tagged proteins were bound to the column, the column was washed with three column volumes of wash buffer (50 mM Tris-HCl pH 7.4, 300 mM NaCl, 10 mM Imidazole, 2 mM β-mercaptoethanol). Proteins were then eluted with a stepwise imidazole gradient (30, 75, 100, 150, 225, 300 mM). Fractions containing the 6xHis-TEV-λ PPase were pooled and incubated overnight with TEV protease at 4°C. After the 6xHis tag was cleaved off, 6xHis tag and His-tagged TEV protease was recaptured with nickel beads for 1 h at 4 degrees. The beads were pelleted by centrifugation and the supernatant, containing the λ PPase protein was concentrated using a 30 kDa cut-off Amicon filter (Merck Millipore) and loaded onto a pre-equilibrated Superdex 200 Increase 10/300 GL column (Cytiva). Proteins were eluted with SEC buffer (25 mM Tris-HCl pH 7.4, 150 mM NaCl, 1 mM DTT). Fractions were analyzed by SDS-PAGE and Coomassie staining. Fractions containing purified λ PPase were pooled. After concentrating the purified protein, the protein was aliquoted and snap-frozen in liquid nitrogen. Proteins were stored at -80°C. A detailed protocol is available (https://doi.org/10.17504/protocols.io.kqdg322bqv25/v1).

To purify mCherry-WIPI2d and mCherry-WIPI3, as described previously for WIPI2d^74^, the coding sequence of WIPI2d or WIPI3 was fused to a N-terminal 6xHis-TEV-mCherry-tag through cloning into a pET-DUET1 vector (RRID:Addgene_223725; RRID:Addgene_223763). After the transformation of the pET-DUET1 vector encoding 6xHis-TEV-mCherry-WIPI2d/WIPI3 in *E. coli* Rosetta pLysS cells (Novagen Cat# 70956-4), cells were grown in 2x Tryptone Yeast extract (TY) medium at 37°C until an OD_600_ of 0.4 and then continued at 18°C. Once the cells reached an OD_600_ of 0.8, protein expression was induced with 100 µM isopropyl β-D-1-thiogalactopyranoside (IPTG) for 16 h at 18°C. Cells were collected by centrifugation and resuspended in lysis buffer (50 mM Tris-HCl pH 7.4, 300 mM NaCl, 2 mM MgCl_2_, 5% glycerol, 1% Triton X-100, 10 mM Imidazole, 2 mM β-mercaptoethanol, cOmplete EDTA-free protease inhibitors (Roche), CIP protease inhibitor (Sigma), and DNase (Sigma)). Cell lysates were sonicated twice for 30 s. Lysates were cleared by centrifugation at 18,000 rpm for 45 min at 4°C in a SORVAL RC6+ centrifuge with an F21S-8x50Y rotor (Thermo Scientific). The supernatant was filtered through an 0.45 µm filter and loaded onto a pre-equilibrated 5 ml His-Trap HP column (Cytiva). After His tagged proteins were bound to the column, the column was washed with three column volumes of wash buffer (50 mM Tris-HCl pH 7.4, 300 mM NaCl, 10 mM Imidazole, 2 mM β-mercaptoethanol). Proteins were then eluted with a stepwise imidazole gradient (30, 75, 100, 150, 225, 300 mM). Fractions containing the 6xHis-TEV-mCherry-WIPI2d/WIPI3 were pooled, concentrated using a 30 kDa cut-off Amicon filter (Merck Millipore) and loaded onto a pre-equilibrated Superdex 200 Increase 10/300 GL column (Cytiva). Proteins were eluted with SEC buffer (25 mM Tris-HCl pH 7.4, 150 mM NaCl, 1 mM DTT). Fractions were analyzed by SDS-PAGE and Coomassie staining. Fractions containing purified mCherry-WIPI2d or mCherry-WIPI3 were pooled. After concentrating the purified protein, the protein was aliquoted and snap-frozen in liquid nitrogen. Proteins were stored at -80°C. A detailed protocol is available (https://doi.org/10.17504/protocols.io.4r3l2qqyql1y/v1).

To purify mCherry-WIPI2d K87A/K88A (RRID:Addgene_223751) or mCherry-WIPI2d IDR (364-425aa) (RRID:Addgene_223790), the same expression and purification methods were used as described above for full-length mCherry-WIPI2d with the exception that for the mCherry-WIPI2d IDR we used the S75 Increase 10/300 column. A adapted protocol is available (https://doi.org/10.17504/protocols.io.5qpvokk8bl4o/v1) To purify GST-WIPI1/GST-WIPI2/GST-WIPI3/GST-WIPI4, we expressed the GST-tagged WIPI1/2d/3/4 from a pCAG backbone encoding GST-TEV-WIPI1/2/3/4 (RRID:Addgene_223798; RRID:Addgene_223799; RRID:Addgene_223800; RRID:Addgene_223801). The protein was expressed in FreeStyle^TM^ HEK293F cells, grown at 37°C in FreeStyle^TM^ 293 Expression Medium (Thermo, 12338-026). The day before transfection, cells were seeded at a density of 0.7 × 10^6 cells per ml. On the day of transfection, a 400 ml culture was transfected with 400 ug of the MAXI-prep DNA, diluted in 13 ml of Opti-MEMR I Reduced Serum Medium (Thermo, 31985-062), and 800 ug Polyethylenimine (PEI 25K, Polysciences CatNo 23966-1), also diluted in 13 ml of Opti-MEM media. One day post transfection, the culture was supplemented with 100 ml EXCELL R 293 Serum-Free Medium (Sigma-Aldrich, 14571C-1000ML). Another 24 h later, cells were harvested by centrifugation at 270 *g* for 20 min. The pellet was washed with PBS to remove medium and then flash-frozen in liquid nitrogen. Pellets were stored at -80°C. For purification of GST-TEV-WIPI1/2/3/4, the cell pellet was resuspended in 25 ml lysis buffer (50 mM Tris-HCl pH 7.4, 300 mM NaCl, 2 mM MgCl_2_, 5% glycerol, 1% Triton X-100, 2 mM β-mercaptoethanol, cOmplete EDTA-free protease inhibitors (Roche), CIP protease inhibitor (Sigma), and DNase (Sigma)). Cell lysates were sonicated twice for 30 s. Cell lysates were cleared by centrifugation at 10,000*g* for 45 min at 4°C with a SORVAL RC6+ centrifuge with an F21S-8x50Y rotor (Thermo Scientific). The supernatant was collected and incubated with pre-equilibrated Glutathione Sepharose 4B beads (GE Healthcare) for 2 h at 4°C with gentle shaking to bind GST-TEV-WIPI1/2/3/4. Samples were centrifuged to pellet the beads and remove the unbound lysate. Beads were then washed twice with wash buffer (50 mM Tris-HCl pH 7.4, 300 mM NaCl, 1 mM DTT), once with high salt wash buffer (50 mM Tris-HCl pH 7.4, 700 mM NaCl, 1 mM DTT), and two more times with wash buffer (50 mM Tris-HCl pH 7.4, 300 mM NaCl, 1 mM DTT). Beads were incubated overnight with 4 ml of 50 mM reduced glutathione dissolved in wash buffer (50 mM Tris-HCl pH 7.4, 300 mM NaCl, 1 mM DTT) at 4°C, to elute GST-tagged WIPI1/2/3/4 from the beads. To collect the supernatant, the beads were collected by centrifugation. The beads were washed twice with 4 ml of wash buffer, and the supernatant was collected. The supernatant fractions were pooled, filtered through a 0.45 µm syringe filter, concentrated with 30 kDa cut-off Amicon filter (Merck Millipore), and loaded onto a pre-equilibrated Superdex 200 Increase 10/300 GL column (Cytiva). Proteins were eluted with SEC buffer (25 mM Tris-HCl pH 7.4, 150 mM NaCl, 1 mM DTT). Fractions were analyzed by SDS-PAGE and Coomassie staining. Fractions containing purified GST-TEV-WIPI1/2/3/4 were pooled. After concentrating the purified protein, the protein was aliquoted and snap-frozen in liquid nitrogen. Proteins were stored at -80°C. A detailed protocol is available (https://doi.org/10.17504/protocols.io.n2bvjnnqxgk5/v1).

To purify the mCherry-tagged or GFP-tagged ATG13/101 subcomplex, we expressed mCherry-tagged ATG13 from a pCAG backbone (RRID:Addgene_223735) together with GST-TEV-ATG101 (RRID:Addgene_171414) or GST-TEV-GFP-tagged ATG13 (RRID:Addgene_223797) together with ATG101 (RRID:Addgene_223796). The subcomplex was expressed in FreeStyle^TM^ HEK293F cells, grown at 37°C in FreeStyle^TM^ 293 Expression Medium (Thermo, 12338-026). The day before transfection, cells were seeded at a density of 0.7 × 10^6 cells per ml. On the day of transfection, a 400 ml culture was transfected with 400 µg of plasmid at a molar 1:1 ratio, diluted in 13 ml of Opti-MEMR I Reduced Serum Medium (Thermo, 31985-062), and 800 ug Polyethylenimine (PEI 25K, Polysciences CatNo 23966-1), also diluted in 13 ml of Opti-MEM media. One day post transfection, the culture was supplemented with 100 ml EXCELL R 293 Serum-Free Medium (Sigma-Aldrich, 14571C-1000ML). Another 24 h later, cells were harvested by centrifugation at 270 *g* for 20 min. The pellet was washed with PBS to remove medium and then flash-frozen in liquid nitrogen. Pellets were stored at -80°C. For purification of the ATG13/101 subcomplex, the cell pellet was resuspended in 25 ml lysis buffer (50 mM Tris-HCl pH 7.4, 200 mM NaCl, 2 mM MgCl_2_, 10% glycerol, 1% Triton X-100, 2 mM β-mercaptoethanol, cOmplete EDTA-free protease inhibitors (Roche), CIP protease inhibitor (Sigma), and Benzonase). Cells were homogenized with a douncer and lysates were cleared by centrifugation at 10,000*g* for 45 min at 4°C with a SORVAL RC6+ centrifuge with an F21S-8x50Y rotor (Thermo Scientific). The supernatant was collected and incubated with pre-equilibrated Glutathione Sepharose 4B beads (GE Healthcare) for 2 h at 4°C with gentle shaking to bind GST-TEV-ATG101/mCherry-ATG13 or GST-TEV-GFP-ATG13/ATG101. Samples were centrifuged to pellet the beads and remove the unbound lysate. Beads were then washed twice with wash buffer I (50 mM Tris-HCl pH 7.4, 200 mM NaCl, 2 mM MgCl_2_, 1 mM DTT, 1% Triton X-100, 10% glycerol) followed by three washes in wash buffer II (50 mM Tris-HCl pH 7.4, 200 mM NaCl, 2 mM MgCl_2_, 1 mM DTT). Beads were incubated overnight with TEV protease in wash buffer (50 mM Tris-HCl pH 7.4, 200 mM NaCl, 2 mM MgCl_2_, 1 mM DTT) at 4°C, to release mCherry- or GFP-tagged ATG13/101 from the beads. To collect the supernatant, the beads were collected by centrifugation. The beads were washed twice with 4 ml of wash buffer, and the supernatant was collected. The supernatant fractions were pooled, filtered through a 0.45 µm syringe filter, concentrated with 10 kDa cut-off Amicon filter (Merck Millipore), and loaded onto a pre-equilibrated Superose S6 Increase 10/300 GL column (Cytiva). Proteins were eluted with SEC buffer (50 mM Tris-HCl pH 7.4, 200 mM NaCl, 1 mM MgCl_2_, 1 mM DTT). Fractions were analyzed by SDS-PAGE and Coomassie staining. Fractions containing both ATG13/101 were pooled. After concentrating the purified protein, the protein was aliquoted and snap-frozen in liquid nitrogen. Proteins were stored at -80°C. A detailed protocol is available (https://doi.org/10.17504/protocols.io.yxmvmepdng3p/v1).

To purify mCherry-ATG13/101 HORMA dimer, we expressed mCherry-tagged ATG13 (1-191aa) from a pCAG backbone (RRID:Addgene_223759) together with GST-TEV-ATG101 (RRID:Addgene_171414). The same expression and purification methods were used as described above for full-length mCherry-ATG13/101. A detailed protocol is available (https://doi.org/10.17504/protocols.io.n92ld8wo9v5b/v1).

To purify GFP-tagged or mCherry-tagged ATG13 IDR, the coding sequence for ATG13 (191-517aa) or ATG13 (230-517aa) were fused to a N-terminal 6xHis-TEV-mCherry-tag through cloning into a pET-DUET1 vector (RRID:Addgene_223762) or by inserting the coding sequence for ATG13 (191-517aa), (205-517aa), (231-517aa), (191-205_231-517aa), (191-230aa), (191-205aa), or (206-230aa) into GST-TEV-EGFP-insert through cloning into a pGEX-4T1 vector (RRID:Addgene_223760; RRID:Addgene_223786; RRID:Addgene_223785; RRID:Addgene_223787; RRID:Addgene_223792; RRID:Addgene_223791; RRID:Addgene_223793). Mutants 3A (M196A/S197A/R199A; RRID:Addgene_223761) and 11A (M196A/S197A/R199A/G202A/T204A/P205A/I207A/M208A/I210A/D213A/H214A; RRID:Addgene_223779) were also expressed according to the protocol below. After the transformation of the pET-DUET1 or pGEX-4T1 vectors encoding the GFP-tagged or mCherry-tagged ATG13 IDR in *E. coli* Rosetta pLysS cells (Novagen Cat# 70956-4), cells were grown in 2x Tryptone Yeast extract (TY) medium at 37°C until an OD_600_ of 0.4 and then continued at 18°C. Once the cells reached an OD_600_ of 0.8, protein expression was induced with 100 µM isopropyl β-D-1-thiogalactopyranoside (IPTG) for 16 h at 18°C. Cells were collected by centrifugation and resuspended in lysis buffer (a) for His-tagged proteins (50 mM Tris-HCl pH 7.4, 300 mM NaCl, 2 mM MgCl_2_, 5% glycerol, 1% Triton X-100, 10 mM Imidazole, 2 mM β-mercaptoethanol, cOmplete EDTA-free protease inhibitors (Roche), CIP protease inhibitor (Sigma), and DNase (Sigma)), or (b) for GST-tagged proteins (50 mM Tris-HCl pH 7.4, 300 mM NaCl, 2 mM MgCl_2_, 5% glycerol, 1% Triton X-100, 2 mM β-mercaptoethanol, cOmplete EDTA-free protease inhibitors (Roche), CIP protease inhibitor (Sigma), and DNase (Sigma)),). Cell lysates were sonicated twice for 30 s. Lysates were cleared by centrifugation at 18,000 rpm for 45 min at 4°C in a SORVAL RC6+ centrifuge with an F21S-8x50Y rotor (Thermo Scientific). The supernatant was filtered through an 0.45 µm filter and loaded onto a pre-equilibrated 5 ml His-Trap HP column (Cytiva), in case of 6xHis-mCherry-tagged ATG13. After His tagged proteins were bound to the column, the column was washed with three column volumes of wash buffer (50 mM Tris-HCl pH 7.4, 300 mM NaCl, 10 mM Imidazole, 2 mM β-mercaptoethanol). Proteins were then eluted with a stepwise imidazole gradient (30, 75, 100, 150, 225, 300 mM). Fractions containing the 6xHis-TEV-mCherry-ATG13 IDR were pooled, concentrated using a 30 kDa cut-off Amicon filter (Merck Millipore). In case of GST-TEV-EGFP-tagged ATG13 IDR, the supernatant was collected after centrifugation and incubated with pre-equilibrated Glutathione Sepharose 4B beads (GE Healthcare) for 2 h at 4°C with gentle shaking to bind GST-TEV-EGFP-ATG13 IDR. Samples were centrifuged to pellet the beads and remove the unbound lysate. Beads were then washed twice with wash buffer (50 mM Tris-HCl pH 7.4, 300 mM NaCl, 1 mM DTT), once with high salt wash buffer (50 mM Tris-HCl pH 7.4, 700 mM NaCl, 1 mM DTT), and two more times with wash buffer (50 mM Tris-HCl pH 7.4, 300 mM NaCl, 1 mM DTT). Beads were incubated overnight with TEV protease at 4°C, to elute GFP-tagged ATG13 IDR from the beads. To collect the supernatant, the beads were collected by centrifugation. The beads were washed twice with 4 ml of wash buffer, and the supernatant was collected. The supernatant fractions were pooled, filtered through a 0.45 µm syringe filter, concentrated with 10 or 30 kDa cut-off Amicon filter (Merck Millipore). Samples were loaded onto a pre-equilibrated Superose 200 Increase 10/300 GL column (Cytiva) or S75 Increase 10/300 column (Cytiva) in case of the smaller peptides (190-230aa and variants thereof). Proteins were eluted with SEC buffer (25 mM Tris-HCl pH 7.4, 150 mM NaCl, 1 mM DTT). Fractions were analyzed by SDS-PAGE and Coomassie staining. Fractions containing purified ATG13 IDR were pooled. After concentrating the purified protein, the protein was aliquoted and snap-frozen in liquid nitrogen. Proteins were stored at -80°C. Detailed protocols are available (https://doi.org/10.17504/protocols.io.8epv5rey4g1b/v1) and (https://doi.org/10.17504/protocols.io.81wgbz4m1gpk/v1).

To purify GFP-tagged ULK1-complex, as described previously^37^, we co-expressed GST-TEV-FIP200-MBP/EGFP-ATG13/ATG101 from a pCAG backbones (RRID:Addgene_171410; RRID:Addgene_171413; RRID:Addgene_189590) in parallel to MBP-Strep-Strep-Flag-TEV-ULK1 (RRID:Addgene_171416). The subcomplex FIP200/EGFP-ATG13/ATG101 was transfected and expressed separately from the ULK1 subunit in FreeStyle^TM^ HEK293F cells, grown at 37°C in FreeStyle^TM^ 293 Expression Medium (Thermo, 12338-026). The day before transfection, cells were seeded at a density of 0.7 × 10^6 cells per ml. On the day of transfection, a 400 ml culture was transfected with 400 µg of plasmid at a molar 1:1 ratio, diluted in 13 ml of Opti-MEMR I Reduced Serum Medium (Thermo, 31985-062), and 800 ug Polyethylenimine (PEI 25K, Polysciences CatNo 23966-1), also diluted in 13 ml of Opti-MEM media. One day post transfection, the culture was supplemented with 100 ml EXCELL R 293 Serum-Free Medium (Sigma-Aldrich, 14571C-1000ML). Another 24 h later, cells were harvested by centrifugation at 270 *g* for 20 min. The pellet was washed with PBS to remove medium and then flash-frozen in liquid nitrogen. Pellets were stored at -80°C. For purification of the FIP200/ATG13/101 subcomplex, the cell pellet was resuspended in 25 ml lysis buffer (50 mM Tris-HCl pH 7.4, 200 mM NaCl, 2 mM MgCl_2_, 1 mM DTT, 10% glycerol, 1% Triton X-100, cOmplete EDTA-free protease inhibitors (Roche), CIP protease inhibitor (Sigma), and Benzonase). Cells were homogenized with a douncer and lysates were cleared by centrifugation at 10,000*g* for 45 min at 4°C with a SORVAL RC6+ centrifuge with an F21S-8x50Y rotor (Thermo Scientific). The supernatant was collected and incubated with pre-equilibrated Glutathione Sepharose 4B beads (GE Healthcare) in case of GST-TEV-FIP200-MBP/EGFP-ATG13/ATG101 overnight at 4°C with gentle shaking to bind GST-TEV-FIP200-MBP/EGFP-13/ATG101, or incubated with Strep-Tactin Sepharose beads overnight at 4°C in case of MBP-TEV-ULK1. Samples were centrifuged to pellet the beads and remove the unbound lysate. Beads were then washed three times with wash buffer I (50 mM Tris-HCl pH 7.4, 500 mM NaCl, 1 mM MgCl_2_, 1 mM DTT, 1% Triton X-100, 10% glycerol) followed by three washes in wash buffer II (50 mM Tris-HCl pH 7.4, 500 mM NaCl, 1 mM MgCl_2_, 1 mM DTT). Beads were incubated for 1 h with 50 mM gluthathione in wash buffer (50 mM Tris-HCl pH 7.4, 200 mM NaCl, 2 mM MgCl_2_, 1 mM DTT) at 4°C in case of FIP200/ATG13/ATG101 subcomplex, to release GFP-tagged FIP200/ATG13/ATG101 from the beads, or 10 mM desthiobiotin to elute ULK1 from the Strep-Tactin beads. The eluates were then mixed in presence of TEV protease and placed on a roller for 1 h at 4°C before being transferred to the fridge to allow complex formation overnight. The next morning, the complex is collected by affinity purification using FIP200-MBP and incubating the complex with Amylose resin (BioLabs) for 1 h at 4°C. The resin was then washed with wash buffer II and finally eluted with 2x 1 ml wash buffer containing 50 mM Maltose (D-maltose monohydrate, ChemCruz). The elutions were pooled, filtered through a 0.45 µm syringe filter, concentrated with 30 kDa cut-off Amicon filter (Merck Millipore), and loaded onto a pre-equilibrated Superose S6 Increase 10/300 GL column (Cytiva). Proteins were eluted with SEC buffer (50 mM Tris-HCl pH 7.4, 500 mM NaCl, 1 mM DTT). Fractions were analyzed by SDS-PAGE and Coomassie staining. Fractions containing the ULK1 complex were pooled. After concentrating the purified protein, the protein was aliquoted and snap-frozen in liquid nitrogen. Proteins were stored at -80°C. A detailed protocol is available (https://doi.org/10.17504/protocols.io.bvn2n5ge).

To purify mCherry-tagged PI3KC3-C1 complex, as published before^74^, the codon-optimized genes were purchased from Genscript and cloned by the Vienna BioCenter Core Facilities (VBCF) Protech Facility as GST-3C-mCherry-ATG14/VPS34/VPS15/BECN1 in a pGBdest vector (RRID:Addgene_187936). The construct was used to generate bacmid DNA, using the Bac-to-Bac system, by amplification in DH10BacY cells ^73^. After the bacmid DNA was verified by PCR for insertion of the transgene, we purified bacmid DNA for transfection into Sf9 insect cells (12659017, Thermo Fisher, RRID:CVCL_0549). To this end, we mixed 2500 ng of plasmid DNA with FuGene transfection reagent (Promega) and transfected 1 million Sf9 cells seeded in a 6 well plate. About 7 days after transfection, the V0 virus was harvested and used to infect 40 ml of 1 million cells per ml of Sf9 cells. The viability of the cultures was closely monitored and upon the decrease in viability and confirmation of yellow fluorescence, we collected the supernatant after centrifugation and stored this as V1 virus. For expressions, we infected 1 L of Sf9 cells, at 1 million cells per ml, with 1 ml of V1 virus. When the viability of the cells decreased to 90-95%, cells were collected by centrifugation. Cell pellets were washed with 1x PBS and flash-frozen in liquid nitrogen. Pellets were stored at -80°C. For purification of GST-3C-mCherry-ATG14/VPS34/VPS15/BECN1, pellets were resuspended in 25 ml lysis buffer (50 mM HEPES pH 7.4, 300 mM NaCl, 0.5% CHAPS, 1 mM DTT, 1 mM MgCl_2_, 1 µl benzonase (Sigma), cOmplete EDTA-free protease inhibitors (Roche), CIP protease inhibitor (Sigma)). Cells were homogenized with a douncer and cell lysates were cleared by centrifugation at 18,000 rpm for 45 min at 4°C in a SORVAL RC6+ centrifuge with an F21S-8x50Y rotor (Thermo Scientific). The supernatant was collected and incubated with pre-equilibrated Glutathione Sepharose 4B beads (GE Healthcare) for 2 h at 4°C with gentle shaking to bind the GST-tagged PI3KC3-CI. Samples were centrifuged to pellet the beads and remove the unbound lysate. Beads were then washed twice with wash buffer I (50 mM HEPES pH 7.4, 300 mM NaCl, 0.5% CHAPS, 1 mM DTT), twice in wash buffer II (50 mM HEPES pH 7.4, 500 mM NaCl, 1 mM DTT), and two more times with wash buffer III (50 mM HEPES pH 7.4, 300 mM NaCl, 1 mM DTT). Beads were incubated overnight with C3 protease, to elute PI3KC3-C1 from the beads. To collect the supernatant, the beads were collected by centrifugation. The beads were washed twice with 4 ml of wash buffer, and the supernatant was collected. The supernatant fractions were pooled, filtered through a 0.45 µm syringe filter, concentrated with 30 kDa cut-off Amicon filter (Merck Millipore), and loaded onto a pre-equilibrated Superose 6 Increase 10/300 GL column (Cytiva). Proteins were eluted with SEC buffer (25 mM HEPES pH 7.4, 200 mM NaCl, 1 mM DTT). Fractions were analyzed by SDS-PAGE and Coomassie staining. Fractions containing purified mCherry-tagged PI3KC3-C1 complex were pooled. After concentrating the purified protein, the protein was aliquoted and snap-frozen in liquid nitrogen. Proteins were stored at -80°C. A detailed protocol is available (https://doi.org/10.17504/protocols.io.8epv59mz4g1b/v1).

To purify GST-LC3A, GST-LC3B, GST-LC3C, GST-GBRP, GST-GBRPL1, GST-GBRPL2, as previously described ^75^, we inserted human LC3/GBRP cDNA in a pGEX-4T1 vector (RRID:Addgene_223726; RRID:Addgene_216836; RRID:Addgene_223727; RRID:Addgene_223728; RRID:Addgene_223729; RRID:Addgene_223730). The last five amino acids of LC3/GBRP were deleted, to mimic the cleavage by ATG4. After the transformation of the pGEX-4T1 vector encoding GST-LC3/GBRP in *E. coli* Rosetta (DE3) pLysS cells, cells were grown in LB medium at 37°C until an OD_600_ of 0.8-1, protein expression was induced with 1 mM IPTG for 4 h at 37°C. Cells were collected by centrifugation and resuspended in lysis buffer (50 mM HEPES pH 7.5, 300 mM NaCl, 2 mM MgCl_2_, 2 mM β-mercaptoethanol, cOmplete EDTA-free protease inhibitors (Roche), and DNase (Sigma)). Cell lysates were sonicated twice for 30 s. Lysates were cleared by centrifugation at 140,000 x*g* for 30 min at 4°C in a Beckman Ti45 rotor. The supernatant was collected and incubated with pre-equilibrated Glutathione Sepharose 4B beads (GE Healthcare) for 2 h at 4°C with gentle shaking to bind GST-LC3/GBRP. Samples were centrifuged to pellet the beads and remove the unbound lysate. Beads were then washed twice with wash buffer (50 mM HEPES pH 7.4, 300 mM NaCl, 1 mM DTT), once with high salt wash buffer (50 mM HEPES pH 7.4, 700 mM NaCl, 1 mM DTT), and two more times with wash buffer (50 mM HEPES pH 7.4, 300 mM NaCl, 1 mM DTT). Proteins were eluted overnight with 20 mM reduced L-glutathione in 50 mM HEPES pH 7.4, 300 mM NaCl, 1 mM DTT buffer. The supernatant was collected, filtered through a 0.45 µm syringe filter, and concentrated using a 10 kDa cut-off Amicon filter (Merck Millipore), and loaded onto a pre-equilibrated Superdex 75 16/600 column (Cytiva). Proteins were eluted with SEC buffer (25 mM HEPES pH 7.4, 150 mM NaCl, 1 mM DTT). Fractions were analyzed by SDS-PAGE and Coomassie staining. Fractions containing purified GST-LC3/GBRP were pooled. After concentrating the purified protein, the protein was aliquoted and snap-frozen in liquid nitrogen. Proteins were stored at -80°C. A detailed protocol is available (https://doi.org/10.17504/protocols.io.3byl4qnbjvo5/v1).

To purify mCherry-tagged OPTN, we cloned human OPTN cDNA in a pETDuet-1 vector with an N-terminal 6xHis tag followed by a TEV cleavage site (RRID:Addgene_190191). After the transformation of the pETDuet-1 vector encoding 6xHis-TEV-mCherry-OPTN in E. coli Rosetta pLySS cells, cells were grown in 2xTY medium at 37°C until an OD600 of 0.4 and then continued at 18°C. Once the cells reached an OD600 of 0.8, protein expression was induced with 50 μM IPTG for 16 h at 18°C. Cells were collected by centrifugation and resuspended in lysis buffer (50 mM Tris-HCl pH 7.4, 300 mM NaCl, 2 mM MgCl2, 5% glycerol, 10 mM Imidazole, 2 mM β-mercaptoethanol, cOmplete EDTA-free protease inhibitors (Roche), CIP protease inhibitor (Sigma), and DNase (Sigma)). Cell lysates were sonicated twice for 30 s. Lysates were cleared by centrifugation at 18,000 rpm for 45 min at 4°C in a SORVAL RC6+ centrifuge with an F21S-8x50Y rotor (Thermo Scientific). The supernatant was filtered through an 0.45 μm filter and loaded onto a pre-equilibrated 5 ml His-Trap HP column (Cytiva). After His tagged proteins were bound to the column, the column was washed with three column volumes of wash buffer (50 mM Tris-HCl pH 7.4, 300 mM NaCl, 10 mM Imidazole, 2 mM β-mercaptoethanol). Proteins were then eluted with a stepwise imidazole gradient (30, 75, 100, 150, 225, 300 mM). Fractions at 75-100 mM imidazole contained the 6xHis-TEV-mCherry-OPTN and were pooled. The pooled samples were incubated overnight with TEV protease at 4°C. After the 6xHis tag was cleaved off, the protein was concentrated using a 50 kDa cut-off Amicon filter (Merck Millipore) and loaded onto a pre-equilibrated Superdex 200 Increase 10/300 GL column (Cytiva). Proteins were eluted with SEC buffer (25 mM Tris-HCl pH 7.4, 150 mM NaCl, 1 mM DTT). Fractions were analyzed by SDS-PAGE and Coomassie staining. Fractions containing purified mCherry-OPTN were pooled. After concentrating the purified protein, the protein was aliquoted and snap-frozen in liquid nitrogen. Proteins were stored at -80°C. A detailed protocol is available (https://doi.org/10.17504/protocols.io.4r3l225djl1y/v1). The negative controls EGFP, mCherry, and GST, were purified as previously described ^4,76^. Plasmids are available from Addgene (RRID:Addgene_223723).

### Microscopy-based bead assay

Glutathione Sepharose 4B beads (GE Healthcare) were used to bind GST-tagged bait proteins, GFP-trap agarose beads (ProteinTech) were used to bind GFP-tagged bait proteins, and RFP-trap agarose beads (ProteinTech) were used to bind mCherry-tagged bait proteins. To this end, 20 µl of beads were washed twice with dH_2_O and equilibrated with bead assay buffer (25 mM Tris-HCl pH 7.4, 150 mM NaCl, 1 mM DTT). Beads were then resuspended in 40 µl bead assay buffer, to which bait proteins were added at a final concentration of 5 µM. Beads were incubated with the bait proteins for 1 h at 4°C at a horizontal tube roller. Beads were then washed three times to remove unbound GST-/GFP-/mCherry-tagged bait proteins and resuspended in 30 µl bead assay buffer. Where indicated, we also added 10 mM MgCl_2_ and 100 µM ATP to the buffer to allow the phosphorylation of targets by kinases or added 1 mM MnCl_2_ to samples containing Lambda Protein Phosphatase. Glass-bottom 384-well microplates (Greiner Bio-One) were prepared with 20 µl samples containing prey proteins at the concentrations described below and diluted in bead assay buffer, and 3 µl of beads were added per well. The beads were incubated with the prey proteins for 30 min prior to imaging, with the exception of experiments containing full-length FIP200, where proteins were co-incubated for 4 h, and experiments where WIPI proteins and cargo receptors were tested for interactions, where proteins were co-incubated for 2 h, before imaging. Samples were imaged with a Zeiss LSM 700 confocal microscope equipped with Plan Apochromat 20X/0.8 WD 0.55 mm objective. Three biological replicates were performed for each experimental condition. A detailed protocol is available (https://doi.org/10.17504/protocols.io.14egn38pzl5d/v1).

### In vitro kinase assays

To verify the activity of the kinases TBK1 and MBP-ULK1, we mixed the kinases with mCherry-tagged OPTN or PI3K-complex (composed of VPS15, VPS34, ATG14, and Beclin1) were mixed in kinase buffer (20 mM Tris-HCl pH 7.4, 150 mM NaCl, 1 mM DTT). The kinases were used at 50 nM and mixed with 200 nM OPTN and 130 nM PI3K complex. The kinase reactions were started by the addition of 2x ATP/MgCl_2_ kinase buffer to a final concentration of 10 mM MgCl_2_ and 100 µM ATP. Protein mixtures were prepared as master mixes and divided over the number of time points. To control for potential protein instability, we induced the latest time point first and then went gradually to the shortest time point. In this way, all protein mixtures were kept at room temperature for the same time, and reactions could be terminated together. Termination of reactions was achieved by the addition of 6x Protein Loading dye and heat inactivation at 95°C for 5 min. Samples were separated on 4-12% SDS-PAGE gels (Thermo Fisher) with PageRuler Prestained protein marker (Thermo Fisher). After the run, the SDS-PAGE gel was transferred to nitrocellulose membranes for western blot analysis. The membranes were then processed further for western blot analysis, as described above. A detailed protocol is available (https://doi.org/10.17504/protocols.io.4r3l225xjl1y/v1).

To verify the activity of kinases Src and CK2, 45 μL of mixes containing either only kinase assay buffer (25 mM Tris-HCl pH 7.4, 150 mM NaCl, 1 mM DTT, and 2 mM MgCl_2_), kinase buffer and substrate (0.5 mg/mL) or kinase buffer, substrate (0.5 mg/mL) and kinase (100nM) were added to individual wells of a Pierce white opaque 96-well plate (Thermo Scientific). Substrate peptides used were RRRDDDSDDD 10-mer (PEP-CK2I-025, Biaffin) and Poly-(Glu,Tyr 4:1) (40217, BPS) for CK2 and Src kinases, respectively. For CK2, a specific inhibitor Silmitasertib CX-4945 (S2248, Selleckchem) was added, where indicated, at a concentration of 1 μM. Reactions were started by the addition of 5 μL ATP in kinase assay buffer, resulting in a final concentration of 100 μM ATP in each of the 50 μL reactions. After 1 h at room temperature (RT) in darkness, 50 μL of Kinase-Glo Max reagent (Promega) was added to each well, to reach a total volume of 100μL. The luciferase reactions were allowed to stabilize for 15 min before measuring luciferase activity at a Spark Multi-Mode Microplate Reader (TECAN). The luciferase activity correlates with ATP quantity, and thus, an inverse relationship between measured luminescence and kinase activity exists. A detailed protocol is available (https://doi.org/10.17504/protocols.io.5jyl82by7l2w/v1).

### Immunoprecipitation

HeLa cells were collected by trypsinization and the cell pellet was washed with PBS once before cells were lysed in lysis buffer (100 mM KCl, 2.5 mM MgCl_2_, 20 mM Tris-HCl pH 7.4, 0.5% NP-40). Samples were lysed for 20 min on ice before cell lysates were cleared by centrifugation at 20,000*g* for 10 min at 4°C. Protein concentrations of the cleared protein lysates were then determined with the Pierce Detergent Compatible Bradford Assay Kit (23246, Thermo Fisher) and equal amounts were incubated with beads. Beads were precoated with GST (negative control), NIX-GST, or BNIP3-GST as described above for the microscopy-based bead assay. HeLa cell lysates were incubated overnight with precoated beads. In the morning, samples were washed three times in lysis buffer before the beads were either submitted for analysis by mass spectrometry or for analysis by SDS-PAGE and western blotting by resuspending the beads in protein loading dye, supplemented with 100 mM DTT, and boiled for 5 min at 95°C. Samples were loaded on 4-12% SDS-PAGE gels (NP0322BOX, Thermo Fisher) with PageRuler Prestained protein marker (Thermo Fisher). Proteins were transferred onto nitrocellulose membranes (RPN132D, GE Healthcare) for 1 h at 4°C using the Mini Trans-Blot Cell (Bio-Rad). After the transfer, membranes were blocked with 5% milk powder dissolved in PBS-Tween (0.1% Tween 20) for 1 h at room temperature. The membranes were incubated overnight at 4°C with primary antibodies dissolved in the blocking buffer, washed three times for 5 min, and incubated with species-matched secondary horseradish peroxidase (HRP)-coupled antibodies diluted 1:10,000 in blocking buffer for 1 h at room temperature. Membranes were washed three times with PBS-T and processed further for western blot detection. Membranes were incubated with SuperSignal West Femto Maximum Sensitivity Substrate (34096, Thermo Fisher) and imaged with a ChemiDoc MP Imaging system (Bio-Rad). Images were analyzed with ImageJ ^72^ (RRID:SCR_003070; https://imagej.net/). A detailed protocol is available (https://doi.org/10.17504/protocols.io.kxygxynzwl8j/v1). The primary antibodies used in this study are: anti-GST (1:5000, Sigma-Aldrich Cat# SAB4200237, RRID:AB_2858197), anti-WIPI1 (1:200, Santa Cruz Biotechnology Cat# sc-376205, RRID:AB_10989262), anti-WIPI2 (1:500, Bio-Rad Cat# MCA5780GA, RRID:AB_10845951), anti-WIPI3 (Santa Cruz Biotechnology Cat# sc-514194, RRID:AB_3101990), anti-WIPI4 (Abcam Cat# ab168532, RRID:AB_3101989), anti-PPTC7 (1:500, Abcam Cat# ab122548, RRID:AB_11127117).

### Sample preparation for mass spectrometry analysis

After the final wash the beads were transferred to a new tube and resuspended in 30 µL 2 M urea in 50 mM ammonium bicarbonate and digested with 75 ng LysC (mass spectrometry grade, FUJIFILM Wako chemicals) and 75 ng trypsin (Trypsin Gold, Promega) at room temperature for 90 min. The supernatant was transferred to a new tube, the beads were washed with 30 µL 1 M urea and 50 mM ammonium bicarbonate, and the supernatant was pooled with the first eluate. Disulfide bonds were reduced with 10 mM dithiothreitol (DTT) for 30 min at room temperature before alkylation of free thiols with 20 mM iodoacetamide for 30 min at room temperature in the dark. The remaining iodoacetamide was quenched with 5 mM DTT for 10 min. The urea concentration was diluted to 1M with 50 mM ammonium bicarbonate. After addition of another 75 ng LysC and 75 ng trypsin, the digestion was continued at 37°C overnight. The digest was stopped by the addition of trifluoroacetic acid (TFA) to a final concentration of 0.5 %, and the peptides were desalted using C18 StageTips ^77,78^.

### Liquid chromatography Mass spectrometry analysis

Peptides were separated on a Vanquish Neo nano-flow chromatography system (Thermo-Fisher), using a trap-elute method for sample loading (Acclaim PepMap C18, 2 cm × 0.1 mm, 5 μm, Thermo-Fisher), and a C18 analytical column (Acclaim PepMap C18, 50 cm × 0.75 mm, 2 μm, Thermo-Fisher), applying a segmented linear gradient from 2% to 35% and finally 80% solvent B (80 % acetonitrile, 0.1 % formic acid; solvent A 0.1 % formic acid) at a flow rate of 230 nL/min over 120min. An Exploris 480 Orbitrap mass spectrometer (Thermo Fisher) coupled to the LC-column with a FAIMS pro ion-source (Thermo-Fisher) using coated emitter tips (PepSep, MSWil), was used with the following settings. The mass spectrometer was operated in DDA mode with two FAIMS compensation voltages (CV) set to -45 and -60 V and 1.5 s cycle time per CV. The survey scans were obtained in a mass range of 350-1500 m/z, at a resolution of 60k at 200 m/z and a normalized AGC target at 100%. The most intense ions were selected with an isolation width of 1.2 m/z, fragmented in the HCD cell at 28% collision energy and the spectra recorded for max. 50 ms at a normalized AGC target of 100% and a resolution of 15k. Peptides with a charge of +2 to +6 were included for fragmentation, the exclude isotope feature was enabled, and selected precursors were dynamically excluded from repeated sampling for 45 seconds.

### Mass spectrometry data analysis

Exploris raw files were first split according to CVs (-45 V, -60 V) using FreeStyle 1.7 software (Thermo Scientific). The resulting split MS data were analyzed with FragPipe (19.1 or 20.0), using MSFragger ^79^, IonQuant ^80^, and Philosopher ^81^. The default FragPipe workflow for label free quantification (LFQ-MBR) was used, except “Normalize intensity across runs” was turned off. Cleavage specificity was set to Trypsin/P, with two missed cleavages allowed. The protein FDR was set to 1%. A mass of 57.02146 (carbamidomethyl) was used as fixed cysteine modification; methionine oxidation and protein N-terminal acetylation were specified as variable modifications. MS2 spectra were searched against the Homo sapiens 1 protein per gene reference proteome from Uniprot (ID: UP000005640, release 2023_03), Spodoptera spp. sequences (UniProt taxonomy ID 7108, release 2023_03) and concatenated with a database of 382 common laboratory contaminants (release 2023.03, https://github.com/maxperutzlabs-ms/perutz-ms-contaminants) and two additional protein sequences corresponding to the expressed transgenic constructs. Computational analysis was performed using Python and the in-house developed library MsReport (versions 0.0.11 and 0.0.19 ^82^. Only non-contaminant proteins identified with a minimum of two peptides were considered for quantitative analysis. LFQ protein intensities reported by FragPipe were log2-transformed and normalized across samples using the ModeNormalizer from MsReport. This method involves calculating log2 protein ratios for all pairs of samples and determining normalization factors based on the modes of all ratio distributions. Missing values were imputed by drawing random values from a normal distribution. Sigma and mu of this distribution were calculated per sample from the standard deviation and median of the observed log2 protein intensities (μ = median sample LFQ intensity – 1.8 standard deviations of the sample LFQ intensities, σ = 0.3 × standard deviation of the sample LFQ intensities).

### Protein structure prediction with AlphaFold2-Multimer

Structures of biochemically identified protein complexes were predicted with AlphaFold-2 Multimer^83,84^. A locally installed version of AlphaFold-2 Multimer was used for structure prediction with 5 models per prediction followed by Amber relaxation. Interaction scores (ipDT) and diagnostic plots (PAE plot and pLDDT plot) as well as the generated structures were manually inspected. Predicted structures were visualized with ChimeraX-1.8^85,86^. A detailed protocol is available (https://doi.org/10.17504/protocols.io.81wgbz25qgpk/v1).

### AlphaFold 3 screen

We used AlphaFold^84^ to screen for putative WIPI2d interactors, by predicting interactions between WIPI2d and known selective autophagy receptors. We employed AlphaFold 3^87^ to run pairwise predictions with 5 models per prediction. Predictions with an ipTM score of > 0.5 were considered putative hits and diagnostic plots (PAE plot and pLDDT plot) as well as the generated structures were manually inspected. We also included FAM134C in our selection for experimental validation due to its ipTM score close to the 0.5 cut-off. The receptors included in the screen were: ATL3 (P82987), BCL2L13 (Q9BXK5), BNIP3 (Q12983), C53 (O94874), CALCOCO1 (Q9P1Z2), CCPG1 (Q9ULG6), FAM134A (Q8NC44), FAM134B (Q9H6L5), FAM134C (Q86VR2), FKBP8 (Q14318), FUNDC1 (Q8IVP5), MCL-1 (Q07820), NBR1 (Q14596), NDP52 (Q13137), NIX (O60238), NLRX1 (Q86UT6), NUFIP1 (Q9UHK0), OPTN (Q96CV9), PHB2 (Q99623), RTN3 (O95197), SEC62 (Q99442), SQSTM1/p62 (Q13501), TAX1BP1 (Q86VP1), TEX264 (Q9Y6I9), YIPF3 (Q9GZM5), YIPF4 (Q9BSR8). Soluble cargo receptors SQSTM1/p62, OPTN, NDP52, NBR1, and TAX1BP1 were predicted as dimers. Predicted structures were visualized with ChimeraX-1.8^85,86^. AlphaFold 3 predictions for FKBP8, TEX264, and FAM134C were validated with AlphaFold-2 Multimer accessed on the COSMIC^2^ server ^88^, resulting in similar predicted structures with exception of FAM134C. Settings for AlphaFold-2 Multimer were one prediction per model, full database, and relaxation of best model. A detailed protocol is available (https://doi.org/10.17504/protocols.io.6qpvr8rm2lmk/v1).

### Molecular dynamics simulations

We obtained the initial complex structure for the simulations from an AlphaFold-2.3 Multimer^83,84^ prediction using the full-length WIPI2d sequence and residues 30 to 82 from NIX.

We truncated the C-terminal IDR of WIPI2d and only used residues 1 to 362 for the simulations. We capped the N-terminus of the NIX fragment with an acetyl-group and the C-termini of both proteins with an aminomethyl-group. We used standard protonation states for a pH of 7.

We ran simulations of the wild-type and the LIR system, which we modelled by manually introducing the W36A and L39A mutations into the wild-type model. We used Gromacs (versions 2023.3 and 2023.4) ^89^ and the amber-disp force field ^90^ for all simulations. We solvated the proteins in water with 150 mM NaCl and neutralizing ions. We energy-minimized the system using the steepest descent algorithm with position restraints of 1000 kJ mol^-1^ nm^-2^ on all heavy atoms and a maximum force of convergence of 1000 kJ mol^-1^ nm^-1^. For equilibration, we performed one NVT and four NPT steps running for 1, 2, 1, 5, and 10 ns, respectively, and with a timestep of 1 fs for the first three steps and 2 fs for the last two steps. We gradually loosened the position restraints on heavy atoms during equilibration, using 1000 kJ mol^-1^ nm^-2^ in step 1 and 2, 500 kJ mol^-1^ nm^-2^ in step 3, 100 kJ mol^-1^ nm^-2^ in step 4, and no restraints in step 5. All equilibration steps and the production run used a v-rescale thermostat ^91^ with a target temperature T of 310 K and a characteristic time τ_T_ of 0.1 ps. The first NPT equilibration used a Berendsen barostat ^92^ with a target pressure p of 1 bar, a characteristic time τ_p_ of 5.0 ps, and a compressibility of 4.5 · 10^-5^ bar^-1^. All other NPT steps and the production run used a Parrinello-Rahman barostat ^93^ with p = 1 bar, τ_p_ = 5.0 ps, and a compressibility of 4.5 · 10^-5^ bar^-1^. Production runs used a timestep of 2 fs and were run for 1 μs. We performed triplicate simulations of both systems by initiating with different starting velocities.

In all simulations, we used a leap-frog integrator, a Verlet cutoff scheme ^94^, a cutoff of 1.0 nm modified with a potential shift for Van-der-Waals interactions, a cutoff of 1.0 nm for Coulomb interactions, and Particle Mesh Ewald for long-range electrostatics ^95^. We applied energy and pressure corrections for long-range Van-der-Waals interactions. We used the LINCS algorithm ^96^ to describe bonds with hydrogens.

We analyzed the behavior of the NIX LIR and its interaction with WIPI2d in these simulations by calculating three different quantities: the number of backbone hydrogen bonds n_h-bonds_ between NIX residues 35 to 39 and WIPI2d residues 129 to 134, the minimum distance d_TRP_ (d_Ala_ in the ΔLIR mutant) between any heavy atom of NIX W36 (A36 in the ΔLIR mutant) and the C_α_ atom of WIPI2d L119 (as a measure of W36/A36 insertion depth), and the minimum distance d_pocket_ between the sidechain heavy atoms of WIPI2d I133 and F169 (as a measure of pocket opening). We used trajectory frames every 1 ns for the analysis. For implementation of the described analysis we used Python3 (RRID:SCR_008394) with Anaconda3 (RRID:SCR_025572+), iPython ^97^, Numpy ^98^, Matplotlib ^99^, and MDAnalysis ^100^. We used VMD ^101^ and ChimeraX ^102^ for visual analysis and renders.

### Quantification and statistical analysis

For the quantification of immunoblots, we performed a densitometric analysis using Fiji software. Graphs were plotted using Graphpad Prism version 9.5.1 (RRID:SCR_002798). Depending on the number of samples, and as specified in the figure legends, we employed either a one-, or two-way ANOVA test with appropriate multiple comparison tests. Statistical significance is indicated with \**P*<0.05, \*\**P*<0.005, \*\*\**P*<0.001, \*\*\*\**P*<0.0001, ns, not significant. Error bars are reported as mean ± standard deviation. To ensure the reproducibility of experiments not quantified or subjected to statistical analysis, we showed one representative replicate in the paper of at least three replicates with similar outcomes for the main figures or at least two replicates for supplementary figures, as indicated in figure legends.

## Acknowledgments

We thank members of the Martens lab, Minghao Chen, Dorotea Fracchiolla, and other members of the Aligning Science Across Parkinson’s (ASAP) Mito911 Team for their help and advice. We thank Daniel Bernklau for optimization of the ATG9-vesicle purification protocol. We thank the Max Perutz Labs BioOptics, Flow Cytometry, and Mass Spectrometry facilities for their technical support. Proteomics analyses were performed by the Mass Spectrometry Facility at Max Perutz Labs using the VBCF instrument pool. We thank Ivana Bilusic Vilagos and the rest of the Vienna BioCenter Core Facilities (VBCF) Protech Facility for help with HEK cell expressions. The schematics were generated with BioRender. Molecular graphics and analyses performed with UCSF ChimeraX, developed by the Resource for Biocomputing, Visualization, and Informatics at the University of California, San Francisco, with support from National Institutes of Health R01-GM129325 and the Office of Cyber Infrastructure and Computational Biology, National Institute of Allergy and Infectious Diseases. This work was supported by a Marie Skłodowska-Curie MSCA Postdoctoral fellowship (101062916 to E.A.), a travel grant from the Flanders Fund for Scientific Research (FWO-Flanders to E.A.), and a Rebecca Cooper Foundation Fellowship (RC20241396 to M.L.). J.F.M.S. and G.H. thank the Max Planck Society and the Clusterproject ENABLE funded by the Hessian Ministry for Science and the Arts for financial support, and the Max Planck Computing and Data Facility for computational resources. This research was funded in whole or in part by Aligning Science Across Parkinson’s (ASAP-000350 to S.M., J.H.H., M.L., G.H.) through the Michael J. Fox Foundation for Parkinson’s Research (MJFF). For the purpose of open access, the authors have applied a CC-BY 4.0 public copyright license to all Author Accepted Manuscripts (AAM) arising from this submission.

## Author contributions

E.A., and S.M. conceived the project. E.A., S.S., A.S.I.C., J.S., J.S.M., and S.M. designed the experiments. E.A., S.S., A.S.I.C., J.S., J.S.M., T.N.N., X.R., M.S., J.R., and G.K. performed the experiments. J.F.M.S. carried out the MD simulations and part of the Alphafold predictions, supervised by G.H. E.A. and S.M. wrote the original draft to which all authors contributed by editing and reviewing.

## Data availability statement

Raw files associated with this work, including predicted AF structures, will be made available on Zenodo. The mass spectrometry proteomics data will be deposited to the ProteomeXchange Consortium via the PRIDE partner repository ^103^.

## Code availability statement

-

